# Protein mimetic amyloid inhibitor potently abrogates cancer-associated mutant p53 aggregation and restores tumor suppressor function

**DOI:** 10.1101/2020.08.10.243154

**Authors:** Loganathan Palanikumar, Laura Karpauskaite, Sarah Hassan, Maheen Alam, Mohamed Al-Sayegh, Ibrahim Chehade, Debabrata Maity, Liaqat Ali, Zackary Falls, Ram Samudrala, Mona Kalmouni, Yamanappa Hunashal, Jemil Ahmed, Shake Karapetyan, Renu Pasricha, Gennaro Esposito, Ahmed J. Afzal, Andrew D. Hamilton, Sunil Kumar, Mazin Magzoub

**Affiliations:** Biology Program, Division of Science, New York University Abu Dhabi, P.O. Box 129188, Saadiyat Island Campus, Abu Dhabi, United Arab Emirates; Department of Biology, SBA School of Science and Engineering, Lahore University of Management Sciences, Lahore, Pakistan; Department of Chemistry, New York University, New York, NY 10003, United States; Core Technology Platforms, New York University Abu Dhabi, P.O. Box 129188, Saadiyat Island Campus, Abu Dhabi, United Arab Emirates; Department of Biomedical Informatics, School of Medicine and Biomedical Sciences, State University of New York (SUNY), Buffalo, NY 14203, USA; Chemistry Program, Division of Science, New York University Abu Dhabi, P.O. Box 129188, Saadiyat Island Campus, Abu Dhabi, United Arab Emirates; INBB, Viale Medaglie d’Oro, 305, 00136 Roma, Italy; Department of Chemistry and Biochemistry and Knoebel Institute for Healthy Aging, The University of Denver, Denver, CO 80210, United States; Physics Program, Division of Science, New York University Abu Dhabi, P.O. Box 129188, Saadiyat Island Campus, Abu Dhabi, United Arab Emirates

**Keywords:** α-helix mimetics, amyloid, apoptosis, cancer therapeutics, cell cycle arrest, DNA-binding domain, oligopyridylamides, mutant p53, pancreatic cancer, protein aggregation, tumor suppressor

## Abstract

Missense mutations in p53 are severely deleterious and occur in over 50% of all human cancers. The vast majority of these mutations are located in the inherently unstable DNA-binding domain (DBD), many of which destabilize the domain further and expose its aggregation-prone hydrophobic core, prompting self-assembly of mutant p53 into inactive cytosolic amyloid-like aggregates. Screening an oligopyridylamide library, previously shown to inhibit amyloid formation associated with Alzheimer’s disease and type II diabetes, identified a tripyridylamide, ADH-6, that potently abrogates self-assembly of the aggregation-nucleating subdomain of mutant p53 DBD. Moreover, ADH-6 effectively targets and dissociates mutant p53 aggregates in human cancer cells, which restores p53’s transcriptional activity, leading to cell cycle arrest and apoptosis. Notably, ADH-6 treatment substantially shrinks xenografts harboring mutant p53 and prolongs survival, while exhibiting no toxicity to healthy tissue. This study demonstrates the first successful application of a bona fide small-molecule amyloid inhibitor as an anticancer agent.

## INTRODUCTION

Dubbed the ‘guardian of the genome’^1^, p53 is a tumor suppressor protein that is activated under cellular stresses, including DNA damage, oncogene activation, oxidative stress or hypoxia^2, 3^. Under normal conditions, p53 levels are kept low by its negative regulator, the E3 ubiquitin ligase MDM2, which targets p53 for proteasome-mediated degradation^2, 4^. The aforementioned cellular stresses disrupt the p53–MDM2 interaction, via phosphorylation of both proteins, and stimulate p53 acetylation, leading to its accumulation and activation^4^. Activated p53 then triggers DNA damage repair, cell cycle arrest, senescence, apoptosis or autophagy, all of which are directed towards the suppression of neoplastic transformation and inhibition of tumor progression^2, 3, 5^.

p53 binds to several DNA sequences, functioning as a sequence-specific transcriptional activator^3, 6^. p53 is also characterized by a high degree of structural flexibility, which facilitates its interactions with a myriad of protein partners, allowing it to exert its function as a master regulator of the cell^6^. Crucially, p53 missense mutations are found in over half of all human cancers, making it the most mutated protein in cancer, and these mutations are associated with some of the most pernicious manifestations of the disease^3, 7^. Thus, p53 has taken on a pivotal role in the realm of cancer research and is considered a key target in the development of cancer therapeutics^4, 7^.

Under physiological conditions, p53 exists as a homotetramer, with each monomer consisting of globular DNA-binding and tetramerization domains, connected by a flexible linker and flanked by intrinsically disordered regions (a transactivation domain followed by a proline-rich region at the N-terminus, and a C-terminal regulatory domain) (Figure 1a)^8^. The DBD consists of a central immunoglobulin-like β-sandwich serving as a scaffold for the DNA-binding surface, which is composed of a loop-sheet-helix motif and two large loops that are stabilized by the tetrahedral coordination of a single zinc ion (Figure 1b)^8^. A majority (*∼*90%) of cancer-associated p53 mutations occur within the DBD, where they cluster into discernible ‘hotspots’^9, 10^, resulting in the protein’s inactivation through alterations in residues that are crucial for either DNA interactions (contact mutants) or proper folding (structural mutants), although it is now apparent that some mutations (such as R248W) possess both characteristics^6^.

**Figure 1.**
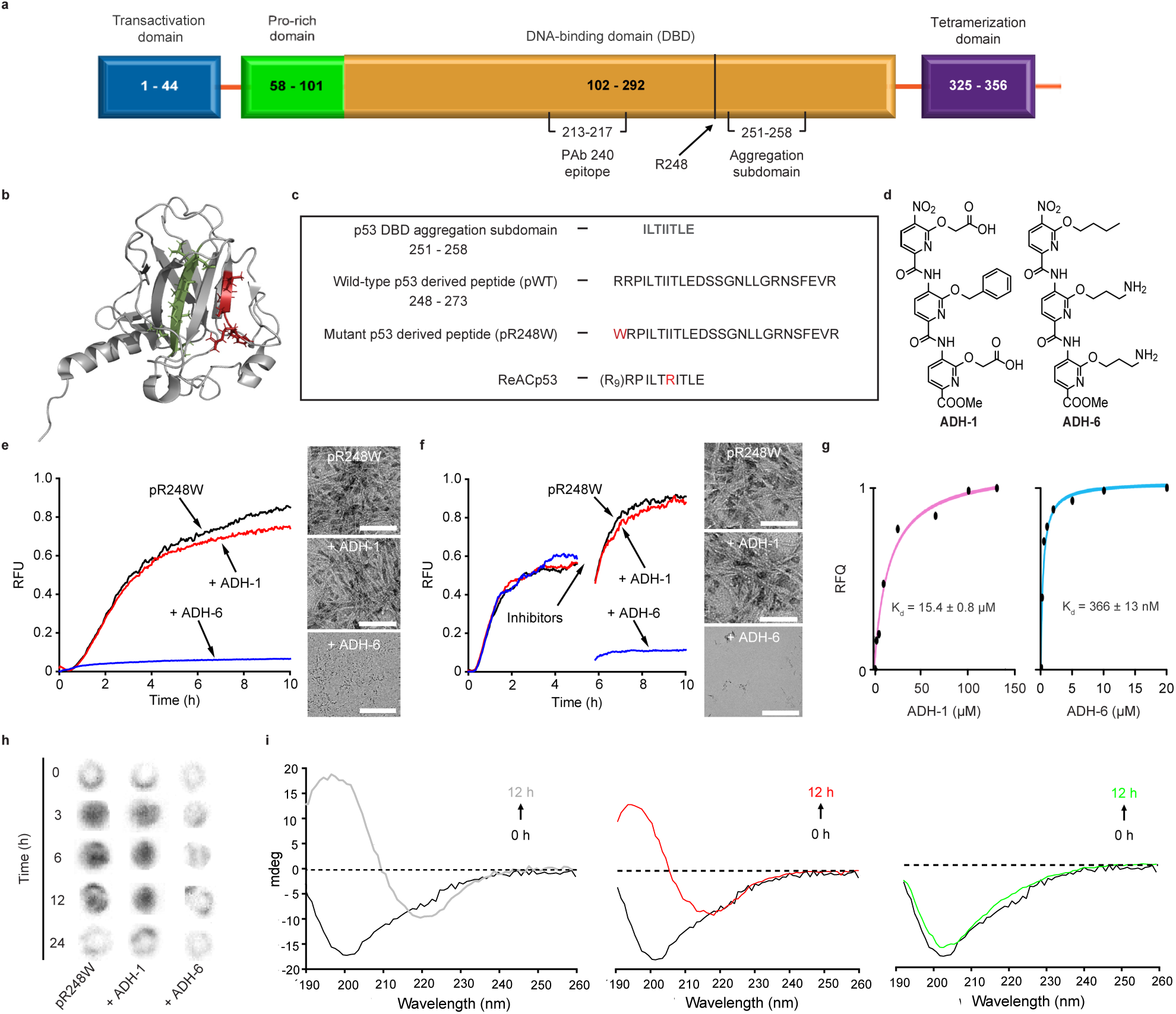
ADH-6 abrogates amyloid formation of aggregation-prone region of p53 DBD. (**a**) Schematic representation of the different domains of p53. The DBD (residues 102–292) contains an aggregation-nucleating subdomain (residues 251–258) that is necessary and sufficient to drive p53 aggregation^14, 17, 26^. Another segment of interest comprises residues 213–217, which is the antigen recognized by the PAb 240 antibody that binds to partially unfolded p53. Also highlighted in the DBD is R248, one of the most common mutation hotspots in p53 (IARC TP53 database; https://p53.iarc.fr)9. (**b**) Structure of p53 DBD. Highlighted are the aggregation-nucleating subdomain (green) and the epitope recognized by PAb 240 (red). Both segments are buried in the fully folded p53 structure. The 3D image was generated using PyMOL 2.3.5 (Schrödinger, New York, NY). (**c**) Primary sequences of the studied WT and mutant R248W p53 DBD-derived peptides, denoted pWT and pR248W, respectively, which span residues 248–273. The peptides include the aggregation-prone 252–258 sequence, as well as R248 and another of the most common mutation hotspots in p53, R273 (IARC TP53 database; https://p53.iarc.fr)9. (**d**) Chemical structures of the oligopyridylamides ADH-1 and ADH-6. (**e**,**f**) Effects of the oligopyridylamides on pR248W amyloid formation. Kinetic profiles (*left panel*) and representative transmission electron microscopy (TEM) images (*right panel*) for aggregation of 25 μM pR248W in the absence or presence of an equimolar amount of ADH-1 or ADH-6 co-mixed at the start of the reaction (**e**) or added during the growth phase (i.e. 5 h after the start of the reaction) (**f**). Kinetic aggregation profiles were acquired by measuring the fluorescence of the thioflavin T (ThT) reporter (λ_ex/em_ = 440/480 nm) at 5-min intervals at 37 °C (*n* = 4). TEM images were acquired at 10 h after the start of the aggregation reaction. Scale bar = 100 nm. (**g**) Characterization of the binding interaction of the oligopyridylamides and pR248W measured using steady-state intrinsic tryptophan fluorescence quenching. A 5 µM solution of pR248W was titrated with increasing concentrations of ADH-1 (*left panel*) or ADH-6 (right panel) and the tryptophan fluorescence after each addition was normalized to account for the dilution (total dilution during the titration was < 1%) and plotted against the ligand concentration. The equilibrium dissociation constants (*K*_d_) were then determined using a one-site specific binding equation (*Equation 1*). (**h**) Effects of the oligopyridylamides on pR248W oligomerization monitored using the dot blot assay. Samples of 10 μM Aβ42 were incubated with or without an equimolar amount of ADH-1 or ADH-6 for 0–24 h, and the presence of oligomers was detected using an amyloid oligomer-specific polyclonal antibody (A11)^33^. (**i**) Effects of the oligopyridylamides on the self-assembly driven structural transition of pR248W. Time-dependent circular dichroism (CD) spectra of 10 µM pR248W alone (*left panel*) or in the presence of an equimolar amount of ADH-1 (*middle panel*) or ADH-6 (*right panel*).

Several studies have reported that various p53 DBD mutants, along with fragments of these proteins, form amyloid-like aggregates in solution, cancer cells lines and tumors^11, 12^. p53 DBD is characterized by low thermodynamic and kinetic stability^8^, and mutations in the domain often decrease its stability further and prompt its unfolding, which leads to exposure of its hydrophobic core^13, 14^. Furthermore, many of the DBD mutations, including the commonly occurring R248W/Q, R273C/H and R175H^10^, involve replacing the cationic arginine, a so-called ‘gate-keeper’ amino acid that prevents protein aggregation using the repulsive effect of its charge^15^, with residues (tryptophan, cysteine, histidine, or glutamine) that have a high aggregation/amyloidogenic potential^15, 16^. Thus, these DBD mutations serve to not only expose the hydrophobic core of the domain, but also to enhance its aggregation propensity. This prompts self-assembly of mutant p53 into amyloid-like aggregates within inactive cellular inclusions that incorporate the wild-type (WT) isoform, thereby blocking the protein’s tumor suppressor function^17^.

Increasing evidence implicates aggregation of mutant p53 (e.g. R248Q, R248W, and R175H) in the associated oncogenic gain-of-function (GoF), i.e. the acquisition of activities that promote tumor growth, metastasis and chemoresistance^11, 12, 18^. For instance, co-sequestration of mutant and WT p53 into inactive cellular inclusions may result in overexpression of antiapoptotic and pro-proliferative genes previously repressed by p53^3, 5^. Aggregation of mutant p53 also induces misfolding of the p53 paralogs p63 and p73, which are then incorporated into the inclusions, facilitated by interactions of the aggregation-prone core of p53 DBD with near identical segments present in the p63 and p73 DBDs^17^. p63 and p73, which are rarely mutated in tumors, have partial functional overlap with p53^6^. However, co-aggregation with mutant p53 suppresses the regulatory functions of p63 and p73, resulting in deficient transcription of target genes involved in cell growth control and apoptosis, which leads to uncontrolled proliferation, invasion and metastasis^17^. Additionally, aggregation of mutant p53 has been shown to induce overexpression of heat-shock proteins, in particular Hsp70^17^, that promote tumor cell proliferation and inhibit apoptosis^19^. Thus, amyloid-like aggregation of mutant p53 may contribute to both its loss of tumor suppressor function and its oncogenic GoF.

Of relevance, replacing a hydrophobic amino acid in the aggregation-prone core of p53 DBD with the ‘gate-keeper’ arginine residue (I254R) abrogates coaggregation of mutant p53 with the WT protein and its paralogs, p63 and p73, as well as abolishes overexpression of Hsp70^17^. Notably, a p53 DBD-derived peptide harboring the aggregation-suppressing I254R mutation (denoted ReACp53; Figure 1c) was shown to block mutant p53 aggregation by masking the aggregation-prone core, which restored the mutant protein to a WT p53-like functionality and markedly reduced cancer cell proliferation *in vitro* and halted tumor progression *in vivo*^14^. These studies indicate that targeting mutant p53 aggregation is a viable and effective cancer therapeutic strategy.

We previously reported the use of oligopyridylamide-based α-helix mimetics to effectively modulate self-assembly of the amyloid-β peptide (Aβ)^20, 21^ and islet amyloid polypeptide (IAPP)^22^, which are associated with Alzheimer’s disease (AD) and type II diabetes (T2D), respectively. α-helix mimetics are small molecules that imitate the topography of the most commonly occurring protein secondary structure, serving as effective antagonists of protein-protein interactions (PPIs) at the interaction interface^23, 24^. The appeal of α-helix mimetics stems from the fact that their side-chain residues can be conveniently manipulated to target specific disease-related PPIs^23, 24^. In this study, we explored whether such a protein mimetic-based approach can be extended towards mutant p53 self-assembly. To that end, we asked the following questions: (i) if intrinsically disordered mutant p53 does indeed aggregate via an amyloid pathway, can the oligopyridylamide-based α-helix mimetics effectively abolish this process; (ii) if successful, does oligopyridylamide-mediated abrogation of mutant p53 aggregation lead to rescue of p53 function and inhibition of cancer cell proliferation *in vitro*; and (iii) can this oligopyridylamide-based strategy be applied to reverse tumor growth *in vivo* without adversely affecting healthy tissue? In addressing these questions, we establish the potential of using functionalized amyloid inhibitors as mutant p53-targeted cancer therapeutics.

## RESULTS

### ADH-6 abrogates amyloid formation of the aggregation-nucleating sequence of p53 DBD

We began with a reductionist approach commonly adopted in the amyloid research field, namely to target a short aggregation-prone segment within the protein of interest^25^. Across most amyloid systems, aggregation can be nucleated in a structure-specific manner by a small stretch of residues, which has a robust independent capacity for self-assembly^25^. Aggregation prediction algorithms developed from biophysical studies of amyloids identified an aggregation-nucleating subdomain (p53 residues 251–258) in the hydrophobic core of the p53 DBD that is necessary and sufficient to drive p53 aggregation^14, 17, 26^ (Figure 1a). This segment forms a *β*-strand within the hydrophobic core of the DBD (Figure 1b). However, many of the mutations in the p53 DBD destabilize its inherently unstable tertiary structure further^27^ and increase exposure of the aggregation-nucleating subdomain which, in turn, promotes aggregation of the protein^14, 17^. Consequently, we chose to focus on p53251–258 but expanded the sequence to include two of the most common mutation hotspots, R248 and R273 (IARC TP53 database; https://p53.iarc.fr)9. Two peptides were synthesized, one corresponding to residues 248–273 of WT p53 DBD (peptide denoted pWT), the other composed of the same sequence but harboring the R248W mutation (peptide denoted pR248W) (Figure 1c). R248W is one of the most common p53 DBD mutation that occurs in a range of malignancies, including pancreatic cancer^9, 10^. Pancreatic cancer is an intractable malignancy that often evades early diagnosis and resists treatment, and is consequently associated with a very poor prognosis: it is the seventh most common cause of death from cancer worldwide, with a five-year survival rate of < 5%^28, 29^.

We tested the effects of 10 compounds (ADH-1–10), based on the same oligopyridylamide molecular scaffold (Figure 1d and Supplementary Figure 1a), on the aggregation of pR248W using the thioflavin T (ThT)-based amyloid kinetic assay^30^. The compounds were selected based on their distinct chemical fingerprints and their ability to modulate amyloid assemblies^20–22^. The p53 DBD-derived peptides alone were characterized by a sigmoidal ThT curve, which is indicative of a nucleation-dependent process typical for amyloids (Supplementary Figure 1b)^31^. Comparing the two DBD-derived peptides, pR248W exhibited the greater aggregation propensity, as evidenced by the shorter lag phase, more rapid elongation phase and higher final ThT fluorescence intensity. This is not surprising given that the mutation involves replacing the cationic arginine, an aggregation ‘gate-keeper’ residue^15^ with the hydrophobic tryptophan, an aromatic residue with the highest amyloidogenic potential of all 20 naturally occurring amino acids^16^. Transmission electron microscopy (TEM) imaging confirmed that the aggregation-prone segment of mutant p53 DBD does indeed form fibrils (Figure 1e).

The 10 oligopyridylamides varied in their antagonist activity against pR248W aggregation. While the anionic ADH-1 did not significantly affect pR248W’s self-assembly, the cationic ADH-6 completely inhibited the peptide’s amyloid formation, as indicated by both the ThT assay and TEM imaging (Figure 1d,e). The determinants of efficacy of the oligopyridylamides appear to be the number and positioning of cationic sidechains (Figure 1d,e and Supplementary Figure 1a,b), suggesting that inhibition of pR248W aggregation occurs through specific interactions involving the compounds’ cationic sidechains. A possibility is that the oligopyridylamide–pR248W binding is stabilized by cation–*π* interactions of a cationic sidechain of the protein mimetic and the aromatic tryptophan residue of the mutant peptide^32^, which provides a basis for strong binding and confers a degree of specificity to the interaction. This is strongly supported by the substantially higher binding affinity of ADH-6 for pR248W (Kd = 366 ± 13 nM) compared to pWT (Kd = 15.4 ± 0.8 µM) (Figure 1g). This suggests that similar cation–*π* interactions between the cationic sidechains of ADH-6 and one or more of the several aromatic residues (tyrosines and tryptophans) present in the DBD^9^ may facilitate strong binding of the oligopyridylamide to full length mutant p53.

In order to restore WT p53-like activity in cancer cells harboring aggregation-prone mutant p53, potential therapeutics would need to dissociate pre-formed mutant p53 aggregates, as well as prevent additional aggregation. Therefore, we tested the capacity of ADH-6 to abrogate pre-formed pR248W aggregates. Addition of ADH-6 to the mutant peptide at the mid-point of the aggregation reaction, when a significant amount of fiber formation had already taken place (Figure 1f and Supplementary Figure 1c), resulted in a marked decrease in the ThT fluorescence intensity and near complete absence of fibers in the TEM images (Figure 1f) at the end of the reaction. The very few fibers that were detected were much shorter and thinner than those observed for the peptide alone (Figure 1f and Supplementary Figure 1d).

To further confirm the capacity of ADH-6 to prevent aggregation of pR248W, we evaluated the effect of the oligopyridylamide on the peptide’s oligomerization using a dot blot immunoassay (Figure 1h). pR248W was incubated with or without an equimolar amount of ADH-6 or ADH-1 for 0–12 h and then detected using A11, an antibody specific for amyloid oligomers, including those of mutant p53^33, 34^ (Figure 1h). The chemiluminescence signal intensity for samples of pR248W alone increased from 0 to 6 h, indicating an increase in the amount of soluble oligomers. Subsequently, the intensity diminished significantly at 12 h due to conversion of the oligomers into fibrils. Treatment with ADH-1 did not significantly change the intensity of the dots, indicating that the oligopyridylamide did not affect oligomerization of pR248W. In marked contrast, in the presence of ADH-6 no formation of pR248W oligomers was observed, as reflected by the weak intensity of the dots throughout the time course of the experiment. Finally, we probed the effect of ADH-6 on the secondary structure of pR248W using CD spectroscopy (Figure 1i). The peptide transitioned from a random coil monomer to a β-sheet structure as it self-assembled into amyloid fibers. pR248W underwent the same conformational transition in the presence of an equimolar amount of ADH-1, which was not surprising given its inability to inhibit the mutant peptide’s oligomerization or amyloid formation (Figure 1e,f,h). On the other hand, in the presence of ADH-6 at an equimolar ratio, pR248W remained in its native conformation for the duration of the experiment, which confirms that ADH-6 potently inhibits self-assembly of pR248W (Figure 1e,f,h). Taken together, these results demonstrate that ADH-6 not only strongly inhibits self-assembly of the aggregation-prone segment of mutant p53 DBD, the oligopyridylamide also effectively dissociates pre-formed aggregates of the segment and prevents further aggregation.

### Computational modeling of ADH-6 binding to WT and mutant p53 DBDs

To evaluate the relative p53 DBD–oligopyridylamide interaction strength, CANDOCK (Chemical Atomic Network-Based Hierarchical Flexible Docking) algorithm^35^ was used to model the binding between the two protein structures (WT and mutant R248W p53 DBDs) and the oligopyridylamides (ADH-1 and ADH-6), as well as two peptide sequences (LTIITLE and LTRITLE) (Figure 2).

**Figure 2.**
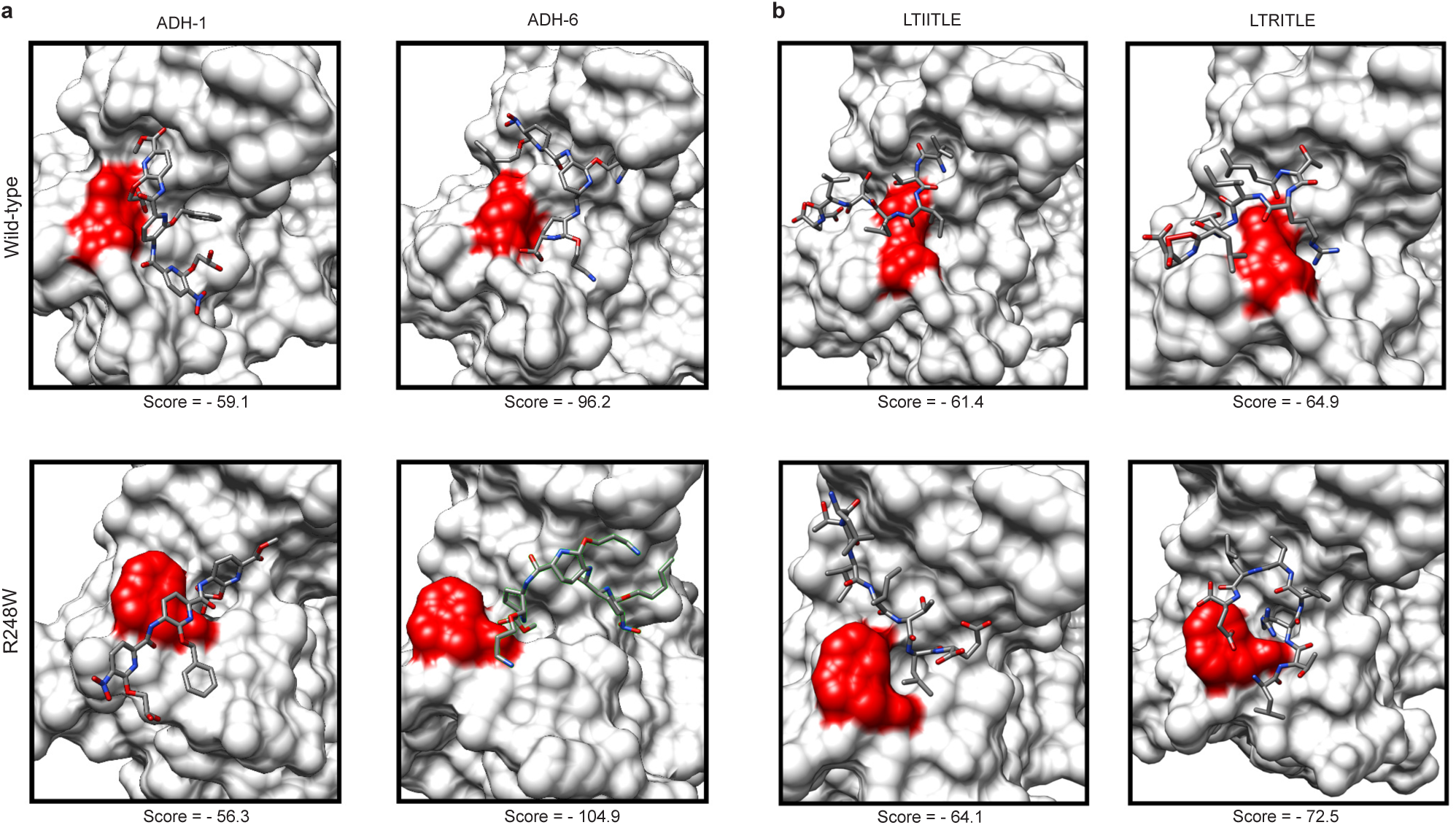
Computational modeling of p53 DBD–oligopyridylamides interaction strength. (**a**,**b**) Top binding modes and the resulting scores for ADH-1 and ADH-6 (**a**) and LTIITLE and LTRITLE (**b**) bound to the modeled structures of p53 (WT and mutant R248W) DBDs. For each binding mode, the DBD is depicted as a white surface, the residue of interest (248) is highlighted in red, and the corresponding compounds and peptides colored by element. The scores are based upon the CANDOCK knowledge-based scoring potential, where lower scores correspond to a more favorable binding.

We first modeled the pose and strength of binding between the hydrophobic core of the DBD variants and the two oligopyridylamides (Figure 2a). ADH-6 docked very strongly to both WT and R248W p53 DBDs (−96.2 and −104.9, respectively) compared to ADH-1 (−59.1 and −56.3 scores for WT and R248W p53 DBDs, respectively). These results show that the interaction of ADH-6 with p53 DBD variants is energetically favored over that of ADH-1. As a control, we tested ADH-7, a potent Aβ inhibitor^20, 21^ that also effectively antagonized pR248W aggregation (Supplementary Figure 1a,b). As expected, ADH-7 docked much more strongly (−101.4 score for both WT and R248W p53 DBDs) compared to ADH-1 (Supplementary Figure 2). Interestingly, ADH-7’s interaction with the mutant DBD was somewhat weaker than that of ADH-6, which correlates with their relative efficacy in inhibiting pR248W aggregation (Figure 1e,f and Supplementary Figure 1a,b). Thus, the docking simulations corroborate the ThT-based fibrillation assay results.

In addition, we determined whether the predicted binding scores for the oligopyridylamides were relatively stronger or weaker than those for known sequences that bind to either cause or inhibit mutant p53 amyloid aggregation. The chosen sequences were: (i) LTIITLE, the aggregation-nucleating sequence of p53 DBD identified by the Eisenberg group (corresponding to p53252–258)^14^; and (ii) LTRITLE, the same sequence but bearing the aggregation-suppressing I254R mutation^14, 17^. The LTIITLE and LTRITLE peptides were docked and the resulting top binding poses are shown in Figure 2b. The predicted scores indicate that LTIITLE binds less favorably than the LTRITLE mutant to both WT (−61.4 and −64.9, respectively) and mutant R248W (−64.1 and −72.5, respectively) p53 DBDs, which is in agreement with the reported ability of the LTRITLE-containing ReACp53 peptide to inhibit mutant p53 amyloid self-assembly^14^. Furthermore, the scores for binding of LTIITLE and LTRITLE to WT and mutant p53 DBDs fall in between the binding scores for ADH-1 and ADH-6 (Figure 2a). These binding scores concur with the results of the pR248W aggregation assays (Figure 1e,f,h), and strongly suggest that ADH-6 would be more effective than either ADH-1 or ReACp53 at antagonizing intracellular mutant p53 aggregation.

### Nuclear magnetic resonance (NMR) spectroscopy characterization of p53 DBD– ADH6 interaction interface

NMR spectroscopy was used to elucidate the interactions between ADH-6 and WT and mutant p53 DBD. Figure 3 shows the overlay of the HSQC maps, with and without ADH-6 addition, for WT (Figure 3a) and mutant R248W (Figure 3b) p53 DBDs. Chemical shift perturbation (CSP) analysis was carried out to determine the protein-ligand interaction interface. The interaction with ADH-6 also involved partially unfolded species that were present in the samples of both DBD variants (Figure 3a,b). Due to this additional involvement, no quantitative estimate was reliably feasible to assess a binding constant of ADH-6 to WT and R248W p53 DBDs. A qualitatively equivalent pattern was observed with either DBD variant (see Supplementary Section 2). The increment of the CSP values with ligand concentration (Figure 3c) is the signature of a fast exchange regime, i.e. no stable complex formation, without any major favorable or adverse consequence of the hotspot mutation^36^ on the ADH-6 affinity. This result does not necessarily conflict with the nanomolar affinity of ADH-6 for pR248W inferred from the fluorescence quenching experiments (Figure 1g), nor the strong interaction between the oligopyridylamide and the hydrophobic core of mutant p53 DBD determined by the docking simulations (Figure 2), because p53 DBD presents numerous interaction sites to ADH-6 (*vide infra*) beyond the aggregation-nucleating subdomain considered in the fluorescence quenching experiments or docking simulations. However, it is also possible that the fast exchange regime observed by NMR may be, in part, a consequence of the experimental conditions used to enable study of the inherently unstable p53 DBDs^36, 37^ (see Supplementary Section 2).

**Figure 3.**
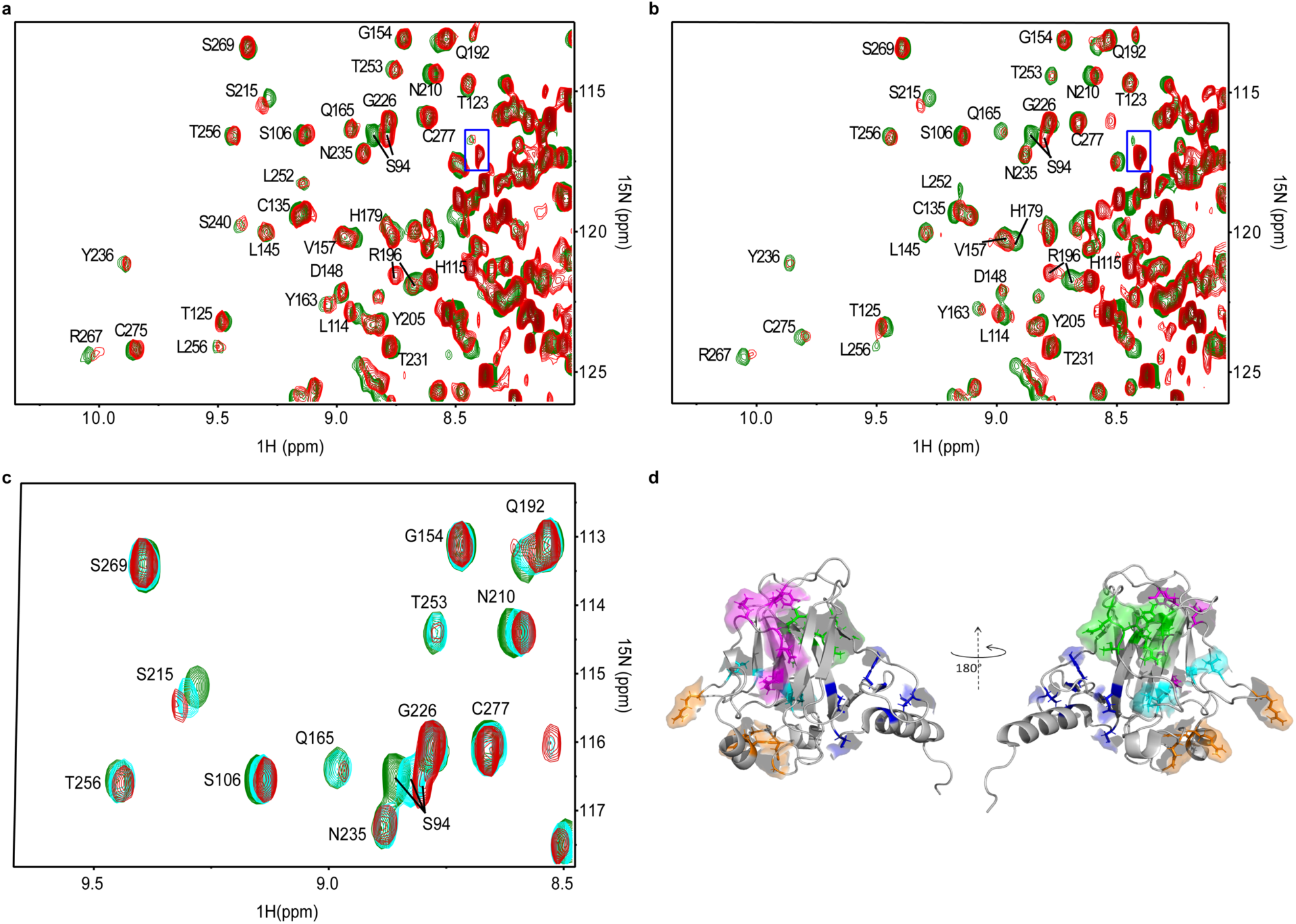
NMR-based determination of p53 DBD–ADH-6 interaction interface. (**a**,**b**) Overlay of ^15^N-^1^H HSQC maps of 19 μM WT (**a**) and 24 μM R248W (**b**) p53 DBD in H_2_O/D_2_O (96/4) with 16.7 mM DTT, without (green contours) or with (red contours) ADH-6 addition (protein:ligand 1:11 in **(a)** and 1:15 in (**b**)). The assignments are reported only outside the rightmost regions. These regions are crowded because of the presence of a partially unfolded species that also interact with ADH-6 as highlighted by the boxed peak in each panel. (**c**) HSQC contour maps overlay of mutant R248W p53 DBD at different protein:ADH-6 ratios (1:0 green, 1:8 cyan and 1:15 red) showing the increment of cumulated chemical shift perturbation (CSP) with ligand concentration (*Equation 2*). (**d**) The five clusters of the two p53 DBD variants (WT and mutant R248W) that show high (> 0.025) or medium (> 0.015) CSP values^99^. Cluster 1 (highlighted in blue) includes residues T118, Y126, E271, C275 and G279; cluster 2 (highlighted in magenta) includes residues R196, E198, G199, L201, Y220 and E221; cluster 3 (green) includes T102, Y103, Q104, G105, L257, L264 and R267; cluster 4 (orange) includes E171, R174, H179, R209 and G244; and cluster 5 (cyan) includes S94, A161, I162, L206 and S215. Clusters 1 and 2 are at the front in the cartoon on the left; clusters 3, 4 and 5 are at the front in the cartoon on the right. The 3D image was generated using PyMOL 2.3.5 (Schrödinger, New York, NY).

The backbone NH signals that exhibit the most significant CSP values in both the considered p53 DBD variants appeared to involve five clusters of residues that are highlighted with different colors in Figure 3d. The first cluster (highlighted in blue) encompasses the area around the start of the C-terminal helical fragment of the p53 DBD (helix 3, PDB:*2FEJ*^37^). The second cluster (magenta) extends along strand 4 of sheet A and the subsequent apical loop. The third cluster (green) is formed by a sequential stretch of the apical loop preceding strand 1 of sheet A along with the initial and final facing fragments of strands 3 and 4 in sheet B, respectively. The fourth cluster (orange) covers an extended surface surrounding the start of helix 2. And the fifth, and final, cluster (cyan) is assembled with contributions from very distant residues, i.e. N-terminal and start and end of strands 6 and 7, respectively, of sheet B. Interestingly, cluster 1 includes E271 and cluster 3 includes L257, L264 and R267, i.e. residues of the p53248–273 segment we selected for our DBD-derived peptides, pWT and pR248W (Figure 1), which has a high frequency of mutations affecting p53 function^14, 36, 37^. In particular, L257 of cluster 3 belongs to the aggregation-nucleating subdomain (p53251–258) that promotes amyloid-like aggregation of mutant p53^14, 36^.

The conspicuous extension of the protein surface region that is affected by ADH-6 interaction, based on the NMR evidence, should not suggest the occurrence of a rather generic interaction altogether. Approximately 75% of residues involved in the clusters of Figure 3d are predicted to be aggregation prone^14^, which provides a basis for the effect of ADH-6 on sensitive locations other than p53251–258 in p53 DBD. Chemical shift changes, however, may also be the consequence of local rearrangements in response to an allosteric interaction. Therefore, only part of the highlighted surface regions of Figure 3d may actually be directly involved in the dynamic contacts with ADH-6. A possible clue for the determination of these contacts may come from the inspection of the relative peak intensities (RI), i.e. the ratio of the peak heights in the presence and absence of ADH-6 (Supplementary Section 2). Dynamic interactions may in fact induce intensity attenuations because of exchange and/or dipolar broadening, thereby providing a further interpretation tool. Unambiguous attenuations of the intensities leading to deviations by more than one standard deviation from the average RI value were observed for K120, T123, Q192 and R196 in the WT DBD, and for Q192 and R196 in the mutant species. The first pair of highly attenuated NHs in the WT variant suggests a contact at cluster 1 (blue; Figure 3d), close to the C-terminal helix, which is consistent with the designed properties of ADH-6. The attenuation of Q192 and R196 amide peaks points to a contact occurring at cluster 2 in both species (magenta; Figure 3d).

Besides attenuations, intensity increments were also observed upon ADH-6 addition, suggesting an increase in local mobility at residues of the C-terminal region (E298 and R306) and E271 (cluster 1). However, it was only with mutant R248W p53 DBD that intensity increments were also sampled at L257 and L264 (cluster 3), thereby highlighting a hotspot mutation effect^14, 36^. In particular, for the mutant species, the mobility increase associated with ADH-6 contact at cluster 2 (at least) seems to involve the adjacent extremities of strands 3 (L264) and 4 (L257) of sheet B, as well as at the opposite extremity of strand 3 (E271), in addition to some points of the C-terminal region. Conversely, for the WT DBD, the mobility increments accompanying ADH-6 contact affect only the residues of the C-terminal region.

### ADH-6 dissociates intracellular mutant p53 aggregates

Given the potent antagonism of self-assembly of the aggregation-prone segment of mutant p53 DBD (Figure 1e,f,h), the high affinity for that segment and full-length mutant p53 DBD (Figures 1h and 3), as well as the multiple binding sites on the DBD (Figure 2), we probed the effects of ADH-6 on intracellular mutant p53 aggregates (Figure 4). MIA PaCa-2 cells harboring aggregation-prone mutant R248W p53 were stained with thioflavin S (ThS), which is commonly used as a marker for intracellular amyloid aggregates, including those of mutant p53 tissue^34, 38^ (Figure 4a,b). For some experiments, MIA PaCa-2 cells were co-stained with ThS and the anti-p53 antibody PAb 240 (Figure 4c–f). PAb 240 is specific for partially unfolded p53 as it recognizes an epitope (residues 213–217; Figure 1a,b) that is buried in folded p53. Since partially unfolded p53 is required for protein aggregation, PAb 240 is often used as a marker for aggregated p53^14^.

**Figure 4.**
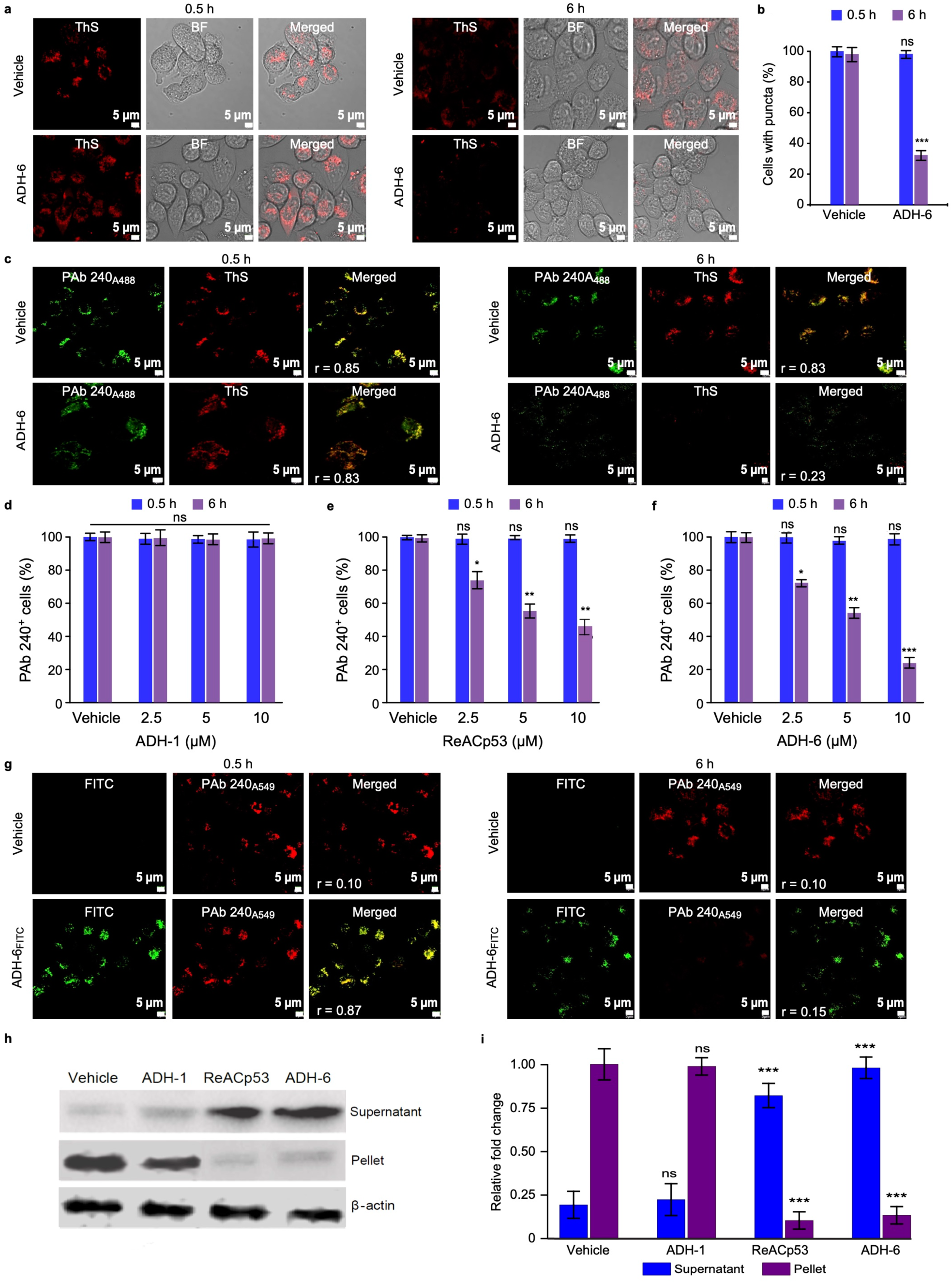
ADH-6 dissociates mutant p53 aggregates in cancer cells. (**a**) Confocal fluorescence microscopy images showing thioflavin S (ThS) staining of mutant p53 (R248W) aggregates in MIA PaCa-2 cells treated with vehicle (0.02% DMSO) or ADH-6 (5 µM) for 0.5 or 6 h. **(b)** Quantification of ThS-positive MIA PaCa-2 cells after treatment with vehicle or ADH-6. (**c**) Confocal fluorescence microscopy images of ThS and PAb 240 antibody staining of R248W aggregates in MIA PaCa-2 treated with vehicle or 5 µM ADH-6 for 0.5 or 6 h. (**d–f**) Quantification of PAb 240-positive MIA PaCa-2 cells after treatment with the indicated concentrations of ADH-1, ReACp53 or ADH-6 for 0.5 or 6 h relative to controls (vehicle-treated cells). (**g**) Colocalization of FITC-labeled ADH-6 (ADH-6FITC) with PAb 240-stained R248W aggregates following incubation with the oligopyridylamide (5 µM) for 0.5 or 6 h. Imaging experiments were performed in triplicate and representative images are shown. Colocalization was quantified using Pearson’s correlation coefficient, r, which measures pixel-by-pixel covariance in the signal level of two images^129^. Scale bar = 5 µm. For quantification, the number of positively stained cells in 3–5 different fields of view are expressed as % of the total number of cells (*n* = 3). (**h,i**) Western blot analysis of the effects of ADH-6 on mutant p53 aggregation. Immunoblots of mutant R248W p53 in the soluble (supernatant) and insoluble (pellet) fractions of MIA PaCa-2 cells treated with vehicle or 5 µM ADH-1, ReACp53 or ADH-6 for 24 h and detected by the anti-p53 antibody DO-7 (**h**). Densitometric quantification of the immunoblot bands of mutant p53 in the soluble and insoluble fractions (*n* = 3) (**i**). \**P* < 0.05, \*\**P* < 0.01, \*\*\**P* < 0.001 or non-significant (ns, *P* > 0.05) for comparisons with controls.

Punctate cytosolic ThS staining was observed in MIA PaCa-2 cells, indicating that mutant R248W p53 self-assembles into amyloid aggregates in the cytosol (Figure 4a,c). Treatment with ADH-6 for 6 h led to a substantial reduction in the number of ThS-positive puncta, with the proportion of cells containing these puncta decreasing to *∼*30% (Figure 4a,b). Strong colocalization of the ThS and PAb 240 signals was observed in the vehicle-treated control cells (Figure 4c), confirming that ThS stains cytosolic mutant p53 aggregates. Treatment of dual-stained MIA PaCa-2 cells with ADH-6 markedly reduced both the ThS and PAb 240 signals (Figure 4c). The effect of ADH-6 on mutant p53 aggregates was dose-dependent, with the PAb 240-positive cells decreasing to *∼*50 and 24% of controls at 5 and 10 µM ADH-6, respectively (Figure 4f). Treatment with ReACp53, which has been reported to disaggregate mutant p53 in clinical samples and stable cells established from patients^14^, also significantly reduced the proportion of ThS- and PAb 240-positive cells (PAb 240-positive cells were *∼*40% of controls at 10 µM ReACp53) (Figure 4e and Supplementary Figure 8). In contrast, ADH-1, which did not antagonize pR248W aggregation (Figure 1e,f), had a negligible effect on the ThS or PAb 240 staining (Figure 4d and Supplementary Figure 8).

As a control, we tested the effects of ADH-1, ReACp53 and ADH-6 on mutant p53 DBD aggregates in plant cells. p53 is not found in plants, which are usually (i.e. in the absence of pathogens) not susceptible to neoplasia^39^. Yet, remarkably, stable transfection of p53 in the model plant *Arabidopsis* was shown to induce early senescence^40^. Thus, plants represent a novel system for studying mutant p53 DBD aggregation. Here, we amplified the genes using primers corresponding to p53 DBD and cloned them in a binary vector expressing an N-terminal yellow fluorescence protein (YFP) tag, under the control of cauliflower mosaic virus (CaMV35S) promoter. The cloned constructs, YFP-tagged WT and mutant R248W p53 DBDs (YFP:p53DBD^WT^ and YFP:p53DBD^R248W^, respectively), were subsequently expressed in *Nicotiana benthamiana* using *Agrobacterium*-mediated transient infiltrations. While the WT DBD accumulated exclusively in the nucleus, expression of R248W p53 DBD resulted in a number of foci spread throughout the cell (Supplementary Figure 9a), indicating that the mutant protein forms cytosolic aggregates in plant cells similar to those detected in mammalian cells (Figure 4). Treatment of the *N. benthamiana* leaves with ADH-6 or ReACp53 resulted in significantly smaller puncta in the cells expressing the mutant p53 DBD (Supplementary Figure 9a,c). Our results are consistent with reports that ReACp53 disaggregates cytosolic mutant p53 puncta^14, 41^, which leads to accumulation of the released protein in the nucleus^14^. On the other hand, ADH-1 did not reduce the size of WT or mutant R248W DBD puncta (Supplementary Figure 9a–c).

To ascertain whether ADH-6 mediated dissociation of mutant p53 aggregates is a result of direct interaction with the oligopyridylamide, PAb 240-stained MIA PaCa-2 cells were treated with FITC-labeled ADH-6 (ADH-6FITC) (Figure 4g). Initially, strong colocalization of ADH-6FITC and PAb 240 was observed, indicating direct interaction of the oligopyridylamide with mutant p53 aggregates. Eventually, the extent of colocalization decreased as the PAb 240 staining was reduced due to disaggregation of mutant p53. These results indicate that, similar to ReACp53^14^, ADH-6 efficiently enters cells to directly interact with and dissociate cytosolic mutant p53 aggregates.

Finally, to corroborate the imaging results, MIA PaCa-2 cells were treated with ADH-1, ReACp53 or ADH-6, lysed and fractionated, and the amount of mutant p53 in the soluble and insoluble fractions was quantified by western blot (Figure 4h,i). As expected, treatment with ADH-1 did not alter the distribution of mutant p53 relative to the vehicle-treated controls, with the aggregation-prone protein strongly detected in the insoluble fraction, but almost completely absent from the soluble fraction. However, treatment with ADH-6 or ReACp53 significantly decreased the mutant p53 content of the insoluble fraction, while markedly increasing the amount of protein in the soluble fraction. These results confirm that ADH-6 targets and converts insoluble cytosolic mutant p53 aggregates into soluble protein.

### ADH-6 causes selective cytotoxicity in cancer cells bearing mutant p53

Next, we determined the effects of the oligopyridylamides on viability of mutant p53-harboring cancer cells using the MTS assay. A screen of the compounds revealed a strong correlation between their toxicity in mutant R248W p53-bearing MIA PaCa-2 cells (Figure 5a and Supplementary Figure 10a,b) and their capacity to antagonize pR24W aggregation (Figure 1e,f and Supplementary Figure 1b). Indeed, ADH-6, which was the most potent antagonist of pR24W amyloid formation, was also the most toxic compound to MIA PaCa-2 cells. ADH-6 reduced MIA PaCa-2 cell viability in a concentration-dependent manner, with an effective concentration (EC_50_) of 2.7 ± 0.4 and 2.5 ± 0.1 µM at 24 and 48 h incubation times, respectively, which was almost half of that for ReACp53 (EC_50_ = 5.2 ± 0.5 and 4.9 ± 0.3 µM at 24 and 48 h incubation, respectively) (Supplementary Figure 10d). On the other hand, ADH-1, which did not inhibit pR248W aggregation, had no adverse effect on viability of MIA PaCa-2 cells (Figure 5a). Importantly, the oligopyridylamides, including ADH-6, were completely nontoxic to WT p53-bearing breast cancer MCF-7 cells (Figure 5b and Supplementary Figure 10c). Moreover, testing the effects of ADH-6 on a range of cancer cell lines, we observed no significant toxicity in those bearing WT p53 (A549, U-87 MG, SW-480 and SW-1990) (Supplementary Figure 10e), but a substantial *∼*75–85% decrease in viability of the ones harboring aggregation-prone R248W/Q mutant p53 (NCI-H1770, HCC70, COLO 320DM and OVCAR-3) at the same oligopyridylamide concentration and incubation time (Supplementary Figure 10f). Notably, ADH-6 did not adversely affect viability of p53 null SKOV-3 cells; however, transfecting the cells with mutant R248W p53 (Supplementary Figure 10g) rendered them susceptible to ADH-6 mediated cytotoxicity (EC_50_ = and 9.9 ± 1.5 and 7.5 ± 1.1 µM at 24 and 48 h incubation, respectively) (Figure 5c,d). These results strongly suggest that the observed cytotoxicity of ADH-6 in cancer cells is directly related to the oligopyridylamide’s capacity to antagonize mutant p53 amyloid formation.

**Figure 5.**
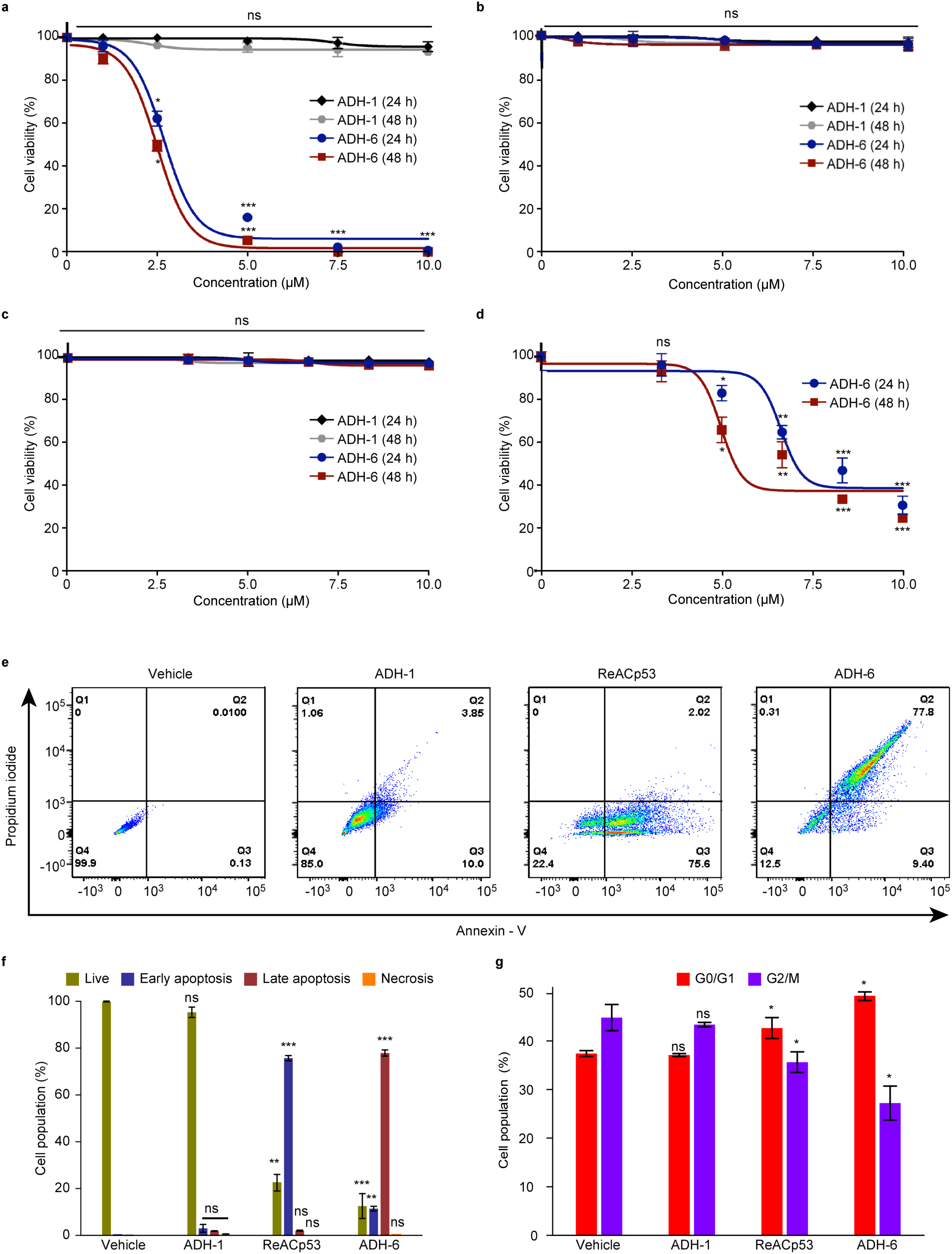
ADH-6 causes death of cancer cells bearing mutant, but not WT, p53. (**a–d**) Effects of ADH-1 and ADH-6 on viability of cancer cells bearing WT or mutant p53. (**a**,**b**) Mutant R248W p53-harboring MIA PaCa-2 (**a**) or WT p53-harboring MCF-7 (**b**) cells treated with increasing concentrations of ADH-6 or ADH-1 for 24 or 48 h. (**c**,**d**) p53 null SKOV-3 before (**c**) and after transfection with R248W p53 (**d**) treated with increasing concentrations of ADH-6 or ADH-1 for 24 or 48 h. Cell viability in (**a**–**d**) was assessed using the MTS assay, with the % viability determined form the ratio of the absorbance of the treated cells to the control cells (*n* = 4). (**e**,**f**) Flow cytometry analysis of annexin V/propidium iodide (PI) staining of MIA PaCa-2 cells that were treated with vehicle (control), or 5 µM ADH-1, ReACp53 or ADH-6, for 24 h (**e**). The bottom left quadrant (annexin V-/PI-) represents live cells; bottom right (annexin V+/PI-), early apoptotic cells; top right (annexin V+/PI+), late apoptotic cells; and top left (annexin V-/PI+), necrotic cells. A summary of the incidence of early/late apoptosis and necrosis in the different treatment groups determined from the flow cytometry analysis of annexin V/PI staining (*n* = 4) (**f**). (**g**) Cell cycle distribution of MIA PaCa-2 cells treated with vehicle (control), or 5 µM ADH-1, ReACp53 or ADH-6, for 6 h as determined by measurement of DNA content using flow cytometry (*n* = 4). \**P* < 0.05, \*\**P* < 0.01, \*\*\**P* < 0.001 or non-significant (ns, *P* > 0.05) compared with controls.

An indicator of restored p53 function is activation of apoptosis. Therefore, following treatment with vehicle, or 5 µM ADH-1, ReACp53 or ADH-6, MIA PaCa-2 cells were stained with Alexa 488-conjugated annexin V/propidium iodide (PI) and quantified by flow cytometry, a common method for detecting apoptotic cells^42^ (Figure 5e,f). As expected, treatment with ADH-1 had a negligible effect. Exposure to ReACp53 resulted in 75.7 ± 1.1 and 2.0 ± 0.2% of cells undergoing early and late apoptosis, respectively, which is in line with the reported capacity of the peptide to induce apoptosis in aggregation-prone mutant p53-bearing cancer cells^14^. However, treatment with ADH-6 led to an even more pronounced effect, with 11.4 ± 0.9 and 77.9 ± 1.3% of cells undergoing early and late apoptosis, respectively. The rescue of p53 activity was further corroborated by cell cycle analysis (Figure 5g and Supplementary Figure 10h). In contrast to ADH-1, treatment with ReACp53 resulted in a small but significant shift in the cell cycle distribution of the asynchronous population, which is a hallmark of p53 activation^14^. However, yet again we observed an even greater effect following exposure to ADH-6, with a larger shift in the cell cycle distribution occurring (i.e. more cells in the G0/G1 phase and fewer in G2/M) relative to ReACp53. Collectively, our results demonstrate that ADH-6 mediated cytotoxicity in cancer cells is due to abrogation of mutant p53 aggregation by the oligopyridylamide, which leads to restoration of WT p53 function.

### ADH-6 induces transcriptional reactivation of p53

To confirm that ADH-6 rescues normal p53 function in MIA PaCa-2 cells, transcriptome analysis was performed to assess the effects of oligopyridylamide on the mutant p53-bearing cells at the global gene level (Figure 6). Total RNA samples were isolated from ADH-6, ADH-1 and vehicle (control, C) treated cells, which was followed by RNA-Seq library preparation. To establish the best condition for differential gene expression analysis, we applied correlation analysis to the data (Supplementary Figure 11a). As assessed by principal component analysis (PCA), there were variable clustering patterns within and in-between samples, showing the highest effect to be based on the treatment at PC1: 36% (Supplementary Figure 11a). ADH-6 treated cell replicates were observed to cluster independently from ADH-1 and C sample replicates. Subsequently, the number of differentially expressed genes (DEGs) was determined based on statistical cut-offs, including *P*-adj < 0.05, *P*-adj < 0.01 and *P*-adj < 0.001 between pairwise comparisons of ADH-6/C, ADH-1/C, and ADH-6/ADH-1. The number of DEGs varied, with the highest observed for ADH-6/C at 485 genes at *P*-adj < 0.05 (Supplementary Figure 11b).

**Figure 6.**
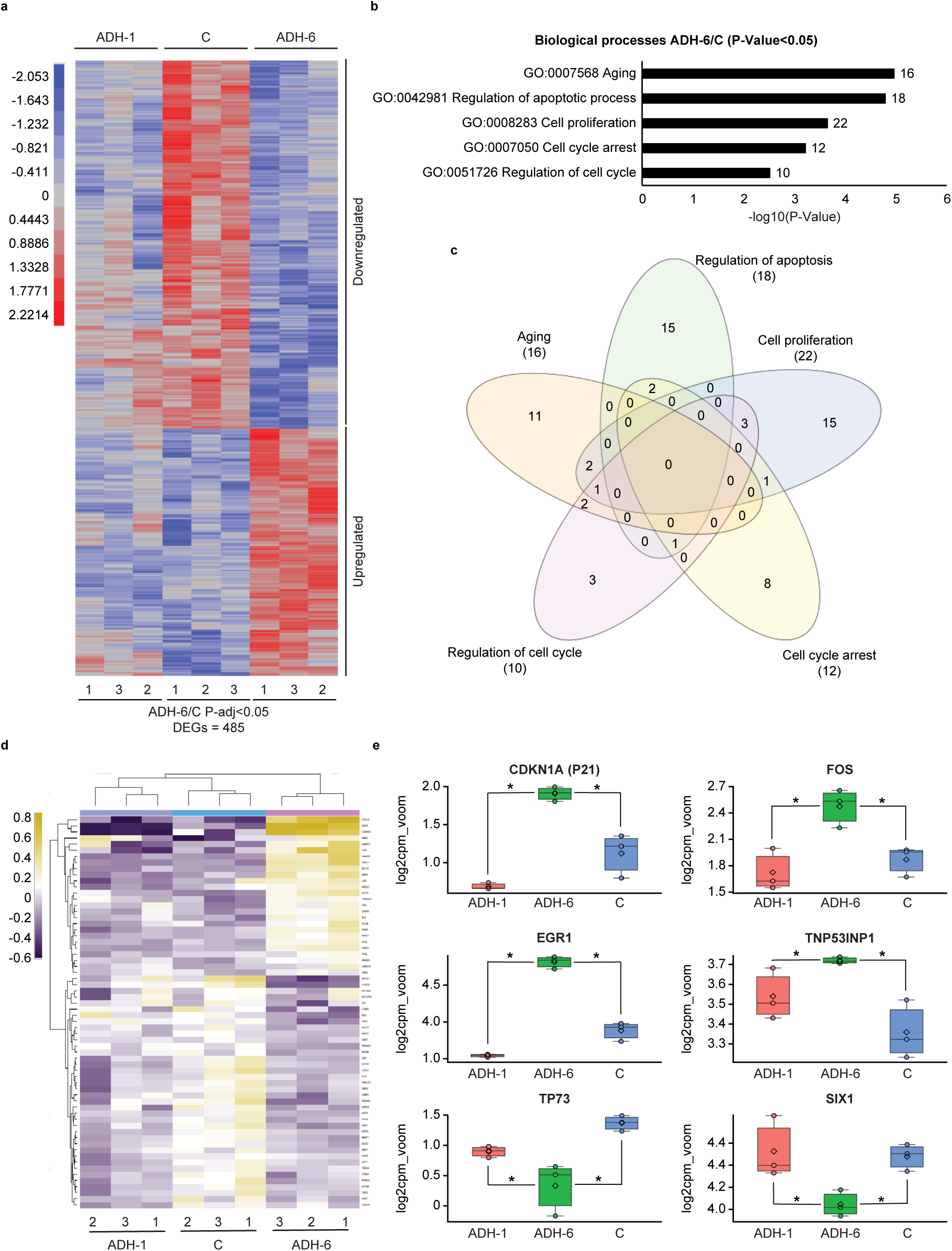
Transcriptome analysis of oligopyridylamide-treated MIA PaCa-2 cells. (**a**) A variance-stabilized transformation (VST) metric heatmap showing expression patterns of DEGs identified based on statistical significance of *P*-adj < 0.05 from the ADH-6 treatment group relative to vehicle-treated controls (C), denoted ADH-6/C. In addition to the ADH-6 treatment group and C, the ADH-1 treatment group was also included. The adjacent legend indicates the scale of expression based on VST count with red signifying upregulation, and blue downregulation, in the ADH-6 group relative to controls. (**b**) Gene ontology (GO) analysis of ADH-6/C (*P*-adj < 0.05) showing biological process term enrichments. Included enrichments are based on a cut-off of *P*-value < 0.05 (normalized to −log10). (**c**) Venn diagram displaying DEGs based on GO term analysis involved in processes including cell cycle, apoptosis, aging and proliferation. (**d**) Metric heatmaps (scaled to log2 mean variance of count per million reads (log2cpm_voom)) based on DEGs from the Venn diagram. (**e**) Gene expression boxplot scaled log2cpm_voom of selected DEGs: Cdkn1a (*top left panel*), Fos (*top right panel*), Egr1 (*middle left panel*), Tp53inp1 (*middle right panel*), Tp73 (*lower left panel*) and Six1 (*lower right panel*). \**P* < 0.05 for comparison amongst the different treatment groups.

To further probe the effects of treatments, the expression patterns of the 485 DEGs identified were analyzed using hierarchical clustering. Following normalization of data count to variance-stabilized transformation (VST) values, distinctive clustering patterns were observed as shown in the metricized heatmap (Figure 6a): 196 DEGs were up-regulated, while 289 DEGs were downregulated, in the ADH-6 treatment group relative to the vehicle treated controls. Looking into the functional aspect of the identified DEGs, gene ontology (GO) term (biological processes) was performed (Figure 6b). Under biological processes, and with a cut-off of *P*-value < 0.05 (normalized by −log10), several enrichments processes were identified, including regulation of cell cycle (GO:0051726), cell cycle arrest (GO:0007050), cell proliferation (GO:0008283), regulation of apoptotic process (GO:0042981) and aging (GO:0007568) (Figure 6b). The selected enrichments were further analyzed, by delineating redundancies between the genes under these conditions (Figure 6c), into a finalized DEGs list. The finalized DEGs list showed variable expression patterns, as indicated in the generated heatmap (Figure 6d). 49 genes that were differentially downregulated, and 25 were differentially upregulated, in the ADH-6 treatment group.

The observed expression patterns indicate transcriptional activation of p53 by ADH-6. For instance, p53 target genes, such as *Cdkn1a, Tp53inp1, Fos* and *Egr1*, were significantly upregulated following treatment with ADH-6 relative to both the ADH-1 and control groups (Figure 6e). *Cdkn1a* (also known as p21) is a cyclin-dependent kinase inhibitor that mediates the progression of the cell cycle in G1 and S phases^43, 44^. Following ADH-6 treatment, *Cdkn1a* was upregulated 0.7–1.3-fold when compared to both control and ADH-1 groups. Interestingly, a 0.4-fold upregulation of *Tp53inp1* was also observed in the ADH-6 treatment group compared to controls. Upon translation, *Tp53inp1* forms a protein complex with homeodomain-interacting protein kinase-2 (HIPK2) or protein kinase C δ (PKCδ) that phosphorylates p53 at residue S46^45^. This stabilizes p53 and enhances its activity, which leads to transcriptional activation of target genes, such as *Cdkn1a*, and subsequent cell growth arrest and apoptosis^45^. Likewise, ADH-6 induced a 0.6–1-fold upregulation of *Fos* relative to treatment with vehicle or ADH-1. *Fos* is part of the activator protein 1 (AP-1) transcription factor complex, which is involved in regulating a number of cellular processes, including differentiation, proliferation, and apoptosis^46^. Of relevance, the first intron of the *Fos gene* contains a p53-responsive element, and overexpression of p53 has been shown to induce *Fos* gene upregulation, which contributes to both p53-mediated apoptosis and cell cycle arrest^46^. Finally, treatment with ADH-6 induced a 0.9–1.3-fold upregulation of *Egr1* compared to both the vehicle and ADH-1 treatment groups. *Egr1* is a transcriptional regulator that can be directly activated by p53 via a non-consensus p53 binding site on the *Egr1* promoter. Although *Egr1* has a range of roles, genotoxic stress-induced upregulation of *Egr1* results in apoptosis in most mammalian cells^47^.

Conversely, treatment with ADH-6 led to significant downregulation of p53 targets *Six1* and *Tp73* compared to controls (Figure 6e). ADH-6 induced a 1-fold downregulation of *Tp73*, a p53 homolog, compared to controls. Interestingly, downregulation of *Tp73*, whose overexpression is associated with advanced-stage cancer^48^, was also observed following treatment of mutant p53-bearing cells with ReACp53^14^. *Six1* is a homeodomain-containing transcription factor that regulates cell migration, invasion and proliferation in progenitor cell populations, but is not expressed in normal adult tissue^49^. Importantly, *Six1* is re-expressed in many cancers and acts as a p53 down-regulator through an MDM2-independent pathway^49^. ADH-6 induced a 0.3-0.5-fold downregulation of *Six1* relative to both controls and ADH-1 treatment groups (Figure 6e).

Other enrichments, such as isoprenoid biosynthesis (GO:0008299), lipid biosynthesis (GO:0008610) and cholesterol biosynthesis (GO:0006695) were also identified in ADH-6 treated cells (Supplementary Figure 11c,d). The heatmap generated from genes involved in these processes revealed upregulation of the mevalonate pathway, which is essential for cancer cell survival and growth^50^. Of relevance, the mevalonate pathway is mediated by non-aggregated mutant p53^51^. Thus, ADH-6 mediated release of overexpressed mutant p53 from the cellular inclusions likely leads to activation of some oncogenic pathways. However, the bulk of rescued mutant p53 behaves similar to the WT protein (Figure 4h) and, possibly along with released p63 and p73, upregulates major tumor-suppressive pathways (Figure 6), which overwhelms the pro-tumor activities and results in the observed inhibition of proliferation via cell cycle arrest and induction of apoptosis (Figure 5).

Lastly, we performed ingenuity pathway analysis (IPA) of the DEGs to identify the upstream transcriptional regulators (TRs) of these genes. Over a dozen TRs were activated in the ADH-6 group, and the 10 candidates shown were identified on the basis of their q-value and z-score thresholds (Supplementary Figure 12a,b). Of these TRs, *TP53* showed robust activation as indicated by the highest q-value in both the ADH-6 vs ADH-1 (Supplementary Figure 12a) and ADH-1 vs control (Supplementary Figure 12b) comparisons. 44 genes in the ADH-6 vs ADH-1 comparison (Supplementary Figure 12c) and 42 in the ADH-6 vs control comparison showed an expression pattern that signified the activation of the *TP53* pathway. To further determine the biological consequence of ADH-6 mediated p53 activation, gene set enrichment analysis (GSEA) was performed. The findings of IPA were corroborated using this approach as the *TP53* pathway was shown to be induced in the ADH-6 treatment group when compared to the ADH-1 group. Activation of the *TP53* pathway was strongly supported by the observed suppression of the *MYC* pathway (Supplementary Figure 12d), which is known to be negatively regulated by p53^52^. Furthermore, the gene set for cell death/apoptosis was induced and the genes involved in cell cycle progression (G2/M) were suppressed in the ADH-6 treatment group (Supplementary Figure 12d), both of which give credence to activation of the *TP53* pathway by the oligopyridylamide. Taken together, our findings strongly support the view that ADH-6 treatment activates the p53 transcriptional response in mutant p53 bearing cancer cells.

### ADH-6 downregulates cancer-promoting phosphoproteins

We subsequently carried out proteome analysis of oligopyridylamide-treated MIA PaCa-2 cells. We chose to focus specifically on the phosphoproteome (Supplementary Figure 13) as many phosphoproteins are involved in regulating major pathways implicated in cancer^53–57^. Unsupervised hierarchical clustering (UHC) revealed two distinct expression profiles pertaining to the ADH-1 and the ReACp53/ADH-6 treatment groups (Figure 7a). PCA analysis further highlighted that the main source of variation in the two groups was the peptide/compound treatment (Figure 7a,b). Furthermore, compared to ADH-1, most of the phosphoproteins downregulated upon ReACp53 treatment were also downregulated in the ADH-6 samples (Figure 7c). Next, we carried out GSEA of hallmark signatures for the differentially expressed and downregulated phosphoproteins. We observed a comparable reduction of phosphoprotein expression in a number of pathways in both the ReACp53 and ADH-6 treated cells (Figure 7d).

**Figure 7.**
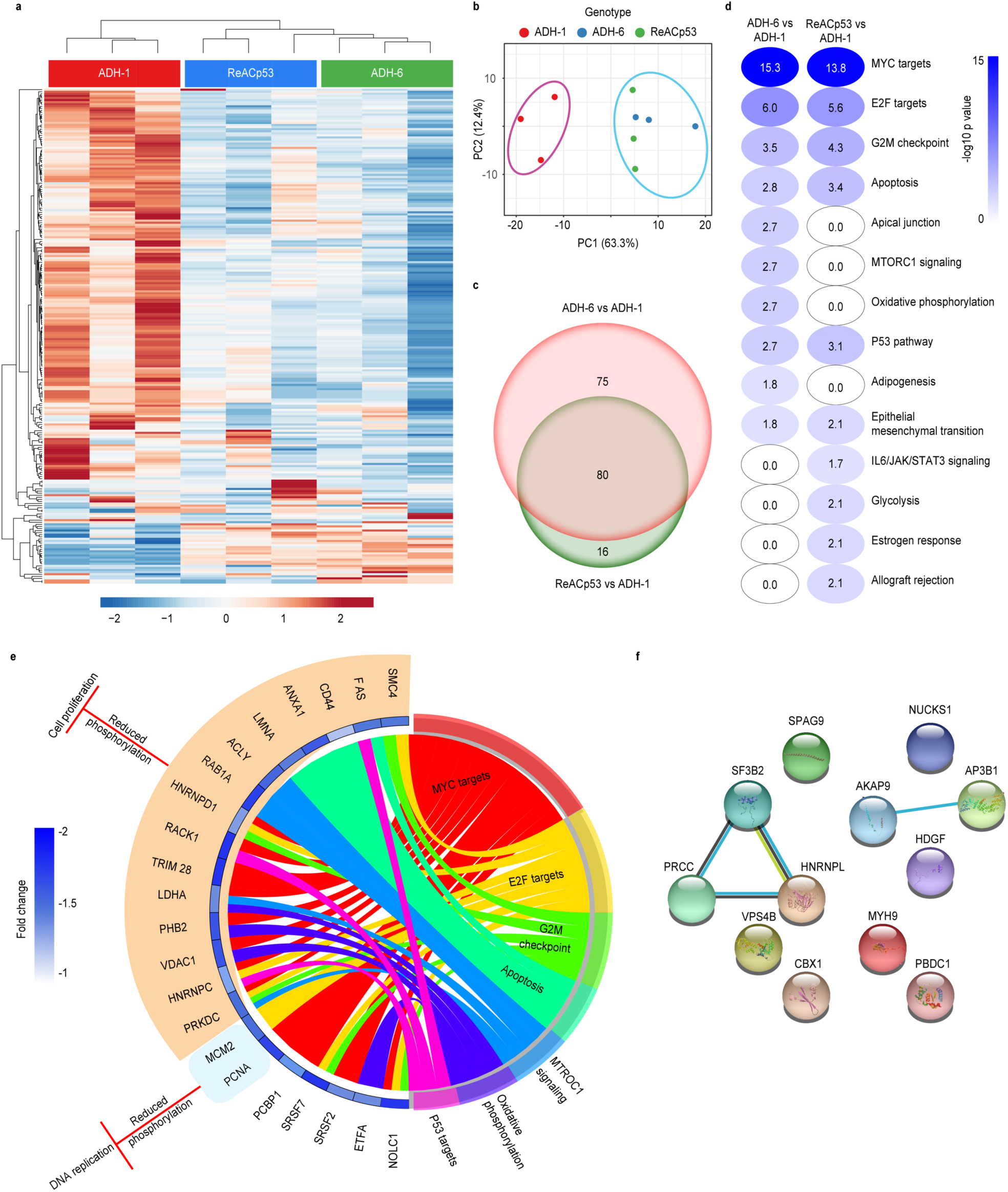
Quantitative proteomic analysis of phosphoprotein expression in oligopyridylamide-treated MIA PaCa-2 cells. (**a**) Heatmap showing differentially expressed phosphoproteins in ADH-1, ReACp53 and ADH-6 treated samples. Unsupervised hierarchical clustering reveals distinct segregation of ADH-1 vs ReACp53 and ADH-6. (**b**) Results from principal component analysis (PCA) showing that the main source of variation in the indicated groups was the compound/peptide treatment. (**c**) Venn diagram showing phosphoproteins that were differentially expressed in the ADH-6 vs ADH-1 and ReACp53 vs ADH-1 comparisons. The number of phosphoproteins that were unique or common in both comparisons are highlighted. (**d**) Gene set enrichment overlap analysis (GSEA) of hallmark signatures for the differentially expressed and downregulated phosphoproteins. The hallmark signatures were ranked based on the −log10 q-value of overlap. Only gene signatures that were significant (q<0.05 or −log10 q-value more than 1.3) were further analyzed. (**e**) GOChord Plot of phosphoproteins that belong to top dysregulated hallmark signatures (shown in **d**). The plot also depicts overlap of phosphoproteins between indicated gene signatures. Fold change of phosphoproteins (ADH-6 vs ADH-1) is represented by the blue track with the color spectrum depicting the level of reduction in phosphoprotein expression in the ADH-6 group. The biological roles of the downregulated phosphoproteins in ADH-6 were inferred from published data. The outer layer of the GOChord plot links the downregulation of phosphoproteins with the inhibition of DNA repair/replication and cell proliferation on the basis of previous reports (Supplementary Table 1a). (**f**) Protein interaction map of upregulated phosphoproteins in ADH-6 vs ADH-1 treatments (biological roles of upregulated phosphoproteins are summarized in Supplementary Table 1b).

The biological roles of the downregulated phosphoproteins were inferred from published data (Supplementary Table 1a), and revealed that these proteins can be divided into two major functional groups that positively regulate either DNA replication/repair or cell cycle progression/proliferation, although many downregulated proteins in the ADH-6 treatment group had overlapping gene signatures (Figure 7e). Importantly, these two major groups, which play a critical role in cancer initiation and progression, are known to be directly regulated by p53^58–62^. This strongly supports that both the ReACp53 and ADH-6 treatments result in reactivation of p53, which downregulates key cancer-promoting phosphoproteins (Supplementary Table 1a). Indeed, numerous reports have shown that p53 directly downregulates the expression of positive regulators of cancer in the p53 pathway, such as CD44^63^ and PCNA^64, 65^, as well as the oncoproteins c-Myc^52, 66^, E2F^67, 68^ and mTORC1^69–71^ (Supplementary Figure 14). Interestingly, only a few upregulated phosphoproteins were identified in our phosphoproteome screen (Figure 7f; Supplementary Table 1b). Together, the transcriptomic and proteomic analyses clearly demonstrate that ADH-6-mediated dissociation of mutant p53 amyloid-like aggregates in cancer cells restores p53 function, leading to cell cycle arrest and activation of apoptosis (Figure 5).

### *In vivo* administration of ADH-6 causes regression of mutant p53-bearing tumors

Having established that ADH-6 potently abrogates mutant p53 amyloid formation and restores WT-like tumor suppressor function *in vitro*, we next assessed the *in vivo* efficacy of the oligopyridylamide (Figure 8 and Supplementary Figure 15). Following intraperitoneal injection, ADH-6 quickly entered circulation, with the peak concentration in serum mice (∼21 µg/mL) occurring at 2 h post-injection (Figure 8a). The *in vivo* circulation half-life^72^ of ADH-6 (t_1/2_ = ∼3.6 h) was much longer than that of ReACp53 (t_1/2_ ∼1.5 h)^14^, or other chemotherapeutics of a comparable size, such as doxorubicin (t_1/2_ < 30 min)^73^ or paclitaxel (t_1/2_ ∼1.7 h)^74^. Moreover, ADH-6 was detected in the plasma up to 48 h after administration, whereas ReACp53 was eliminated from the bloodstream in 24 h^14^. The relatively long *in vivo* circulation time of ADH-6 should facilitate greater accumulation in tumor tissue. Indeed, the amount of ADH-6 in the MIA PaCa-2 xenografts increased continuously over 48 h post treatment (Figure 8b). The high *in vivo* stability suggested that ADH-6 would exhibit potent antitumor activity.

**Figure 8.**
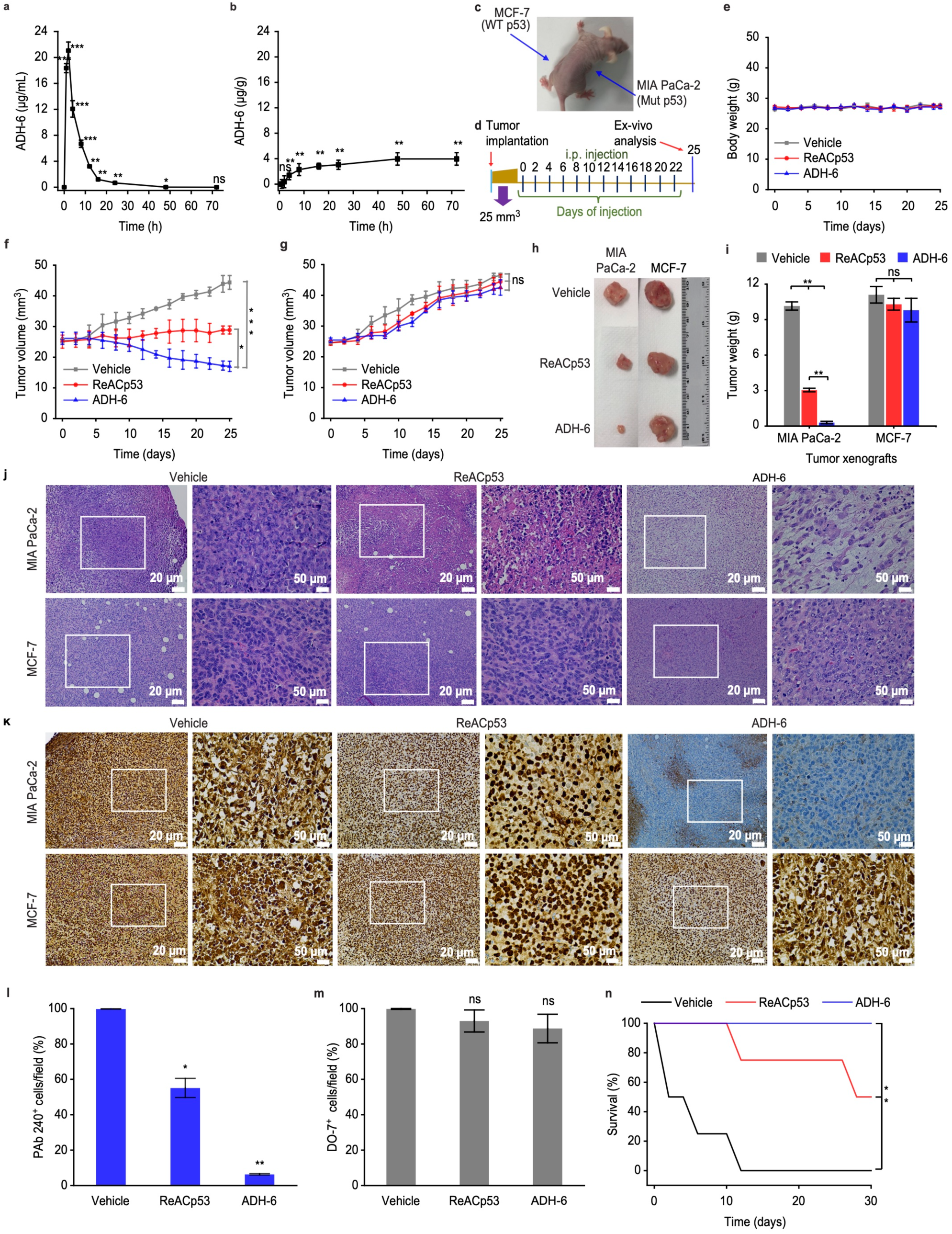
ADH-6 causes regression of tumors bearing mutant, but not WT, p53. (**a,b**) *In vivo* pharmacokinetics of ADH-6. Concentration of ADH-6 in plasma (**a**) and in MIA PaCa-2 xenografts **(b)** of mice (*n* = 5 per group), after an intraperitoneal injection of ADH-6 (15 mg kg^-1^), was quantified using LC–MS/MS^132^. (**c,d**) Design of the tumor reduction studies. A representative mouse bearing both MIA PaCa-2 (mutant R248W p53) and MCF-7 (WT p53) xenografts (**c**) and treatment schedule for the dual xenograft model (**d**). Once the tumor volume reached ∼25 mm^3^, the mice were randomized into the different treatment groups (*n* = 8 per group), which were injected intraperitoneally with vehicle (0.02% DMSO), ReACp53 (716.4 µM) or ADH-6 (716.4 µM). Injections were done every 2 days for a total of 12 doses, with the first day of treatment defined as day 0. (**e**) Bodyweight changes of the tumor-bearing mice in the different treatment groups monitored for the duration of the experiment. (**f,g**) Tumor volume growth curves for the MIA PaCa-2 (**f**) and MCF-7 (**g**) xenografts in the different treatment groups over 25 days of treatment (*n* = 8 per group). Tumor volume was measured via high-precision calipers using *Equation 3*. (**h,i**) Tumor mass analysis for the different treatment groups. After 25 days of treatment, 4 mice per treatment group were sacrificed and the tumor tissues were isolated and imaged (**h**) and subsequently measured weighed to determine the tumor mass (**i**). (**j**) Hematoxylin and eosin (H&E)-stained images of xenograft sections from the different treatment groups following 25 days of treatment. Images on the right are magnified views of the boxed regions in the images on the left. Scale bar = 20 μm (50 μm for the magnified views). (**k–m**) Immunohistochemistry (IHC) analysis of the residual xenografts. Images of sections of MIA PaCa-2 and MCF-7 xenografts stained using the anti-p53 PAb 240 and DO-7 antibodies, respectively, from the different treatment groups (**k**). Images on the right are magnified views of the boxed regions in the images on the left. Scale bar = 20 μm (50 μm for the magnified views). Quantification of PAb 240 (**l**) and DO-7 (**m**) positive cells in 3–5 different fields of view expressed as % of the total number of cells (*n* = 4). (**n**) Survival curves for the vehicle, ReACp53 and ADH-6 treatment groups over 30 days (*n* = 4 per group). \**P* < 0.05, \*\*\**P* < 0.001 or non-significant (ns, *P* > 0.05) for comparison with controls and amongst the different treatment groups.

To test this hypothesis, mice bearing MIA PaCa-2 xenografts were treated with ADH-6, ADH-1 or ReACp53 (155.6 µM in PBS) (Supplementary Figure 15a,b). Treatment consisted of intraperitoneal (IP) injections every 2 days, for a total of 12 doses (Supplementary Figure 15b). IP injection was chosen as it was the administration route for the control ReACp53 peptide^14^. As expected, ADH-1 did not affect tumor growth (Supplementary Figure 15d–f,h). Both ADH-6 and ReACp53 substantially reduced tumor growth relative to the saline-treated control group. However, of the two treatments, ADH-6 exhibited significantly greater antitumor efficacy (Supplementary Figure 15d–f,h). Moreover, ADH-6 prolonged survival appreciably compared to ReACp53 (Supplementary Figure 15g).

To confirm its specificity for aggregation-prone mutant p53 *in vivo*, we tested ADH-6 on a dual xenograft model: mice bearing MIA PaCa-2 (mutant R248W p53) xenografts on one flank and MCF-7 (WT p53) xenografts on the other flank as an internal control (Figure 8c). For this model, treatment consisted of IP injections every 2 days, for a total of 12 doses, of ADH-6 or ReACp53 (716.4 µM in 0.02% DMSO) or vehicle (0.02% DMSO) (Figure 8d). DMSO was introduced to increase the solubility of ADH-6 and allow administration of a higher dose of the oligopyridylamide and ReACp53, which was comparable to that used in the study by Eisenberg *et al*.^14^ While treatment with ADH-6 or ReACp53 did not have a significant effect on growth of the MCF-7 xenografts, both the protein mimetic and peptide markedly reduced the MIA PaCa-2 tumor growth relative to the vehicle treated control group (Figure 8f–i). Following treatment with ReACp53, the tumors increased from an initial volume of 25 ± 2 to 29 ± 1 mm^3^ (compared to an increase from 25 ± 1 to 44 ± 2 mm^3^ in the vehicle treated controls; Figure 8f), while the tumor mass was ∼30% that of the controls (Figure 8h,i). However, ADH-6 again exhibited significantly greater antitumor efficacy than ReACp53. Treatment with the oligopyridylamide reduced the MIA PaCa-2 xenografts from an initial volume of 26 ± 2 to 17 ± 2 mm^3^ after treatment (Figure 8f), while the tumor mass was ∼3% that of the controls (Figure 8h,i). Histological analysis of tumor tissues using hematoxylin and eosin (H&E) staining confirmed the greater potency of ADH-6 compared to ReACp53 against MIA PaCa-2 xenografts (Figure 8j and Supplementary Figure 15h). Immunohistochemistry (IHC) analysis of residual tumor sections revealed that treatment with ADH-6 markedly decreased mutant p53 levels in the MIA PaCa-2 xenografts, but did not significantly alter the amount of WT p53 in MCF-7 xenografts (Figure 8k– m). Thus, ADH-6 effectively targets aggregation-prone and inactive mutant, but not functional WT, p53 *in vivo*.

ADH-6 and ReACp53 did not adversely affect the bodyweight of treated mice (Figure 8e and Supplementary Figure 15c), and H&E-stained heart, lung, liver, kidney and spleen sections showed no apparent abnormalities or lesions (Supplementary Figures 16 and 17). Our observations are in agreement with the reported high tolerability and lack of measurable toxicity in healthy tissue *in vivo* of ReACp53^14^. Taken together, these results demonstrate that ADH-6 potently shrinks xenografts harboring aggregation-prone mutant p53 *in vivo*, leading to prolonged survival, while exhibiting no toxicity to healthy tissue.

## DISCUSSION

A wide range of disorders are associated with misfolding and self-assembly of functional proteins or peptides into amyloids, which are aggregates that are characterized by a fibrillar morphology, a predominantly β-strand secondary structure, high thermodynamic stability, insolubility in common solvents and resistance to proteolytic digestion^75–77^. These amyloid diseases include AD, T2D, Parkinson’s disease, transmissible spongiform encephalopathies (or prion diseases) and Huntington’s disease^76, 77^. Rather unexpectedly, it is now evident that a subset of mutant p53-associated cancers can also be classed as amyloid diseases. The conformationally flexible p53 normally exists in an equilibrium between native/folded, partially unfolded and aggregated states^14, 78^. At the core of p53 is the thermodynamically unstable DBD, which houses the vast majority of cancer-associated mutations^9, 10^. Many of these mutations destabilize the DBD further and prompt its unfolding, leading to exposure of the normally hidden aggregation-nucleating subdomain, p53_251–258_^13, 14^. This shifts the equilibrium towards the aggregated state^13, 14^, culminating in sequestration of mutant p53 in inactive cytosolic amyloid-like aggregates that often coopt the WT isoform and its paralogs, p63 and p73^17^.

The amyloidogenic nature of mutant p53 aggregation was first reported almost two decades ago^79^. Subsequent studies revealed amyloid-like mutant p53 aggregates in cancer cell lines, *in vivo* animal models and human tumor biopsies^11, 12^. Increasing evidence now implicates these aggregates in mutant p53-associated loss of tumor suppressor function and oncogenic GoF^3, 5, 12, 14, 17, 19^. Intriguingly, there are reports of transmission of this oncogenic GoF phenotype, whereby mutant p53 amyloid aggregates (in the form of oligomers or fibril fragments) are trafficked to contiguous cells and seed aggregation of endogenous p53, suggesting that aggregation-prone mutant p53 bearing cancers share a common mechanism of propagation with prion diseases^11, 12^. It is therefore rather surprising that, so far, there are only a handful of reports of therapeutic strategies targeting mutant p53 aggregation. The most notable of these is ReACp53, a sequence-specific peptide inhibitor that blocks mutant p53 aggregation by masking the DBD’s aggregation-nucleating p53_251-258_ segment^14, 17^. This prevents further aggregation and shifts the folding equilibrium toward p53’s functional, WT-like state. Treatment with ReACp53 rescued p53 function in aggregation-prone mutant p53-bearing human ovarian and prostate cancer cells, leading to inhibition of cell proliferation *in vitro* and tumor shrinkage *in vivo*^14, 41^. More recently, a bifunctional small molecule (denoted L^I^), with a structure based on the amyloid reporter ThT, was reported to restore zinc binding in mutant p53 and to modulate its aggregation^80^. L^I^ inhibited mutant p53 aggregation and restored the protein’s transcriptional activity, leading to apoptosis in human gastric cancer cells *in vitro*. While promising, the *in vivo* efficacy of L^I^ is yet to be determined.

Here, we have extended our protein mimetic-based approach, previously developed to modulate various aberrant PPIs^20–22^, towards mutant p53 aggregation. Screening a focused library of oligopyridylamide-based α-helix mimetics, we identified a cationic tripyridylamide, ADH-6, that potently inhibited oligomerization and amyloid formation of pR248W, a mutant p53 DBD-derived peptide containing both the aggregation-nucleating sequence and the commonly occurring R248W mutation, by stabilizing the peptide’s native conformation (Figure 1). It should be noted that ADH-6 only modestly antagonized Aβ amyloid formation^20^, indicating a degree of specificity of the oligopyridylamide for mutant p53. Notably, ADH-6 also dissociated pre-formed pR248W aggregates, and prevented further aggregation of the peptide. Subsequent studies in human pancreatic carcinoma MIA PaCa-2 cells harboring mutant R248W p53 (Figure 4), and *N. benthamiana* cells transfected with YFP-tagged WT and R248W p53 DBDs (Supplementary Figure 9), showed that ADH-6 effectively dissociates intracellular mutant p53 amyloid-like aggregates (Figure 4). Consistent with the aggregation assays and intracellular imaging, molecular docking simulations revealed that the interaction of ADH-6 with the hydrophobic core of mutant p53 DBD is stronger than that of DBD– LTIITLE (i.e. DBD self-interaction), thereby providing a thermodynamic basis for ADH-6’s amyloid inhibition mechanism (Figure 2). The simulations also revealed that the DBD–ADH-6 interaction is energetically favored over DBD–ReACp53. The stronger DBD–ADH-6 interaction, which should favor a long-lived adduct, partly accounts (*vide infra*) for the greater efficacy of the oligopyridylamide, relative to ReACp53, at inhibiting mutant p53 aggregation and shifting the folding equilibrium towards soluble, functional p53 (Figure 4 and Supplementary Figures 8 and 9).

NMR spectroscopy was used to gain insights into the interactions between ADH-6 and mutant R248W p53 DBD at the molecular level. The experiments revealed that the α-helix mimetic interacts with not only the aggregation-nucleating subdomain, but also several other regions of mutant p53 DBD (Figure 3). This is not surprising given that the DBD contains several helical regions (Figure 1), as well as non-helical regions in which amino acids – from neighboring or distal parts of the protein – nevertheless align to present target side-chains with a topology that matches that of the surface functionalities of ADH-6 (Figure 3). On the other hand, ReACp53, by virtue of its sequence, specifically targets the aggregation-nucleating subdomain (p53_251-258_) of the DBD^14^. However, it has been shown that mutant p53 unfolding may expose other aggregation-prone sites, besides p53_251-258_, which can contribute to the protein’s self-assembly and enable it to partially circumvent the inhibitory effects of ReACp53^81^. Thus, the combination of targeting of multiple regions of mutant p53 DBD, and stronger interaction with the hydrophobic core of the DBD, enables ADH-6 to antagonize intracellular mutant p53 aggregation more effectively than ReACp53.

Helical intermediates play a role in the amyloid assembly of not only Aβ and IAPP, but also a number of other disease-associated intrinsically disordered proteins, including *α*-synuclein (Parkinson’s disease), prion protein (PrP, prion diseases) and tau (tauopathies)^82–84^. A distinct possibility is that the intrinsically disordered mutant p53 aggregates via a similar pathway, whereby destabilized α-helical intermediates transition to stable, β-sheet rich amyloid aggregates. Indeed, there are reports that aggregation-prone p53 mutants, and fragments of these mutants, sample helical structures during the process of self-assembling into amyloid fibers^85, 86^. This raises the possibility that, in addition to the monomeric protein, ADH-6 also targets pre-amyloid helical intermediates of mutant p53’s amyloid aggregation pathway. By stabilizing mutant p53 in its monomeric or helical intermediate states, ADH-6 is able to effectively prevent the protein’s amyloid aggregation, thereby strongly shifting the folding equilibrium towards the functional, WT-like state.

Several lines of evidence from multiple, complementary, experiments strongly suggest that ADH-6-rescued mutant p53 behaves similarly to its functional WT counterpart: (i) a second screen, testing the effects of the oligopyridylamides on viability of mutant R248W p53 bearing pancreatic carcinoma MIA PaCa-2 cells (Figure 5 and Supplementary Figure 10), revealed a strong correlation between the compounds’ cytotoxicity and their capacity to antagonize pR248W aggregation (Figure 1 and Supplementary Figure 1), and again identified ADH-6 as the most potent compound in the library; (ii) ADH-6 is significantly more toxic to MIA PaCa-2 cells than the control ReACp53 peptide (Figures 5 and Supplementary Figure 10), which reflects their relative capacities to abrogate mutant p53 aggregation (Figure 4 and Supplementary Figure 8 and 9); (iii) ADH-6 induced substantial loss of viability in a range of human cancer cells bearing mutant p53, but was completely nontoxic to WT p53 harboring human cancer cells (Supplementary Figure 10); (iv) p53 null SKOV-3 cells were insensitive to ADH-6 treatment but, upon transfection with mutant R248W p53, the cells became susceptible to ADH-6 mediated cytotoxicity (Figure 5); and (v) further analysis of its effects on mutant p53 harboring cancer cells revealed that ADH-6 induces cell cycle arrest and apoptosis, both of which are indicators of restored p53 function (Figure 5 and Supplementary Figure10).

RNA-Seq confirmed that ADH-6 treatment resulted in transcriptional reactivation of p53 in MIA PaCa-2 cells (Figure 6 and Supplementary Figures 11 and 12). Specifically, we observed activation of the *TP53* and apoptosis pathways, and suppression of genes involved in cell cycle progression (G2/M) and the MYC pathway, which are known to be negatively regulated by p53^52^. More in-depth analysis revealed upregulation of specific p53 target genes, such as *Cdkn1a, Tp53inp1, Fos* and *Egr1*. Conversely, treatment with ADH-6 led to significant downregulation of *Tp73,* which was also observed following treatment of mutant p53-bearing cells with ReACp53^14^, and the *Six1* oncogene. Consistent with the transcriptome analysis, the proteome analysis showed that ADH-6 downregulates key cancer-promoting phosphoproteins that are known to be directly negatively regulated by p53^58–62^ (Figure 7, Supplementary Figures 13–15 and Supplementary Table 1). Thus, together with the aggregation assays, intracellular imaging and cell viability/toxicity results, the transcriptome and proteome analyses clearly demonstrate that ADH-6 dissociates mutant p53 amyloid-like aggregates in cancer cells, and that the released protein is restored to a functional form, which elicits the observed inhibition of proliferation via cell cycle arrest and induction of apoptosis (Figure 5).

To establish whether the *in vitro* effects of ADH-6 are recapitulated *in vivo*, we evaluated the effects of ADH-6 on mice bearing MIA PaCa-2 (mutant R248W p53) tumors, alone or with MCF-7 (WT p53) tumors on the opposite flank as an internal control (Figure 8 and Supplementary Section 5). ReACp53, serving as a positive control, reduced MIA PaCa-2 tumor growth and prolonged survival relative to the vehicle treated control groups, which is in agreement with the peptide’s reported ability to inhibit growth of aggregation-prone mutant bearing tumors^14^. However, ADH-6 was markedly more effective at decreasing MIA PaCa-2 tumor volume and mass, and prolonging survival, compared to ReACp53. The greater *in vivo* efficacy of ADH-6 compared to ReACp53 correlates well with their relative capacities to dissociate intracellular mutant p53 aggregates (Figure 4) and to induce toxicity in mutant p53 bearing cancer cells (Figure 5 and Supplementary Figure 10), as well as with their relative *in vivo* stabilities. Peptides possess a number of pharmaceutically desirable properties, including the ability to selectively bind to specific targets with high potency, thereby minimizing off-target interactions and reducing the potential for toxicity^87^. However, a major disadvantage of peptides is their low *in vivo* stability^73^. Small molecules, with their constrained backbone, are inherently more stable than peptides^24, 88^, which is reflected in the substantially longer *in vivo* circulation half-life and prolonged presence in the bloodstream of ADH-6 relative to ReACp53. The extended *in vivo* circulation time of ADH-6 facilitates greater accumulation in tumor tissue and increased anti-tumor activity. Importantly, ADH-6 did not affect growth of control MCF-7 xenografts, underlining the oligopyridylamide’s specificity for tumors that harbor aggregation-prone mutant p53. Additionally, detailed necropsies of major organs revealed no damage or alterations, demonstrating the lack of toxicity of ADH-6 to healthy tissue. Taken together, our results show that ADH-6 effectively shrinks tumors bearing aggregation-prone mutant p53 *in vivo*, and prolongs survival, without displaying the non-specific toxicity that is common to conventional cancer therapeutics.

Mutant p53-associated cancers are predicted to lead to the deaths of more than 500 million people alive today^89^. This is fueled, in part, by the increasing incidences of notoriously difficult to treat malignancies, such as the highly lethal pancreatic cancer, which is on course to becoming the second leading cause of cancer-related mortality in Western countries within the next 5–10 years^90^. Consequently, there is a pressing need for new therapeutic strategies to supplement or supplant current cancer treatments. In the present study, we focused on a largely neglected – with a few notable exceptions – property of a sizeable subset of p53 mutants, namely their propensity to self-assemble into amyloid-like aggregates. This aggregation is implicated in mutant p53-associated tumor suppressor loss of function, oncogenic GoF and, potentially, prion-like propagation of these phenotypes^3, 5, 11, 12, 14, 17, 19^. Testing protein mimetics originally designed to antagonize amyloid formation associated with AD and T2D^20–22^, we identified a cationic tripyridylamide, ADH-6, that effectively abrogates mutant p53 amyloid-like aggregation in human cancer cells, which restores p53’s transcriptional activity, leading to cell cycle arrest and induction of apoptosis. Importantly, ADH-6 treatment causes regression of xenografts harboring mutant, but not WT, p53 and prolongs survival, with no visible toxicity to healthy tissue. This study effectively establishes a bridge between amyloid diseases and cancer, providing a foundation for cross-informational approaches in the design of potent mutant p53-targeted cancer therapeutics.

## MATERIALS AND METHODS

### Reagents

p53-derived peptides (p53p and R248Wp) and ReACp53 (sequences shown in Figure 1a) were synthesized by Selleck Chemicals (Houston, TX) using standard Fmoc methods. 2-deoxy-D-glucose, acetone, acetonitrile (ACN), 3′,5′-Dimethoxy-4′-hydroxyacetophenone (acetosyringone), bovine serum albumin (BSA), calcium chloride (CaCl_2_), chloroform, dimethyl sulfoxide (DMSO), 1,4-dithiothreitol (DTT), ethylenediaminetetraacetic acid (EDTA), gentamycin, glucose, glycolic acid, N-(2-hydroxyethyl) piperzine-N’-(2-ethanesulfonic acid) (HEPES), 1,1,1,3,3,3-hexafluoro-2-propanol (HFIP), indole-3-acetic acid (IAA), isoflurane, kanamycin, magnesium chloride (MgCl2), methanol, 2-(N-morpholino)ethanesulfonic acid (MES), N-hydroxysuccinimide (NHS), paraformaldehyde, phosphate buffered saline (PBS), phosphotungstic acid (PTA), rifampicin, Sephadex G-25 DNA Grade column, sodium azide, sodium bicarbonate (NaHCO3), sodium chloride (NaCl), sodium hydroxide (NaOH), triethylammonium bicarbonate (TEAB), thioflavins S and T (ThS and ThT), trifluoroacetic acid (TFA), Triton X-100, Trizma, Tween 20 and Trypan Blue were all purchased from Sigma-Aldrich (St. Louis, MO). C18 Tips, Dead Cell Apoptosis kit, formic acid, G418 sulfate (Geneticin), Hoechst 33342, hydroxylamine, Lipofectamine LTX Reagent, Lurai-bertani (LB), Opti-MEM Reduced Serum Medium, Pierce BCA Protein Assay kit, Pierce Trypsin/Lys-C Protease Mix and tandem mass tag (TMT)-labeling kit were from Thermo Fisher Scientific (Waltham, MA). CellTiter 96 AQueous One Solution (MTS) Cell Proliferation Assay was purchased from Promega (Madison, WI).

### Synthesis of oligopyridylamide-based protein mimetics

The protocols for synthesis, fluorescent dye-labeling and characterization of the relevant oligopyridylamides are presented in Supplementary Section 6. For each experiment, fresh stocks of the oligopyridylamides (10 mM) were prepared in DMSO and filtered using 0.22 μm Ultrafree-MC spin filters (Sigma).

### Peptide preparation

Peptides (pWT, pR248W and ReACp53; sequences shown in Figure 1a) were synthesized by Selleck Chemicals (Houston, TX) using standard Fmoc methods. The peptides were purified inhouse by reverse-phase high-performance liquid chromatography (Waters 2535 QGM HPLC), and purity was subsequently verified using mass spectrometry (Agilent 6538 QToF LC/MS). After purification, peptides were aliquoted into 0.5 mL Protein LoBind tubes (Eppendorf), lyophilized and stored at −80 °C until needed. For each experiment, fresh peptide stock solutions were prepared in DMSO and filtered using 0.22 μm Ultrafree-MC spin filters. pR248W concentration was determined by absorbance measurements at 280 nm (ε = 5690 cm^−1^ M^−1^ for tryptophan) on a Lambda 25 UV/Vis Spectrophotometer (PerkinElmer, Waltham, MA) using quartz cuvettes (1 cm path-length).

### Thioflavin T (ThT)-based aggregation assay

Peptide (pWT and pR248W) amyloid formation kinetics were measured in quadruplicate in black 96-well plates with flat bottom (Corning Inc., NY) using a Synergy H1MF Multi-Mode microplate reader (BioTek, Winooski, Vermont). The aggregation was initiated by dilution of peptide from a freshly prepared stock solution (1 mM in DMSO) to PBS containing ThT, with or without oligopyridylamides. For some experiments, the oligopyridylamides were added (from a stock solution of 10 mM in DMSO) at the indicated timepoint after the start of the aggregation reaction. To maintain identical conditions, an equal amount of DMSO was added to the wells with peptide only reactions. Final concentrations in the wells were: 25 μM pWT or pR248W; 0 or 25 μM oligopyridylamide; 50 μM ThT. Peptide aggregation was monitored by shaking and measuring the ThT fluorescence (λ_ex/em_ = 440/480 nm) at 5-min intervals at 37 °C. The sample data were processed by subtracting the blank and renormalizing the fluorescence intensity by setting the maximum value to one.

### Transmission electron microscopy (TEM)

25 μM p248W was incubated, alone or co-mixed with an equimolar concentration of ADH-1, ADH-6 or ReACp53, for 10 h at 37 °C. Thereafter, the solution was vortexed and a 5 μL droplet of the solution was placed on a freshly plasma-cleaned copper/formvar/carbon grid (400 mesh, Ted Pella, Redding, CA). After 2 min, the droplet was wicked away by carefully placing the grid perpendicularly on a filter paper. The grid was then gently washed by dipping the upper surface in a water droplet and dried as described above using a filter paper. Next, the grid was placed upside down for 20 s in a 1% phosphotungstic acid (PTA) solution that had been passed through a 0.2 μm filter and centrifuged for 10 min at 14,000 rpm to remove potential PTA agglomerates. The PTA solution was wicked away and the grid was further dried for 10 min under cover at room temperature. Images of at least five different grid regions were acquired per experimental condition on a Talos F200X TEM (Thermo Fisher Scientific) equipped with a Ceta 16M camera operated at an accelerating voltage of 200 kV. Velox software (Thermo Fisher Scientific) was used for image analysis.

### Dot blot immunoassay

Samples of 10 μM pR248W, in the absence or presence of an equimolar amount of the ligands (ADH-1 or ADH-6), were incubated for various durations. The samples were then applied to a nitrocellulose membrane and dried for 1 h at room temperature or overnight at 4 °C. The membranes were subsequently blocked with 5% nonfat milk in TBST buffer (10 mM Tris, 0.15 M NaCl, pH 7.4, supplemented with 0.1% Tween 20) for 2 h at room temperature, washed thrice with TBST buffer and incubated overnight at 4 °C with the polyclonal A11 antibody (1/1000 dilution in 5% nonfat milk in TBST buffer; Life Technologies Corp., Grand Island, NY). Next, the samples were washed with TBST buffer (×3) and incubated with horseradish peroxidase (HRP)-conjugated anti-rabbit IgG (1/500 dilution in 5% nonfat free milk in TBST buffer; Santa Cruz Biotechnology, Dallas, TX) at room temperature for 1 h. The dot blots were then washed with TBST buffer (×5), developed using the ECL reagent kit (Amersham, Piscataway, NJ), and finally imaged using a Typhoon FLA 9000 instrument (GE Healthcare Life Sciences, Pittsburgh, PA) with the settings for chemiluminescence.

### Circular dichroism (CD) spectroscopy

Each freshly prepared pR248W stock solution was diluted in PBS to a concentration of 10 μM, alone or with an equimolar concentration of the ligand (ADH-1 or ADH-6), and transferred to a 1-mm quartz cuvette. CD spectra were recorded from 190 to 260 nm, with a bandwidth of 0.5 nm, a step size of 0.5 nm and a time-per-point of 10 s, at 25 °C using a Chirascan Plus Circular Dichroism Spectrometer (Applied Photophysics Limited, UK) with a Peltier temperature control system. For each experiment, the CD spectra from three independent trials were averaged and baseline corrected.

### Tryptophan fluorescence quenching assay

The binding interaction between the oligopyridylamides (ADH-1 and ADH-6) and mutant p53 DBD-derived aggregation domain (pR248W) was characterized using a previously published steady-state intrinsic tryptophan fluorescence quenching assay^91, 92^. Briefly, a 1 mM stock solution of pR248W was prepared in PBS. The peptide solution was then diluted to a final concentration of 5 μM and equilibrated at 20 °C for 5 min before commencing titration. Subsequently, the ligands (from a stock concentration of 10 mM in DMSO) were serially added to the peptide solution, which was stirred and allowed to equilibrate for 10 min before the intrinsic tryptophan fluorescence was measured from 300 to 420 nm (λ_ex_ = 280 nm) on a Cary Eclipse Fluorescence Spectrophotometer (Agilent, Santa Clara, CA). The background corrections in fluorescence changes because of dilution effects with the stepwise addition of ligands were made by buffer titrations. Curve fitting and the analysis of the data were performed using Prism 8.4.2 (GraphPad Software, Inc., La Jolla, CA, USA). All binding curves were fit to the one site-specific binding model to determine equilibrium dissociation constants (*K*_d_):

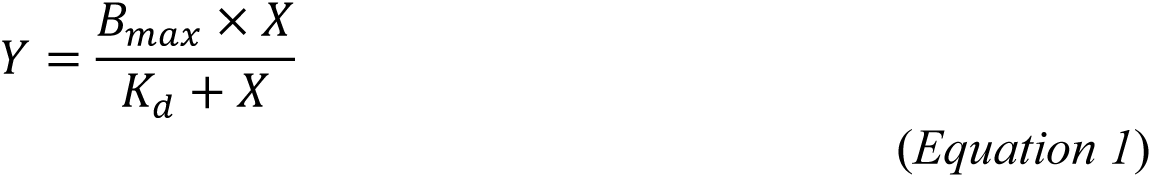

where *X* is the final concentration of the ligand, *Y* is the percentage increase in pR248W fluorescence quenching relative to the control fluorescence signal (i.e. pR248W alone), and *B*_max_ is the extrapolated maximum specific binding.

### Nuclear magnetic resonance (NMR) spectroscopy

NMR samples were prepared with the DBDs of WT and mutant R248W p53. The ^15^N-uniformly labeled proteins were expressed and purified as previously described^36^ by Giotto Biotech S.r.l. (Sesto Fiorentino, Italy) and supplied as 1 mg/mL solutions in 50 mM Tris, 200 mM KCl, 5 mM DTT (pH 7.5). After mild concentration by ultracentrifugation and spectroscopic inspection, the original buffer was replaced by dialysis, first against 50 mM KH_2_PO_4_/ K_2_HPO_4_, 150 mM KCl, 5 mM DTT (pH 6.8), and subsequently against H_2_O with 5 mM DTT which resulted in final pH values of 5.6–5.8. The protein concentrations were in the 40–50 μM range in Tris and phosphate buffers, and in the 19–24 μM range in water, with 4–5% D_2_O always present. Additions of a few μL of ADH-6 (from either 5 or 24 mM solutions in H_2_O, or 200 mM solution in DMSO), were performed to assess the protein-drug interaction.

NMR spectra were collected at 14.0 T (^1^H resonance at 600.13 MHz) on a Bruker Avance III NMR system equipped with triple resonance cryoprobe. All the spectra were acquired at 293.2 K. Two-dimensional ^15^N-^1^H HSQC experiments^93^ were executed using sensitivity-improved Echo/Antiecho-TPPI pure phase detection in F1, gradient-based coherence selection and flip-back pulse for solvent suppression^94–96^. To improve the resolution, the corresponding TROSY-based experiments^97, 98^ were also collected. All acquisitions were carried out over spectral widths of 40 ppm and 16 ppm in F1 and F2 dimensions, respectively, with 128 time-domain points in t1, 128–256 scans × 2048 points in t2 and 64 dummy scans to achieve steady state. The NMR data were processed with TOPSPIN version 4.0.2. Prior to Fourier transformation, linear prediction in t1 (up to 256 points) and zero filling were applied to yield a final data set of 2K*×*1K points. The chemical shift variations (Δ*δ*) of the protein peaks from ^15^N-^1^H HSQC spectra were analyzed in terms of cumulated chemical shift perturbation (CSP)^99^:

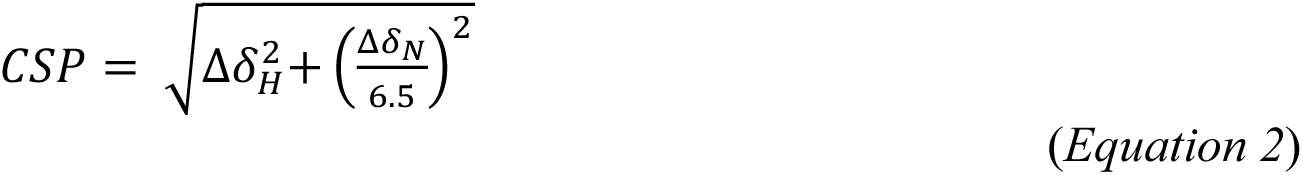

to properly account for both ^1^H (Δ*δH*) and ^15^N (Δ*δN*) frequency changes upon ligand interaction.

### Molecular docking simulations

High confidence homology models of the DBD of p53 were generated using the I-TASSER (Iterative Threading Assembly Refinement) Suite^100, 101^. The modeling resulted in two three-dimensional protein structures, WT and mutant R248W p53. The two structures were used in subsequent docking simulations, which were performed using the CANDOCK (Chemical Atomic Network-Based Hierarchical Flexible Docking) algorithm^35^. CANDOCK is a hierarchical, fragment-based docking with dynamics protocol that quickly and accurately generates the lowest energy binding pose for a particular compound–protein interaction using a knowledge-based statistical potential of mean force^35, 102^.

### Cell culture

All cell lines were purchased from American Type Culture Collection (ATCC). Prior to use, the cells were authenticated and tested for mycoplasma contamination by Charles River Laboratories (Margate, United Kingdom). Human pancreatic cancer MIA PaCa-2 (ATCC no. CRL-1420) and SW-1990 (ATCC no. CRL-2172) cells, human breast cancer MCF-7 (ATCC no. HTB-22) and HCC70 (ATCC no. CRL-2315) cells, and human brain cancer U-87 MG (ATTC no. HTB-14) cells were cultured in DMEM (Sigma) supplemented with 10% fetal bovine serum (FBS; GE Healthcare Life Sciences, Logan, UT), 4 mM L-glutamine, 1 mM sodium pyruvate and 1% penicillin/streptomycin (all from Sigma). Human ovarian cancer OVCAR-3 (ATCC no. HTB-161) and SKOV-3 (ATCC no. HTB-77) cells, and human colon cancer SW-480 (ATCC no. CCL-228) and COLO 320DM (ATCC no. CCL-220) cells, were cultured in RPMI 1640 (Sigma) supplemented with 10% FBS, 4 mM L-glutamine, 1 mM sodium pyruvate and 1% penicillin/streptomycin. Human lung cancer A549 (ATCC no. CCL-185) and NCI-H1770 (ATCC no. CRL-5893) cells were cultured in Ham′s Nutrient Mixture F12 (Sigma) supplemented with 10% FBS, 4 mM L-glutamine and 1% penicillin/streptomycin. All cells were cultured in 5% CO2 at 37 °C, and viability was monitored regularly using the Trypan Blue exclusion test on a TC20 automated cell counter (Bio-Rad, Hercules, CA). Once the cells reached ∼95% confluence, they were split (using 0.25% trypsin-EDTA; Sigma) into fractions and propagated or used in experiments.

### Thioflavin S (ThS) fluorescence imaging

The benzothiazole dye thioflavin S (ThS) was used to stain intracellular mutant p53 aggregates^38^. MIA PaCa-2 cells were seeded at a density of 2×10^4^ cells/well in 500 μL complete DMEM in 4-chambered 35 mm glass bottom Cellview cell culture dishes (Greiner Bio-One, Monroe, NC). After culturing for 24 h, the medium was replaced fresh medium containing 125 μM of ThS and incubated for 1 h. Thereafter the medium was removed, the cells were washed and fresh DMEM containing 10 µM ADH-1, ADH-6 or ReACp53 was added and incubated for 0.5–6 h. Finally, the medium was once again replaced with fresh DMEM and the cells were imaged on an Olympus Fluoview FV-1000 confocal laser scanning microscope, using a 63*×* Plan-Apo/1.3 NA oil immersion objective with DIC capability. Image analysis was done using the Fiji image processing software.

### Immunofluorescence

MIA PaCa-2 and MCF-7 cells were seeded at a density of 2×10^4^ cells/well in 500 μL complete DMEM in 4-chambered 35 mm glass bottom Cellview cell culture dishes and cultured for 24 h. Following treatment with vehicle or 2.5–10 µM ADH-1, ReACp53 or ADH-6 (unlabeled or FITC-labeled, ADH-6_FITC_) for 0.5 or 6 h, the cells were rinsed with ice-cold PBS, fixed with 4% paraformaldehyde, permeabilized with 0.5% Triton X-100, and blocked in 5% BSA. Subsequently, the cells were incubated with the anti-p53 antibody PAb 240 (5 μg/ml; Abcam, Cambridge, UK), overnight at 4 °C, which was followed by incubation with Alexa 549-labeled secondary antibody (1:800; Abcam) for 1 h at room temperature. Finally, the cells were imaged on an Olympus Fluoview FV-1000 confocal laser scanning microscope, and the images were analyzed using the Fiji image processing software.

### Plant cell studies

#### Plants and growth conditions

*Nicotiana benthamiana* were sown in soil and grown at 24 °C (morning) and 22 °C (night) in a growth chamber under an 8 h-light/16 h-dark cycle. After two weeks, the seedlings were transplanted into pots (one seedling per pot) and grown for a further 5–6 weeks before conducting *Agrobacterium tumfaciens* mediated transient assays.

#### Plasmid construction

WT and mutant R248W p53 DNA binding domains (DBDs) were cloned using the Gateway system (Thermo Fisher Scientific). Genes for WT and R248W p53 DBDs were respectively amplified from the vector pCMV-Neo-Bam carrying WT and R248W p53 constructs. Gene specific primers for the p53 DBDs were designed to incorporate the CACC tag at the 5’ end of both constructs. The following primers were used for amplifying p53 DBDs: p53 forward, 5’-**<u>cacc</u>**TCATCTTCTGTCCCTTCCCAGAAAACC-3’; p53 reverse, 5’-**tca**GGGCAGCTCGTGGTGAGGCTCCCCT-3’; and p53 reverse, 5’-GGGCAGCTCGTGGTGAGGCTCCCCT-3’ (note that the underlined CACC tag in lowercase is not a part of the p53 sequence). Phusion DNA Polymerase (New England Biolabs, Ipswich, MA) was used for the gene specific amplification of the p53 DBD constructs. The PCR amplified DNA constructs were subcloned in pENTR Directional TOPO (D-TOPO) (Thermo Fisher Scientific). The gateway binary vectors pEarley Gate 101 (35S promoter, C:YFP) and pEarley Gate 104 (35S promoter, N:YFP) were used as the destination vectors for the p53 DBD constructs. The WT p53 DBD was cloned with a YFP tag at the N terminus and an amino acid sequence, RVGAPTQLSCTKWC, at the C-terminal end that corresponds to the att site (5’-aagggtgggcgcgccgacccagctttcttgtacaaagtggTGC<u>TAG</u>-3’). The R248W p53 DBD, also cloned with a YFP tag at the N terminus, had a stop codon (<u>TAA</u>) at the C-terminal end. (Note: the nucleotides in lowercase are the att sites, while the uppercase nucleotides are part of the vector. The stop codon is underlined.) Since the entry and destination vectors had the same antibiotic selection marker (kanamycin), we followed the PCR amplification based (PAB) method^103^ to recombine the DNA fragment from the entry vector into the destination vector. The DNA fragment was amplified from the entry vector using M13 primers (M13 forward, 5’-GTAAAACGACGGCCAG-3’, and M13 reverse, 5’-CAGGAAACAGCTATGAC-3’) and was subsequently recombined with the destination vector using LR Clonase (Thermo Fisher Scientific). The plasmids were then transformed in *Agrobacterium tumfaciens* strain GV3101.

#### Agrobacterium-mediated transient expression

*N. benthamiana* leaves were infiltrated with *A. tumfaciens* carrying the desired constructs as described previously^104, 105^. Briefly, *A. tumfaciens* strain GV3101 carrying the WT and mutant R248W p53 DBDs were grown overnight at 28 °C in Lurai-bertani (LB) medium containing the appropriate antibiotics (100 µg/mL kanamycin, 50 µg/mL gentamycin, or 100 µg/mL of rifampicin). Cells were re-suspended in induction media (10 mM MES, pH 5.6, 10 mM MgCl2 and 200 µM acetosyringone) and incubated for 2 h at room temperature before infiltration (at a final OD of 1.0) of *N. benthamiana* leaves.

#### Confocal microscopy analysis

Intracellular localization of YFP:p53DBD^WT^ and YFP:p53DBD^R248W^ was observed at 72 h post infiltration (hpi). To determine the effects of ADH-1, ReACp53 and ADH-6 on p53 DBD aggregation, at 48 hpi the oligopyridylamides/peptide were diluted in 10 mM MgCl_2_ to a final concentration of 5 µM and then infiltrated into the *N. benthamiana* leaves for 24 h. The leaves from the infiltrated area were isolated and cells towards the abaxial side were imaged on a NIKON LV100 upright microscope and the images processed using the ECLIPSE LV software.

#### Protein extraction and SDS-PAGE

To verify that WT and mutant R248W p53 DBDs were expressed, total protein was extracted from the plant tissue as described previously^106^. Briefly, the leaf tissue was homogenized in 300 µL extraction buffer (20 mM Tris, pH 7.5, 150 mM NaCl, 1mM EDTA, 1% Triton X-100, 0.1% SDS, 5 mM DTT and 10*×* plant protease inhibitor cocktail (Sigma)), and the samples were centrifuged at 12,000 rpm for 20 min at 4 °C to remove insoluble debris. Thereafter, 5*×* SDS-PAGE loading dye was added to 100 µL of the collected supernatant (total protein) and the mixtures were heated for 10 min at 65 °C. The protein samples were then resolved on a 12% SDS-PAGE gel and subsequently transferred to a nitrocellulose membrane. Finally, the membrane was probed with 1:5000 dilution of anti-GFP antibody (1:400; Santa Cruz) to detect YFP-tagged p53 DBD^WT^ and p53 DBD^R248W^.

### Cell fractionation and western blot analysis

Cell fractionation and western blot were done as previously described^107, 108^. Briefly, MIA PaCa-2 cells were treated with 5 μM ADH-1, ReACp53 or ADH-6 for 24 h. Thereafter, the cells were washed with ice-cold PBS and lysed by using 100 μL lysis buffer (Bio-Rad). After confirming lysis by light microscopy, the cell lysate was centrifuged (16,000×g for 10 minutes at 4 °C) and the pellet (insoluble fraction) and supernatant (soluble fraction) were separated. The protein content of the two fractions was determined using the bicinchoninic acid (BCA) protein assay^109^. Samples (20 µg/lane) were electrophoresed on a 6–12% SDS polyacrylamide gel in a Mini-PROTEAN Tetra Cell (Bio-Rad) and subsequently transferred to a nitrocellulose membrane (Sartorius AG, Göttingen, Germany), which was blocked with 5% nonfat milk and then incubated with mouse anti-p53 antibody DO-7 (1:1000; Santa Cruz), or anti-*β*-actin antibody (1:2000; Santa Cruz), overnight at 4 °C. This was followed by incubation for 3 h at room temperature with horseradish peroxidase-conjugated mouse IgG antibody (1:10,000; Santa Cruz). Finally, the samples were visualized using Clarity Western ECL Substrate (Bio-Rad) and analyzed using a Bio-Rad Gel Doc XR+ Gel Documentation System.

### Transfection of SKOV-3 cells

SKOV-3 cells were plated at a density of 1*×*10^5^ cells/well in 2 mL complete medium in 6-well plates and cultured for 48 h. For each well of cells to be transfected, 2.5 μg pCMV-Neo-Bam p53 R248W vector (plasmid 1647; Addgene, Watertown, MA) containing the *amp*^r^ gene was diluted in 500 μL Opti-MEM Reduced Serum Medium in an Eppendorf tube, to which 10 μL Lipofectamine LTX Reagent was added, mixed gently and incubated for 30 min at room temperature. The medium in the wells was then replaced with the Opti-MEM Reduced Serum Medium containing the DNA– Lipofectamine LTX Reagent complexes and the cells were incubated in 5% CO2 at 37 °C for 24 h. One well of transfected cells was harvested and blotted for mutant p53 using standard western blot conditions, with un-transfected cells used as a control. Subsequently, the other transfected cells were harvested and plated in a T25 flask (Corning). After culturing until ∼95% confluence was reached, the cells were selected with 500 μg/mL G418 sulfate-supplemented medium, and maintained thereafter in medium containing 100 μg/mL G418 sulfate.

To verify that the transfection was successful, the pCMV-Neo-Bam p53 R248W vector containing the *amp*^r^ gene was purified from the transfected SKOV-3 cells using the Qiaprep Miniprep Kit (Qiagen, Hilden, Germany) according to the manufacturer’s instructions. The miniprep-purified DNA sample was subsequently quantified using a NanoDrop spectrophotometer. PCR was carried out to amplify the *amp*^r^ gene using the following primers: forward (5’-AGATTATCAAAAAGGATCTTCACCT-3’) and reverse (5’-CCTCGTGATACGCCTATTTTTATAG-3’) (Integrated DNA Technologies, Leuven, Belgium). Each PCR reaction was prepared in 25 μL volume containing: 0.5 μL of each primer (0.2 μM final primer concentration), 5 μL DNA and 6.5 μL nuclease-free water and 12.5 μL PCR MasterMix (Thermo Fisher Scientific). The conditions for each PCR reaction were: initial denaturation at 95 °C for 30 s, 35 cycles of denaturation at 95 °C for 30 s, annealing at 50 °C for 30 s, and extension at 68 °C for 1 min. As a negative control, a PCR reaction was carried out with the same reagents sans DNA. The amplified products were then resolved by electrophoresis using a 1.2% E-gel containing a SYBR Safe stain (Thermo Fisher Scientific), which was visualized on an E-Gel Imager Blue-Light Base (Thermo Fisher Scientific). The presence of the successfully amplified *amp*^r^ PCR product was confirmed using a standard 500 bp ladder.

### Cell viability/toxicity assays

Cell viability/toxicity was measured using two complementary assays as previously described^73^: i) CellTiter 96 AQueous One Solution (MTS) assay, which measures reduction of the tetrazolium compound MTS (3-(4,5-dimethylthiazol-2-yl)-5-(3-carboxymethoxyphenyl)-2-(4-sulfophenyl)-2H-tetrazolium, inner salt) to soluble formazan, by mitochondrial NAD(P)H-dependent dehydrogenase enzymes, in living cells^110, 111^; and ii) Dead Cell Apoptosis assay, in which Alexa 488-conjugated annexin V is used a sensitive probe for detecting exposed phosphatidylserine in apoptotic cells^112^, and red-fluorescent PI, a membrane impermeant nucleic acid binding dye, assesses plasma membrane integrity and distinguishes between apoptosis and necrosis^113^.

For the MTS assay, cells were seeded at a density of 2×10^4^ cells/well in 100 μL complete medium in standard 96-well plates. After culturing for 24 h, the medium was replaced with serum-free medium containing the oligopyridylamides at the desired concentrations, and incubated for the indicated durations, at 37 °C. Thereafter, the incubation medium was replaced with fresh complete medium, and 20 μL MTS reagent was added to each well and incubated for 4 h at 37 °C. Finally, absorbance of the soluble formazan product (*λ* = 490 nm) of MTS reduction was measured on a Synergy H1MF Multi-Mode microplate-reader (BioTek, Winooski, VT), with a reference wavelength of 650 nm to subtract background. Wells treated with vehicle were used as control, and wells with medium alone served as a blank. MTS reduction was determined from the ratio of the absorbance of the treated wells to the control wells.

For the Dead Cell Apoptosis assay, MIA PaCa-2 cells were treated with 5 μM ADH-1, ADH-6 or ReACp53 for 24 h at 37 °C. Subsequently, the cells were washed with ice-cold PBS, harvested by trypsinization, centrifuged and re-suspended in 1×annexin-binding buffer (10 mM HEPES, 140 mM NaCl, 2.5 mM CaCl_2_, pH 7.4) to a density of ∼1×10^6^ cells/mL. The cells were then stained with 5 µL Alexa 488-conjugated annexin V and 0.1 µg PI per 100 µL of cell suspension for 15 min at room temperature. Immediately afterwards, fluorescence was measured using flow cytometry on a BD FACSAria III cell sorter (BD Biosciences, San Jose, CA), and the fractions of live (annexin V-/PI-), early and late apoptotic (annexin V+/PI- and annexin V+/PI+, respectively), and necrotic (annexin V-/PI+) cells were determined using the BD FACSDiva software.

### Cell cycle analysis

MIA PaCa-2 cells were seeded at a density of 1×10^6^ cells/well in 1 mL complete DMEM. After culturing for 24 h, the medium was replaced with serum-free DMEM containing 5 µM ADH-1, ReACp53 or ADH-6, and the cells were incubated for 6 h. Subsequently, the cells were washed with PBS, harvested by trypsinization and centrifuged (800*×*g for 5 min). The supernatant was discarded, and the cells were washed with PBS and fixed in 70% cold ethanol overnight at 4 °C. The cells were then washed with PBS, filtered through a nylon sieve, and centrifuged again (800*×*g for 5 min), and the supernatant was discarded. The cells were stained with Nuclear Green CCS1 (Abcam) for 30 min at 37 °C and analyzed using flow cytometry (BD FACSAria III cell sorter)^114, 115^.

### RNA sequencing (RNA-Seq)

#### RNA isolation and purification

Total RNA was extracted from MIA PaCa-2 cells (1×10^6^ cells/ well) treated with vehicle (control (C), *n* = 3) or oligopyridylamides (5 µM ADH-1 or ADH-6, *n* = 3) using a combination of both TriZol (Thermo Fisher Scientific) and RNAeasy Mini Kit (Qiagen) with modification of manufacturers’ protocol. Prior to extraction, cells were washed once with 1 mL PBS, suspended in 1 mL TriZol LS reagent and transferred to 1.5 mL Lobind tubes (Eppendorf) along with 0.2 mL chloroform. The TriZol lysate and chloroform mix was vortexed and centrifuged (12,000 rpm at 4 °C) for 15 min, forming three distinctive phases of DNA (lower phase), protein (middle phase) and RNA (upper phase). The upper aqueous phase (∼400 mL of RNA) was transferred to a new 1.5 mL tube where an equivalent volume of 70% ethanol was added and mixed. The mixture was further purified using an RNAeasy mini spin column (Qiagen) according to the manufacturer’s protocol. RNA concentrations were determined using a NanoDrop 2000 spectrophotometer (Thermo Fisher Scientific, Waltham, MA).

#### RNA-Seq library preparation, sequencing and processing

Total RNA quality was estimated based on the RNA integrity number (RIN) using a BioAnalyzer 2100 (Agilent, Santa Clara, CA). RNA samples with a RIN > 8 were used for library preparation. RNA-Seq libraries were prepared using an Illumina TruSeq Stranded mRNA prep kit according to the manufacturer’s protocol. Samples were barcoded, multiplexed and sequenced (100 bp pair-end) using a NextSeq 550 System (Illumina, San Diego, CA). DESeq2 computational pipeline^116^ was used to estimate the raw count reads aligned to the reference genome. Human reference genome (GRCh38/hg38) from UC Santa Cruz Genome Browser (https://genome.ucsc.edu/) was utilized as a reference genome^117^. Computations were run on a Linux based command system on the NYU Abu Dhabi High Performance Computing (HPC) server platform Dalma (https://wikis.nyu.edu/display/ADRC/Cluster+-+Dalma).

#### Bioinformatic and computational analysis

Correlation (i.e. PCA, distance dendrogram) and expression (i.e. heatmaps, boxplots) analyses were generated either via RNA-Seq START (Shiny Transcriptome Analysis Resource Tool) application (NYU Abu Dhabi Center of Genomic and Systems Biology (NYUAD-CGSB) Bioinformatics Online Analysis and Visualization Portal (http://tsar.abudhabi.nyu.edu/))118 or JMP genomics software (http://www.jmp.com/software/genomics/). For gene ontology (GO) analysis, differentially expressed genes (DEGs), based on corrected statistical significance (*P*-adj < 0.05) lists, were submitted to Database for Annotation, Visualization Integrated Discovery (DAVID) bioinformatics tool (https://david.ncifcrf.gov/home.jsp/)119, thereby determining characteristics under the category of biological processes. Enrichment term values were based on statistical significance of *P* < 0.05, which were normalized to −log10 (*P-value*). GO analysis charts were prepared using Excel.

Ingenuity pathway analysis (IPA) was used to identify transcriptional regulators (TRs) responsible for transcriptional dysregulation in ADH-6 vs ADH-1 and ADH-6 vs control comparisons. For each TR, the q-value and z-score, along with the number of genes, was obtained. The number of genes that showed a pattern consistent with a TR’s activated or repressed status were plotted along with the z-score and log10-q-value in a 2D plot generated in R. Gene set enrichment analysis (GSEA) was performed using the variance transformed counts obtained for DESeq2 analysis against the Molecular Signatures Database (MSigDB)^120, 121^ for identification of hallmark signatures. The enrichment plots, along with normalized enrichment scores and q-values, were obtained from GSEA and are shown on each plot.

### Quantitative proteomics

#### Protein extraction and preparation of digests

MIA PaCa-2 cells (1×10^6^ cells/ well) were treated with vehicle or 5 µM ADH-1, ReACp53 or ADH-6 for 16 h, harvested, centrifuged and lysed by adding five cell-pellet volumes of lysis buffer (100 μL lysis buffer for every 20 μL cell pellet). The cell lysate was centrifuged (16,000×g for 10 minutes at 4 °C) and the supernatant was transferred to a 1.5 mL LoBind tube. The protein content of the sample was determined using the BCA protein assay. An aliquot (100 µg of protein per treatment) was diluted to a final volume of 100 µl with 100 mM TEAB buffer. Thereafter, the proteins were reduced (10 mM DTT for 1 h at 55 °C), S-alkylated (25 mM IAA, 30 min in the dark) and precipitated by adding six volumes (∼600 µL) of pre-chilled (−20 °C) acetone for 16 h. The protein pellet collected by centrifugation (8000×g for 10 min at 4 °C), washed with pre-chilled acetone, air dried and re-suspended in 100 µL TEAB (100 mM). The proteins were digested with 1:40 (w/w) MS-grade Pierce Trypsin/Lys-C Protease Mix for 24 h at 37 °C^122^.

#### Phosphopeptide enrichment using titanium dioxide (TiO_2_)

Protein digests were dried using SpeedVac and re-suspended in 1 mL loading buffer (80% acetonitrile (ACN), 5% TFA and 1 M glycolic acid)^123^. TiO2 beads (0.6 mg per 100 µg peptide solution) were added, mixed well and placed on a shaker for 15 min at room temperature. The solution was then table centrifuged to pellet the beads, which were subsequently mixed with 100 µL loading buffer, transferred to LoBind tube and table centrifuged. Thereafter, the beads were washed first with 100 µL wash buffer 1 (80% ACN, 1 % TFA) followed by 100 µL wash buffer 2 (20% ACN, 0.2% TFA). The bead pellet, collected by table centrifugation, was dried using SpeedVac, mixed with 50 µL alkaline solution (28% NH3 in H_2_O, Thermo Fisher Scientific) and incubated for 15 min at room temperature for phosphopeptide elution. The solution was table centrifuged and the eluate was passed over a C18 Tip to recover the phosphopeptides. Finally, the C18 tip was washed with 20 µL of 30% ACN solution, collected in the same tube, and dried using SpeedVac.

#### Tandem mass tag (TMT) labeling

The phosphopeptide enriched digests were re-suspended in 100 µL TEAB (100 mM). A labeling kit was used according to the manufacturer’s protocol for TMT labeling^124^. Briefly, TMT10plex amine-reactive reagents (0.8 mg per vial) were re-suspended in 41 μL anhydrous ACN and all 41 μL of each reagent was added to each sample and mixed briefly. Reactions were allowed to proceed at room temperature for 1 h, which were then quenched by addition of 8 μL of 5% hydroxylamine for 15 min. Subsequently, the samples were combined in equal amount (1:1), transferred to a LoBind tube, and dried using SpeedVac. The TMT-labeled peptide mixtures were desalted on an Isolute MFC18 cartridge (100 mg/3 mL; Biotage, Charlotte, NC). Eluted peptides were fractionated into 5 equal aliquots, dried and resuspended in 20 µL of 0.1% formic acid solution prior to liquid chromatography with tandem mass spectrometry (LC*-*MS*-*MS).

#### LC-MS/MS of TMT labeled peptides using MS_2_

Liquid chromatography was performed on a fully automated EASY-nLC 1200 (Thermo Fisher Scientific) fitted with a C18 column (PepMap RSLC, 75 μm inner diameter, 15 cm length; Thermo Fisher Scientific), which was kept at a constant temperature of 40 °C. Mobile phases consisted of 0.1% formic acid for solvent A and 0.1% formic acid in ACN for solvent B. Samples were loaded in solvent A, and a linear gradient was set as follows: 0–5% B for 5 min, followed by a gradient up to 30% B in 50 min and to 60% B in 65 min. A 10-min wash at 95% B was used to prevent carryover, and a 15-min equilibration with 0% B completed the gradient. The liquid chromatography system was coupled to a Q Exactive HF Hybrid Quadrupole-Orbitrap (Thermo Fisher Scientific) equipped with an Easy-Spray Ion Source and operated in positive ion mode. The spray voltage was set to 1.7 kV, S-lens RF level to 35 and ion transfer tube to 275 °C. The full scans were acquired in the Orbitrap mass analyzer that covered an *m/z* range of 350–1500 at a resolution of 120,000. The automatic gain control (AGC) target was set to 3E6 and maximum ion time to 50 ms. The MS^2^ analysis was performed under data-dependent mode to fragment the top 15 most intense precursors using higher-energy collisional dissociation (HCD) fragmentation. The MS^2^ parameters were set as follows: resolution, 60 000; AGC target, 1E5; minimum AGC target, 8.0E3; intensity threshold, 8.0E4; maximum ion time, 100 ms; isolation width, 1.2 *m/z*; precursor charge state, 2–7; peptide match, preferred; dynamic exclusion, 30 s; and fixed first mass, 100 *m/z*. The normalized collision energy for HCD was set to 32%.

#### Relative protein quantification using TMT

Relative protein abundances in the different treatment groups were determined using PEAKS studio proteomics software (version 10.0, build 20190129) with the following settings^125^: enzyme, trypsin; instrument, Orbitrap; fragmentation, HCD; acquisition, data-dependent acquisition (DDA) without merging the scans; and precursor and fragment mass tolerance, 20 ppm and 0.5 Da, respectively. The correct precursor was detected using mass only. Peptide identifications were performed within PEAKS using the UniProtKB_TrEMBL database with human taxonomy. PEAKS PTM search tool was used to search for peptides with fixed (carbamidomethylation and TMT10plex) and variable (phosphorylation (STY), oxidation (M) and deamidation (NQ)) modifications. SPIDER search tool was used for finding novel peptides that were homologous to peptides in the protein database. The maximum number of variable posttranslational modifications per peptide was 5, and the number of *de novo* dependencies was 18. The *de novo* score threshold for SPIDER was 15 and Peptide hit score threshold 30. All search tools were utilized in the PEAKS studio proteomics software. False discovery rate (FDR) threshold was set to 1%. Reporter ion quantification with TMT-10plex (CID/HCD) was used. PEAKS was allowed to autodetect the reference sample and automatically align the sample runs. Reporter ion intensity ≥ 1E4, significance ≥ 16, fold change ≥ 1 and 2 unique peptides per protein were used to filter the quantified protein lists.

Further subgroup analysis of differentially expressed phosphoproteins for ADH-6 vs ADH-1 and ReACp53 vs ADH-1 was done by t-test for each group, and only those proteins that differed significantly (*P* < 0.05) were used for further analysis. Fold change of protein expression was obtained from the averaged intensity value for each protein in ADH-6/ADH-1 or ReACp53/ADH-1. The raw data used for this analysis is provided in Supplementary Data File.

#### Principal component and pathway analyses

PCA was performed to analyze the main source of variation within and amongst the different treatment groups. Raw counts were used for differently abundant proteins (Supplementary Data File) and were plotted using the ClustVis application^126^. GSEA was performed for the “compute overlap function” using MSigDB to identify hallmark signatures. For this analysis, the protein accession IDs were converted to gene symbols using David Gene ID conversion tool^127^. Significance was assessed by FRD or q-value of the hypergeometric *P*-value corrected for multiple hypothesis testing using the Benjamin and Hochberg procedure^128^. The data was ordered based on −log10 q-values.

#### Plots and network plots

Heatmaps showing differentially expressed phosphoproteins were generated in R using the Pearson correlation analysis^129^. GO-Chord plot showing the pathways and the corresponding proteins that are part of each pathway/gene signature was also generated in R. Biological roles of the phosphoproteins in cancer were further inferred from published data (Supplementary Table 1). Upregulated phosphoproteins were depicted using the protein-protein interaction network using STRING^130^.

### *In vivo* tumor inhibition studies

All animal experiments were approved by the NYU Abu Dhabi Institutional Animal Care and Use Committee (NYUAD-IACUC; Protocol No. 18-0001), and were carried out in accordance with the Guide for Care and Use of Laboratory Animals^131^. Athymic nude NU/J mice (Foxn1^nu^; The Jackson Laboratory, Bar Harbor, ME) were bred in-house by the NYU Abu Dhabi Vivarium Facility in a 12 h light/dark schedule.

For pharmacokinetics, tumor-bearing mice (tumor volume ∼25 mm^3^) were randomly assigned to saline and ADH-6 treatment groups. A single dose of saline or ADH-6 (15 mg kg^-1^) was administered via intraperitoneal (i.p.) injection. Mice (*n* = 3 per time point) were sacrificed at 0.5, 1, 2, 4, 8, 16, 24, 48 and 72 h following injection. The blood was collected via terminal cardiac puncture using K3-EDTA as an anticoagulant under isoflurane anesthesia and processed for plasma by centrifugation (1500×*g* for 5 min). Plasma and tissues were placed in cryopreservation vials, preserved by snap freezing using liquid N2, and stored at −80 °C until analysis by LC–MS/MS, which was performed according to a previously published protocol^132^. Briefly, ADH-6 stock solutions (1.0 mg/mL) were prepared in acetonitrile (ACN). The matrix for the standard curve and quality controls (QC) consisted of control mouse plasma for all plasma samples, or control tissue homogenate for the tissue being analyzed. ADH-6 was extracted from 50 μL standard, QC, or unknown sample by protein precipitation with 200 μL ACN/0.1% formic acid containing 20 ng/mL ADH-6 internal standard.^5^ Samples were vortexed for 5 min, then centrifuged at 5000×*g* for 10 min at 4 °C. Subsequently, 150 µL supernatant was transferred to a clean 1.5 mL tube, lyophilized under N2, and reconstituted in 60 μL ACN/0.1% formic acid. 50 µL of this was transferred to a salinized glass 96-well plate insert containing 50 μL ddH_2_O and 10 μL of the sample was injected for analysis by LC–MS/MS.

For the single mutant p53-harboring xenograft inhibition studies, 5×10^5^ viable MIA PaCa-2 cancer cells were injected subcutaneously into the right flank of each mouse at age 6–8 weeks. For the dual, WT and mutant p53-bearing, xenograft model, 5×10^5^ viable MIA PaCa-2 and MCF-7 cancer cells were injected into the right and left flanks, respectively, of each mouse at age 6–8 weeks. Mice were assessed daily for overt signs of toxicity. Tumor volume was measured via high-precision calipers (Thermo Fisher Scientific) using the following formula:

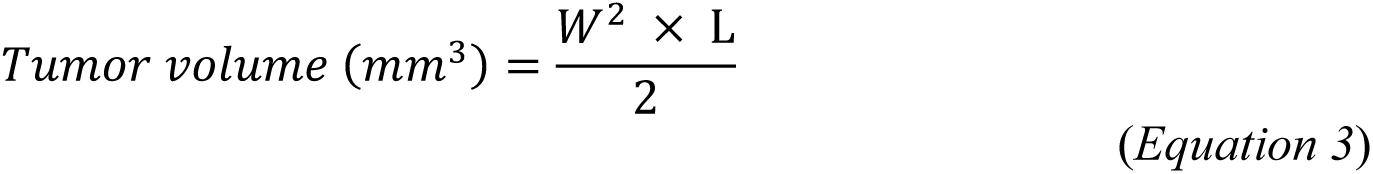

where W and L are tumor width and length in mm, respectively^133^. Mice were euthanized once tumor volume approached burden defined by NYUAD-IACUC.

Once the tumor volume reached ∼25 mm^3^, the mice were randomized into 4 treatment groups (*n* = 8 per group), which were injected intraperitoneally with: saline or 156.4 µM ADH-1, ReACp53 or ADH-6, for the single xenograft model; vehicle (0.02% DMSO) or 716.4 µM ReACp53 or ADH-6, for the dual xenograft model. Injections were done every 2 days for a total of 12 doses, with the first day of treatment defined as day 0. Body weight and tumor volume were recorded for the duration of treatment, and survival (*n* = 4 per group) was monitored for a total of 30 days. At the end of treatment, 4 mice per treatment group were sacrificed and the tumor tissues were isolated to determine the tumor mass.

Isolated tumors and vital organs were formalin-fixed and paraffin-embedded and sectioned into 7-μm slices. For hematoxylin and eosin (H&E) analysis, the tissue sections were dewaxed and stained using standard procedures as previously described^73^. For immunohistochemistry (IHC) analysis, the samples were prepared according to a published protocol^14^. Briefly, the tissue sections were treated by heat-induced epitope retrieval in 10 mM citrate buffer (pH 6.0) for antigen recovery, blocked with 8% BSA, then incubated overnight at 4 °C in primary antibodies (1:400 DO-7 or 5 μg/mL PAb 240). This was followed by sequential 45 min incubations at room temperature in biotinylated secondary antibody and streptavidin-horseradish peroxidase (Abcam). Finally, the signal was visualized upon incubation with the 3,3’-diaminobenzidine (DAB; Thermo Fisher Scientific) substrate. The tissue sections were imaged on a NIKON LV100 upright microscope and processed using the ECLIPSE LV software.

### Statistics and reproducibility

For *in vitro* studies, investigators were blinded for all parts of the experiments (treatment, data acquisition and data analysis), and a different investigator carried out each part. For *in vivo* studies, power calculation was used to select sample sizes from the NYU Abu Dhabi Institutional Animal Care and Use Committee (NYUAD-IACUC) Protocol (Protocol No. 18-0001). Confidence intervals in this work represent the standard deviation across at least three biological replicates (i.e. n *≥* 3). Statistical analysis was performed using Prism 8.4.2 (GraphPad Software, Inc.). Statistical significance between two groups was assessed by an unpaired *t*-test, and among three or more groups by two-way analysis of variance (ANOVA) followed by Tukey’s *post hoc* test. *P* < 0.05 was considered to be statistically significant.

## SUPPLEMENTARY INFORMATION

Supplementary Sections, Figures and Data are available online or from the authors.

## ACKNOWLEDGEMENTS

The authors thank Khulood Alawadi (Assistant Director, Research Visualization, Design, and Manufacturing, NYU Abu Dhabi) for preparing the graphic illustrations. The authors also thank the NYU Abu Dhabi Center for Genomics and Systems Biology (NYUAD-CGSB) for use of their BD FACSAria III for flow cytometry measurements and Illumina NextSeq 550 System for RNA-Seq. Confocal fluorescence imaging, NMR, quantitative proteomics and TEM experiments were carried out using the Core Technology Platforms (CTP) resources at NYU Abu Dhabi. This work was supported by funding from NYU Abu Dhabi and an ADEK Award for Research Excellence grant (AARE17-089) to M. Magzoub. Additional support was provided by funding from NYU to A.D. Hamilton and the University of Denver to S. Kumar.

## AUTHOR CONTRIBUTIONS

M.M. conceptualized and designed the project in consultation with A.D.H. and S. Kumar. M.M., A.D.H. and S. Kumar supervised the project and provided funding. S. Kumar designed the oligopyridylamides, and S. Kumar and D.M. synthesized the compounds. M.M. and L.P. designed the aggregation, intracellular imaging, cell viability/toxicity, cell cycle distribution and related experiments, and L.P., L.K., S.H., M.K., R.P. and S. Karapetyan conducted these experiments. M.M. and L.P. designed the *in vivo* tumor studies, and L.P. conducted these studies. M.A.-S., I.C., L.K., L.P. and M.M. planned the RNA-Seq experiments, M.A.-S., I.C., L.K. and L.P. conducted these experiments, and M.A.-S., I.C., L.K. and A.J.A. performed the analysis. A.J.A., L.A., L.P. and M.M. planned the quantitative proteomics experiments, L.A. and L.P. conducted these experiments, and A.J.A., M.A. and L.A. performed the analysis. G.E. designed the NMR experiments, G.E., Y.H. and L.P. conducted these experiments, and G.E. and Y.H. performed the analysis. Z.F. and R.S. performed the molecular docking simulations. A.J.A. and M.A. planned the plant experiments, and M.A. and L.P. conducted these experiments. S. Kumar designed the oligomerization and peptide binding experiments, and J.A. conducted these experiments. S. Kumar and L.P. conducted the CD experiments. M.M. and L.P. wrote the manuscript with contributions from S. Kumar, A.J.A., M.A., M.A.-S., L.K., I.C., S.H., G.E., L.A., Z.F. and R.S. All authors subsequently reviewed and edited the manuscript.

