## Supplementary Information for "Protein mimetic amyloid inhibitor potently abrogates cancer-associated mutant p53 aggregation and restores tumor suppressor function"

*Short Title*: Protein mimetic inhibitor of mutant p53 aggregation

*Key Words*: α-helix mimetics, amyloid, apoptosis, cancer therapeutics, cell cycle arrest, DNA-binding domain, oligopyridylamides, mutant p53, pancreatic cancer, protein aggregation, tumor suppressor

### TABLE OF CONTENTS

**SECTION 1. Supplementary amyloid aggregation data**

Supplementary Figure 1. Effects of oligopyridylamides on amyloid formation of aggregation prone region of p53 DBD

**SECTION 2. Supplementary p53 DBD–ADH-6 interaction data**

Supplementary Figure 2. Computational modeling of p53 DBD–ADH-7 interaction strength
Supplementary Figure 3. $^{15}N$-$^{1}H$ HSQC spectrum of WT p53 DBD in Tris buffer
Supplementary Figure 4. $^{15}N$-$^{1}H$ HSQC spectra of WT and mutant p53 DBDs in phosphate buffer
Supplementary Figure 5. $^{15}N$-$^{1}H$ HSQC spectra of WT and p53 DBDs in $H_2O/D_2O$ (96/4)
Supplementary Figure 6. Chemical Shift Perturbation (CSP) values from $^{15}N$-$^{1}H$ HSQC spectra of WT and mutant p53 DBDs in the presence of ADH-6 at the indicated protein-to-ligand ratios
Supplementary Figure 7. $^{15}N$-$^{1}H$ HSQC maps of WT and mutant p53 DBDs in $H_2O/D_2O$ (96/4)

**SECTION 3. Supplementary intracellular localization and cell viability/toxicity data**

Supplementary Figure 8. Effects of ADH-1 and ReACp53 on cytosolic mutant p53 aggregates in MIA PaCa-2 cells
Supplementary Figure 9. ADH-6 reduces puncta in plant cells expressing mutant, but not WT, p53 DBD
Supplementary Figure 10. Effects of the oligopyridylamides on cancer cells harboring WT and mutant (R248W) p53

**SECTION 4. Supplementary transcriptome and proteome analysis**

Supplementary Figure 11. Determination of best condition for differential gene expression analysis
Supplementary Figure 12. Identification of transcriptional regulators of dysregulated genes in oligopyridylamide-treated MIA PaCa-2 cells
Supplementary Figure 13. Phosphoproteome analysis
Supplementary Figure 14. A Simplified model of p53 mediated regulation of DNA replication/ repair and cell cycle progression/proliferation
Supplementary Table 1. Biological roles of downregulated/upregulated phosphoproteins in DNA repair/replication and cycle progression/proliferation

**SECTION 5. Supplementary *in vivo* tumor reduction data**

Supplementary Figure 15. Effect of ADH-6 on xenografts bearing aggregation-prone mutant p53 *in vivo*
Supplementary Figure 16. Histological analysis of vital organs following treatment with lower doses of ADH-6
Supplementary Figure 17. Histological analysis of vital organs following treatment with higher doses of ADH-6

**SECTION 6. Synthesis and characterization of ADH-6**

Supplementary Figure 18. $^{1}H$ NMR of ADH-6-NCS
Supplementary Figure 19. $^{1}H$ NMR of ADH-$6_F$ (ADH-$6_{FITC}$)

### SECTION 1. Supplementary amyloid aggregation data

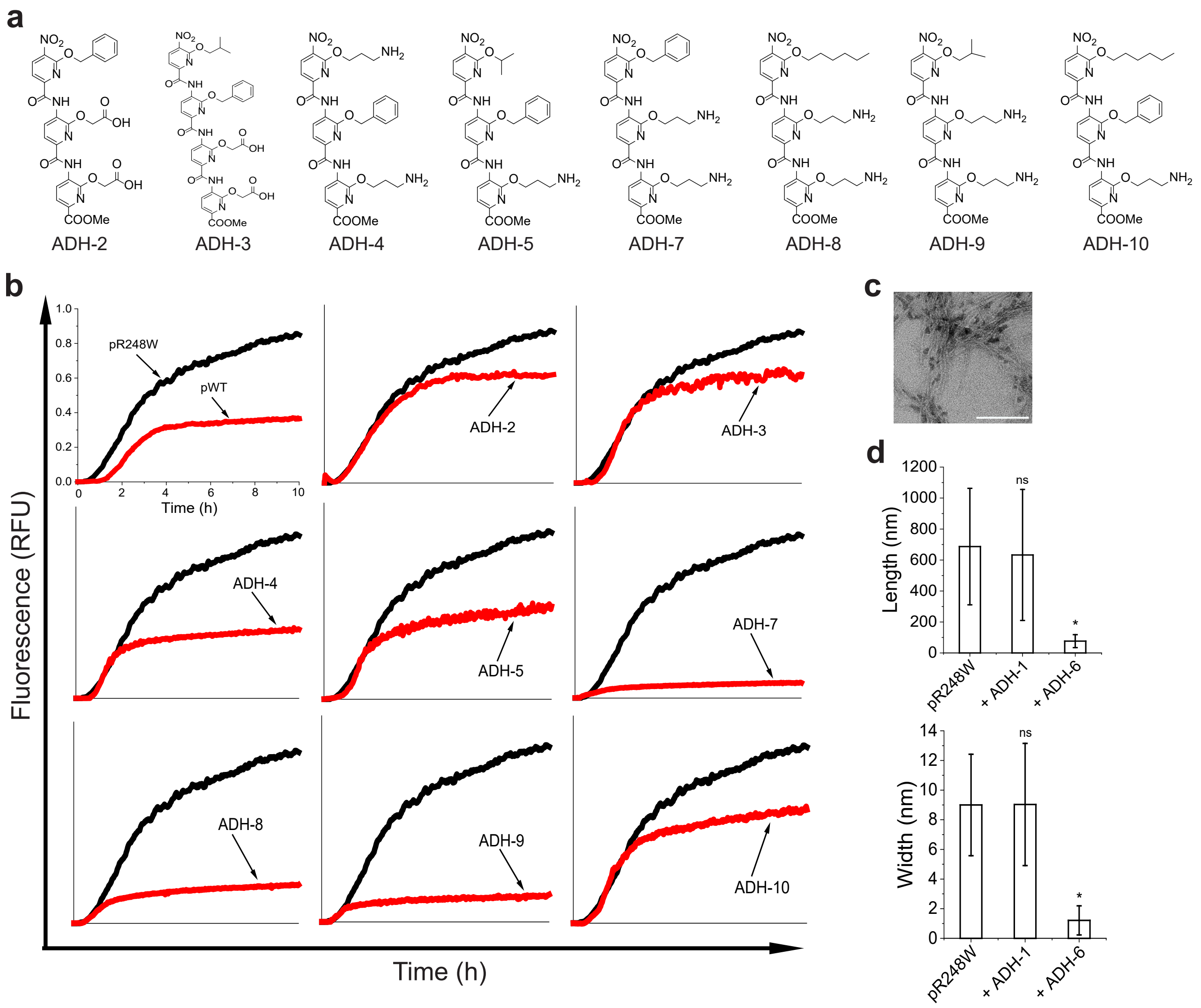


**Supplementary Figure 1. Effects of oligopyridylamides on amyloid formation of aggregation-prone region of p53 DBD. (a)** Chemical structures of the oligopyridylamides used in this study (except ADH-1 and ADH-6, which are shown in Figure 1d). (**b**) Effects of the oligopyridylamides on pR248W amyloid formation. Kinetic profiles for aggregation of 25 µM pR248W vs pWT, and for 25 µM pR248W in the absence or presence of an equimolar amount of the oligopyridylamides co-mixed at the start of the reaction. Aggregation profiles were acquired by measuring the fluorescence of the thioflavin T (ThT) reporter ($\lambda_{ex/em}$ = 440/480 nm) at 5-min intervals at 37 °C (*n* = 4). (**c**) Representative transmission electron microscopy (TEM) image for aggregation of 25 µM pR248W in the presence of an equimolar amount of ADH-6 added during the growth phase of (i.e 5 h after the start of the reaction). TEM image was acquired at 10 h after the start of the aggregation reaction. Scale bar = 500 nm. (**d**). Quantification of fibril lengths and widths for 25 µM pR248W in the absence or presence of an equimolar amount of ADH-1 or ADH-6 added during the growth phase (i.e. 5 h after the start of the reaction; representative TEM images shown in Figure 1f). For quantification, 3–5 different fields of view were used (*n* = 3). *$P$ < 0.05 or non-significant (ns, $P$ > 0.05) compared pR248W alone.

### SECTION 2. Supplementary p53 DBD–ADH-6 interaction data

#### 2.1. Supplementary molecular docking simulations

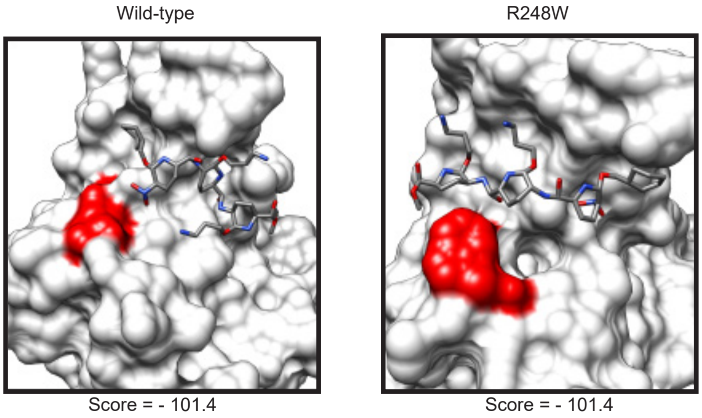


**Supplementary Figure 2. Computational modeling of p53 DBD–ADH-7 interaction strength.** (**a**,**b**) Top binding modes and the resulting scores for ADH-7 bound to the modelled structures of WT (*left panel*) and mutant R248W (*right panel*) p53 DBDs. For each binding mode, the DBD is depicted as a white surface, the residue of interest (248) is highlighted in red, and the corresponding compounds and peptides colored by element. The scores are based upon the CANDOCK knowledge-based scoring potential, where lower scores correspond to a more favorable binding.

#### 2.2. Supplementary NMR data

The initial $^{15}N$-$^{1}H$ HSQC NMR spectra of p53 DBD obtained in Tris buffer showed several differences compared to the maps reported by Rasquinha *et al.*[1] Besides numerous chemical shift displacements, the resonance intensities appeared weak and smeared despite the sample concentration not being particularly low (50 μM) (Supplementary Figure 3). As was previously reported, studying the NMR spectrum of p53 DBD is challenging due to the inherent instability of the domain[1,2], which is responsible for the conformational heterogeneity over intermediate timescales that leads to substantial line broadening of some signals.

In order to reproduce the spectral quality previously obtained[1], the protein sample was dialyzed to replace the Tris buffer with phosphate buffer. The quality of the HSQC contour plots improved, as did the comparability with the previously published spectra (Supplementary Figure 4). Most of the backbone amide assignments illustrated by Rasquinha et al.[1] could be transferred to our data. This also allowed us to recognize the presence of several new peaks, mostly clustered in the region of unstructured peptides, arising from a partially unfolded species. The onset of this species was ascribed to the effect of ultracentrifugation and buffer exchange manipulations on a protein of limited stability. Since the NMR pattern of the folded form was observed to be stable for a reasonable length of time (at least 15 days) for both the WT and R248W mutant species, notwithstanding the presence of a partially unfolded conformer, we decided to proceed to the titration with the designed oligopyridylamide ligand, namely ADH-6. However, we could not observe any effect on either protein variant due to the systematic precipitation of the oligopyridylamide after its addition to the protein samples. Regardless of the concentration of the aqueous titrant we employed to reach protein-to-ligand ratios between 1:2 and 1:10, no effect could be detected in the HSQC spectra, but the sedimentation at the bottom of the NMR tube of a white powder that had to be the added compound given the constancy of the protein spectra. The same result was obtained after adding a concentrated ADH-6 solution in DMSO to the R248W p53 DBD mutant, to reach a protein-to-ligand nominal ratio of the order of 1:100. It is worth noting that the protein did not exhibit any relevant instability effect due to 1 μL of DMSO even after 8 days.

To overcome the incompatibility of ADH-6 with p53 DBD in saline, the protein samples were further dialyzed against pure water containing only 5 mM DTT (necessary to preserve the reduced state of the 7 free Cys residues in the p53 DBD domain). Although the pH was lowered by 1 unit (for R248W p53 DBD) or 1.2 units (for WT p53 DBD) with respect to the initial value (6.8), the spectra of both proteins still showed good similarity to the literature[1]. Additionally, the partially unfolded conformers formed after the first buffer exchange were preserved (Supplementary Figure 5). The assignment by analogy could be performed and both protein samples could be challenged with an addition of aqueous ADH-6 leading to protein-to-ligand ratios of 1:11 (for WT p53 DBD) and 1:8 to 1:15 (for R248W p53 DBD). This time several specific peak shifts were observed with both proteins as illustrated in the panels of Figure 2 in the main text.

The experimental chemical shift perturbation (CSP) values that were determined after addition of ADH-6 affect, to different extents, approximately the same group of residues in the two studied variants of p53 DBD (Supplementary Figure 6). However, the interaction with ADH-6 involved also the partially unfolded species that were present in the samples of both variants (Figure 2). Due to this additional involvement and the signal-to-noise ratios, no quantitative estimate was reliably feasible to assess a binding constant of ADH-6 to the WT and mutant R248W DBDs. Finally, reaching a ligand-to-protein ratio of ~30 resulted in unfolding of both WT and mutant proteins (Supplementary Figure 7).

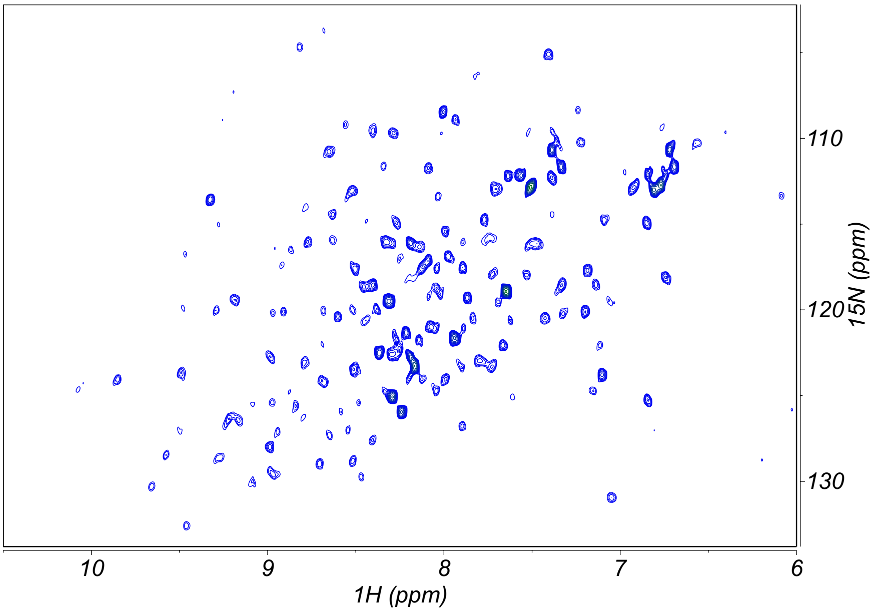


**Supplementary Figure 3.** $^{15}N$-$^{1}H$ HSQC spectrum of 53 µM WT p53 DBD in 50 mM Tris, 200 mM KCl, 5 mM DTT (pH 7.5), recorded at 14 T (600 MHz $^{1}H$ frequency) and 293.2 K.

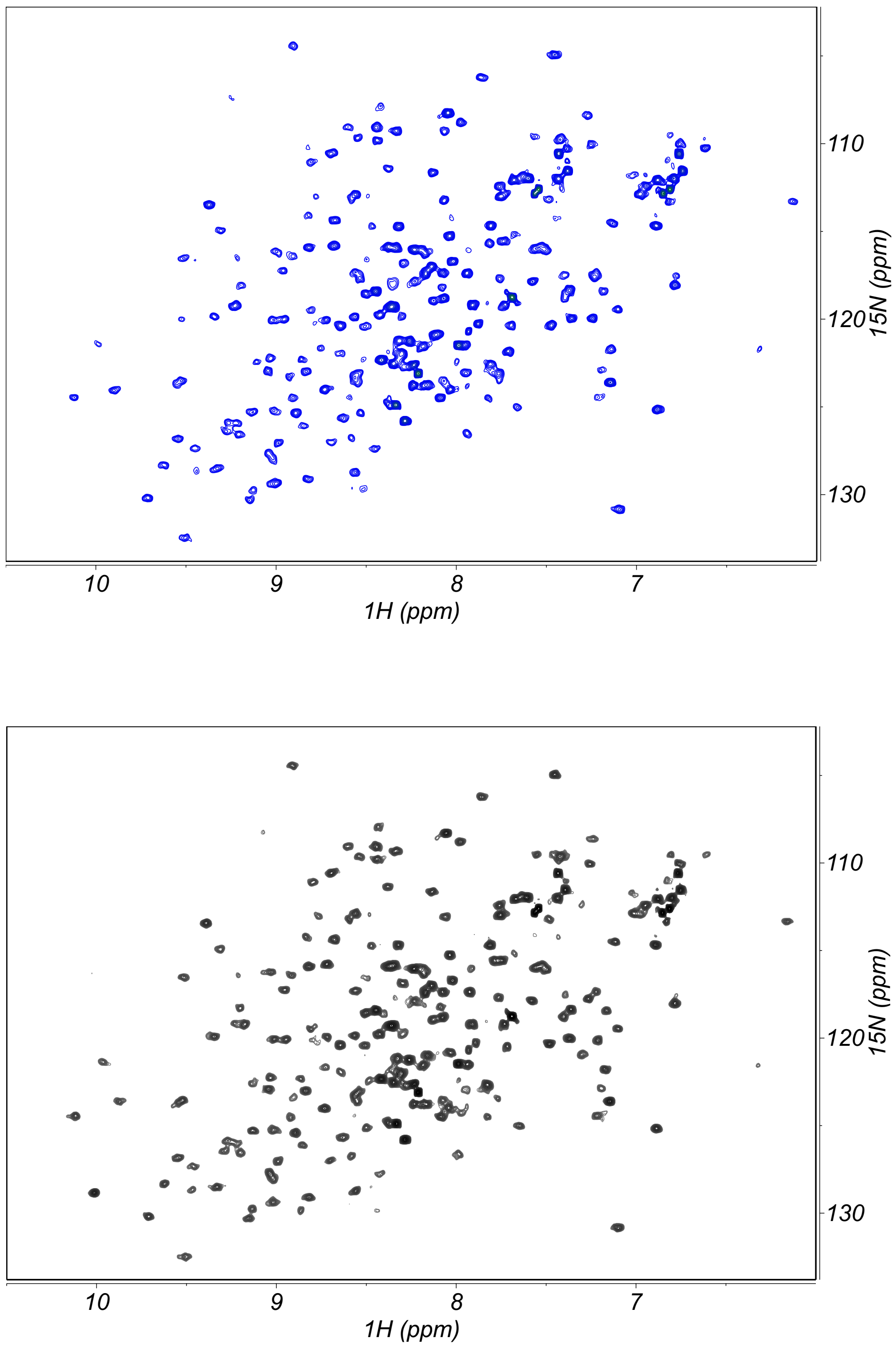


**Supplementary Figure 4.** $^{15}$N-$^{1}$H HSQC spectra of 45 µM WT (*upper panel*, blue contours) and 41 µM mutant R248W (*lower panel*, black contours) p53 DBDs in 47 mM phosphate buffer, 141 mM KCl, 4.7 mM DTT (pH 6.8), recorded at 14 T (600 MHz $^{1}$H frequency) and 293.2 K.

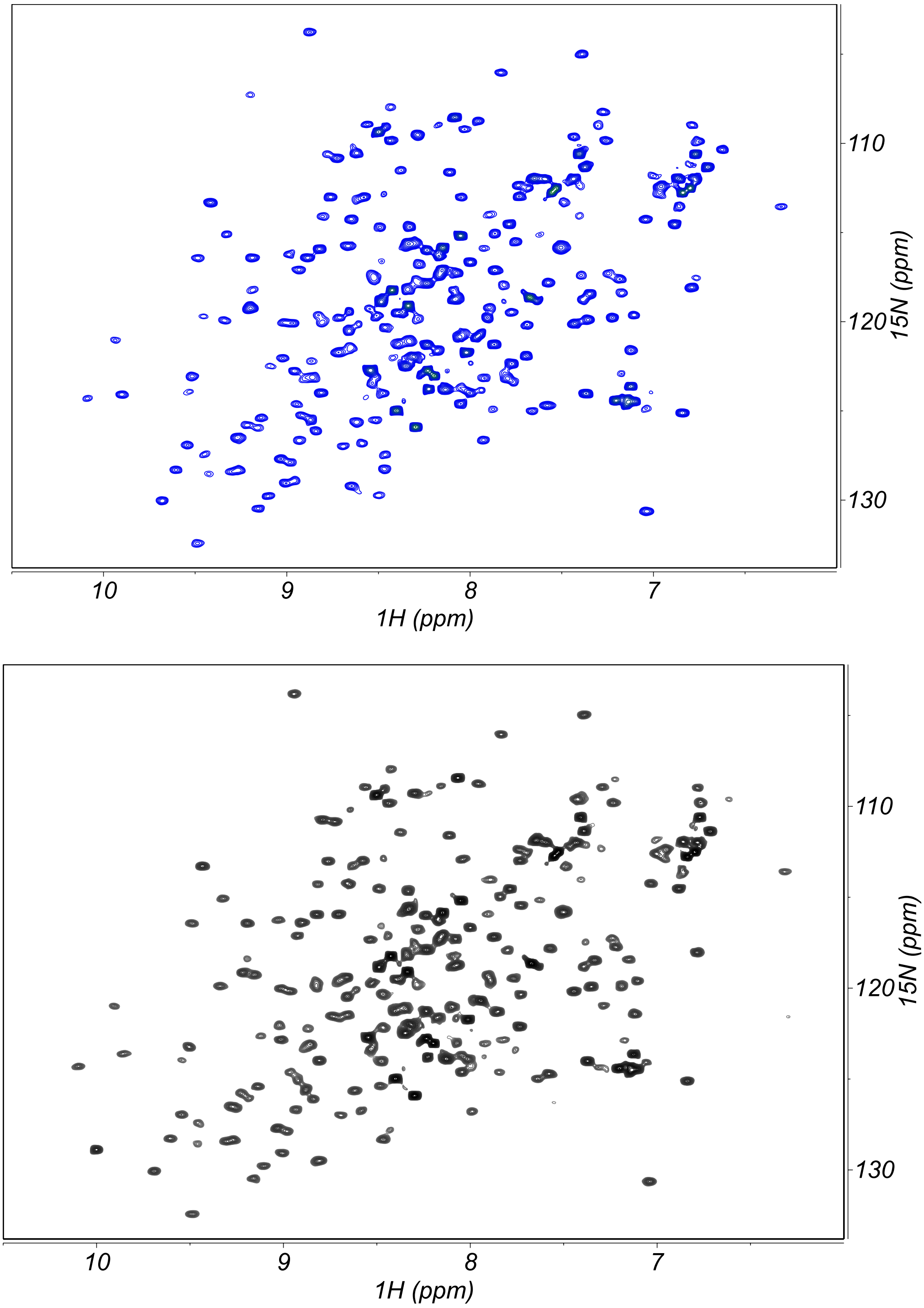


**Supplementary Figure 5.** $^{15}$N-$^{1}$H HSQC spectra of 19 µM WT (*upper panel*, blue contours) and 24 µM R248W (*lower panel*, black contours) p53 DBDs in $H_2O/D_2O$ (96/4), 16.7 mM DTT, pH 5.6 (WT DBD) or 5.8 (mutant DBD), recorded at 14 T (600 MHz $^{1}$H frequency) and 293.2 K.

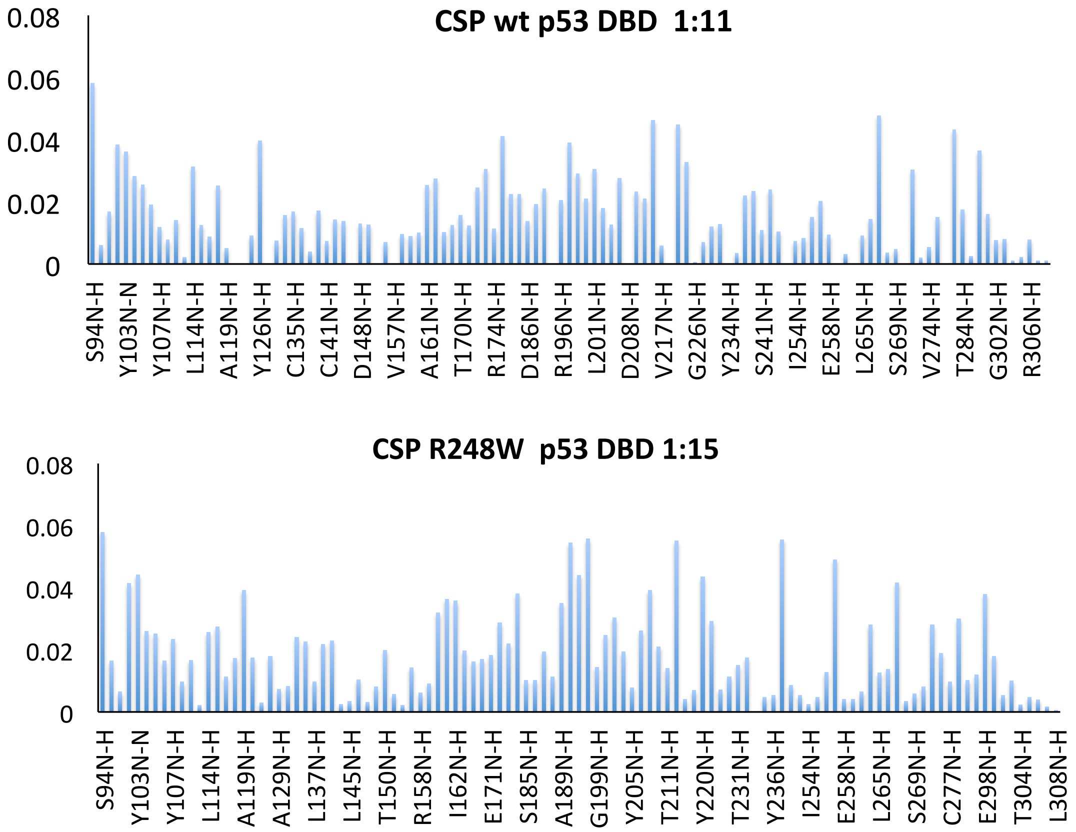


**Supplementary Figure 6.** Chemical Shift Perturbation (CSP) values from $^{15}N$-$^{1}H$ HSQC spectra of WT and R248W mutant p53 DBDs in the presence of ADH-6 at the indicated protein-to-ligand ratios.

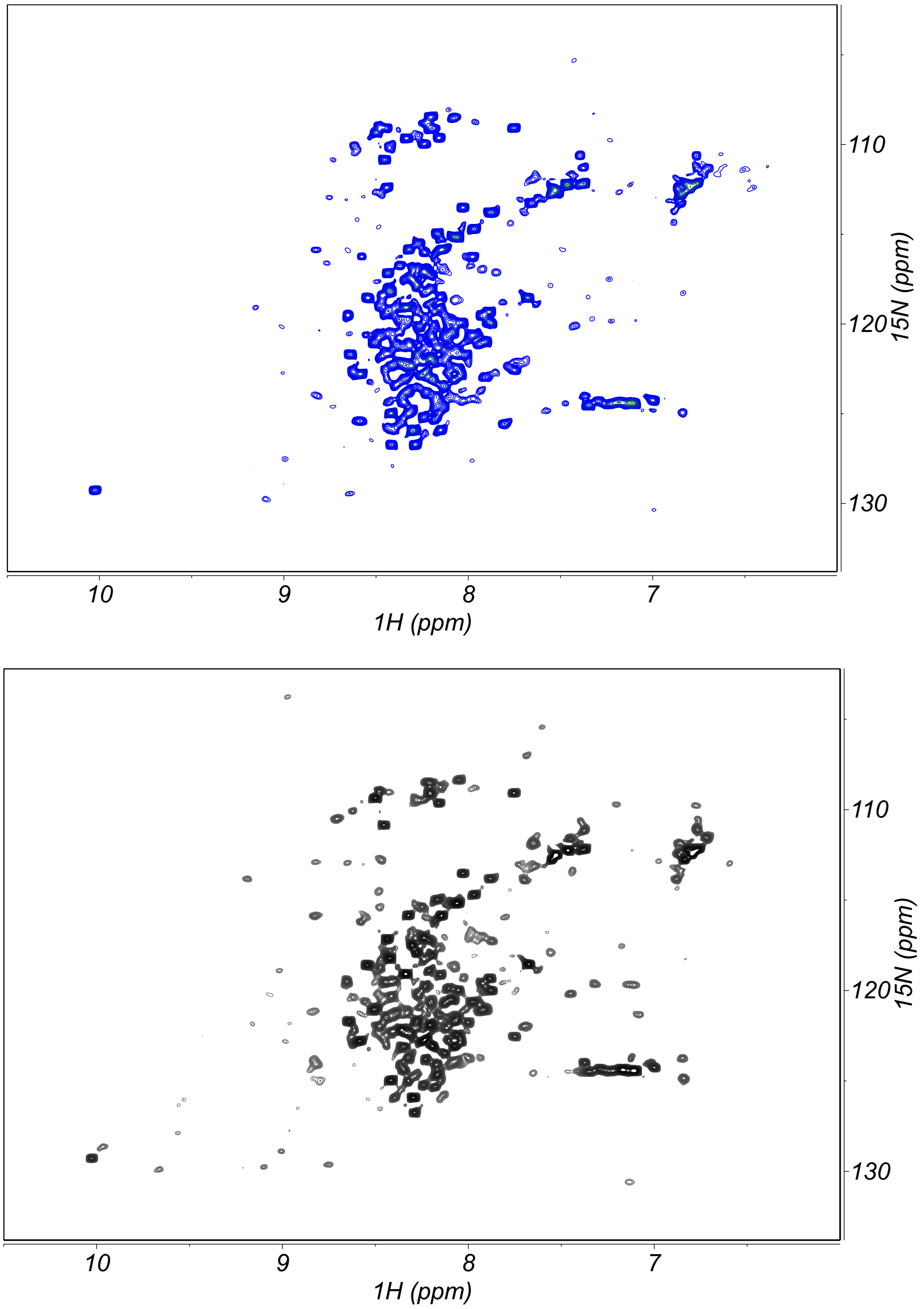


**Supplementary Figure 7.** $^{15}N$-$^{1}H$ HSQC maps of 18.5 μM WT (*upper panel*, blue contours) and 23.5 μM R248W (*lower panel*, black contours) p53 DBDs in $H_2O/D_2O$ (96/4), 16.4 mM DTT, pH 5.6 (WT DBD) or 5.8 (mutant DBD), following the addition of aqueous ADH-6 reaching a protein-to-ligand ratio of 1:30. Both proteins appear extensively unfolded. The spectra were recorded at 14 T (600 MHz $^{1}H$ frequency) and 293.2 K.

### SECTION 3. Supplementary intracellular localization and cell viability/toxicity data

#### 3.1. Supplementary intracellular localization imaging

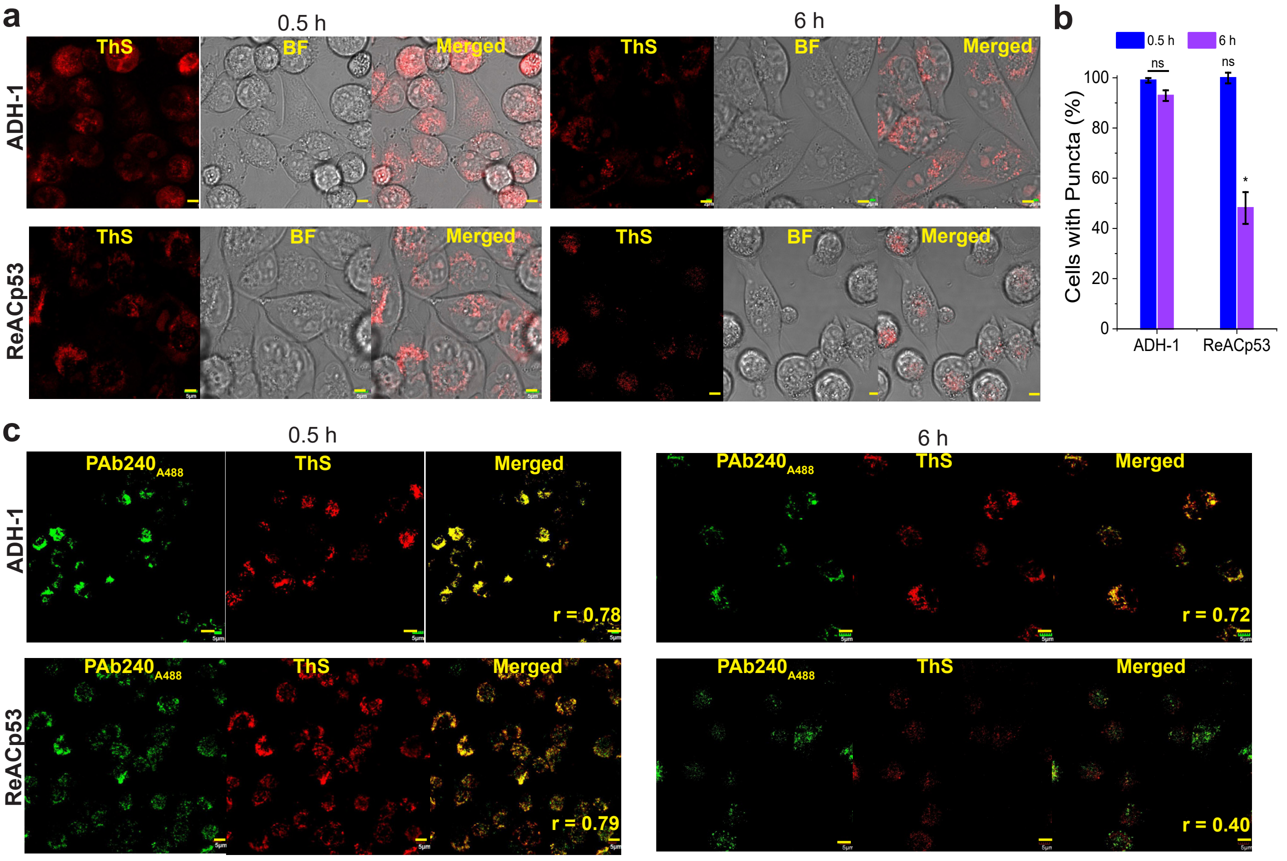


**Supplementary Figure 8. Effects of ADH-1 and ReACp53 on cytosolic mutant p53 aggregates in mutant p53 bearing cancer cells.** (**a**) Confocal fluorescence microscopy images showing thioflavin S (ThS) staining of mutant p53 (R248W) aggregates in MIA PaCa-2 cells treated with vehicle (0.02% DMSO) or 5 µM ADH-1 or ReACp53 for the indicated durations. **(b)** Quantification of ThS-positive MIA PaCa-2 cells after treatment with ADH-1 or ReACp53. The number of positively stained cells in 3–5 different fields of view are expressed as % of the total number of cells ($n$ = 3). *$P$ < 0.05 or non-significant (ns, $P$ > 0.05) for comparisons with controls and between the two incubation times within the same treatment group. (**c**) Confocal fluorescence microscopy images of ThS and PAb 240 antibody staining of R248W aggregates in MIA PaCa-2, treated with 5 µM ADH-1, ReACp53 or ADH-6 for the 0.5 or 6 h. Imaging experiments were performed in triplicate and representative images are shown. Colocalization was quantified using Pearson's correlation coefficient, r[3]. Scale bar = 5 µM.

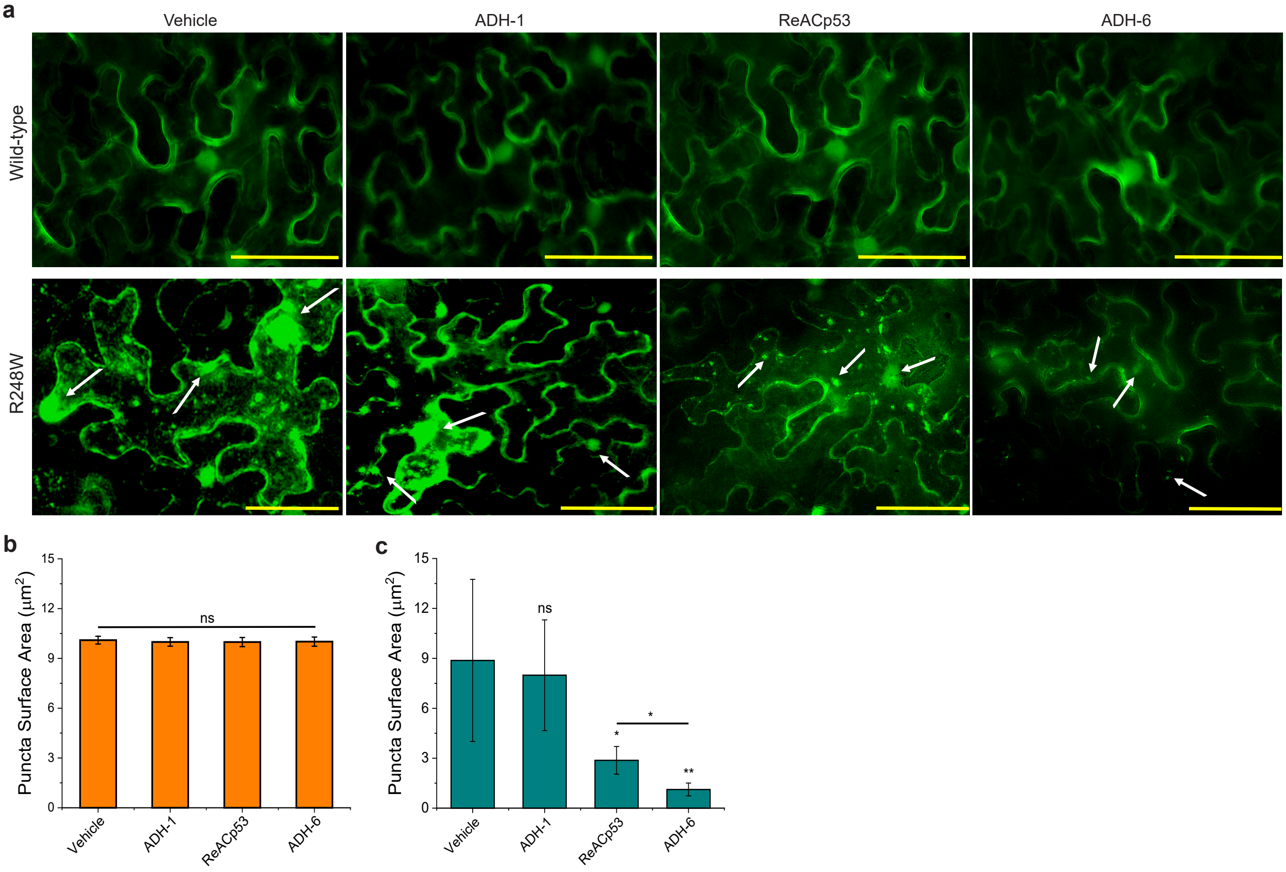


**Supplementary Figure 9. ADH-6 reduces puncta in plant cells expressing mutant, but not WT, p53 DBD.** YFP:p53DBD$^{wt}$ and YFP:p53DBD$^{R248W}$ (YFP-tagged WT and mutant R248W p53 DBDs, respectively) were expressed from the 35S-promoter by agroinfiltrations in *Nicotiana benthamiana* leaves. YFP:p53DBD$^{WT}$ and YFP:p53DBD$^{R248W}$ were infiltrated at an OD of 1.0. 5 µM ADH-1, ReACp53 and ADH-6 were introduced 48 hpi into the bacterial infiltrated region of the leaf, and the effects of the treatments on YFP:p53DBD$^{WT}$ and YFP:p53DBD$^{R248W}$ puncta were observed 24 h later using confocal fluorescence microscopy. (**a**) Images of *N. benthamiana* leaves expressing YFP:p53DBD$^{WT}$ (*upper panels*) or YFP:p53DBD$^{R248W}$ (*lower panels*) treated with ADH-1, ReACp53 or ADH-6. (**b**,**c**) Effects of the treatments on YFP:p53DBD$^{WT}$ (**b**) and YFP:p53DBD$^{R248W}$ (**c**) puncta in *N. benthamiana* leaves. Puncta sizes were quantified in 3–5 different fields of view ($n$ = 3). *$P$ < 0.05, **$P$ < 0.01 or non-significant (ns, $P$ > 0.05) for comparison with vehicle-treated controls and amongst treatment groups.

#### 3.2. Supplementary cell viability/toxicity data

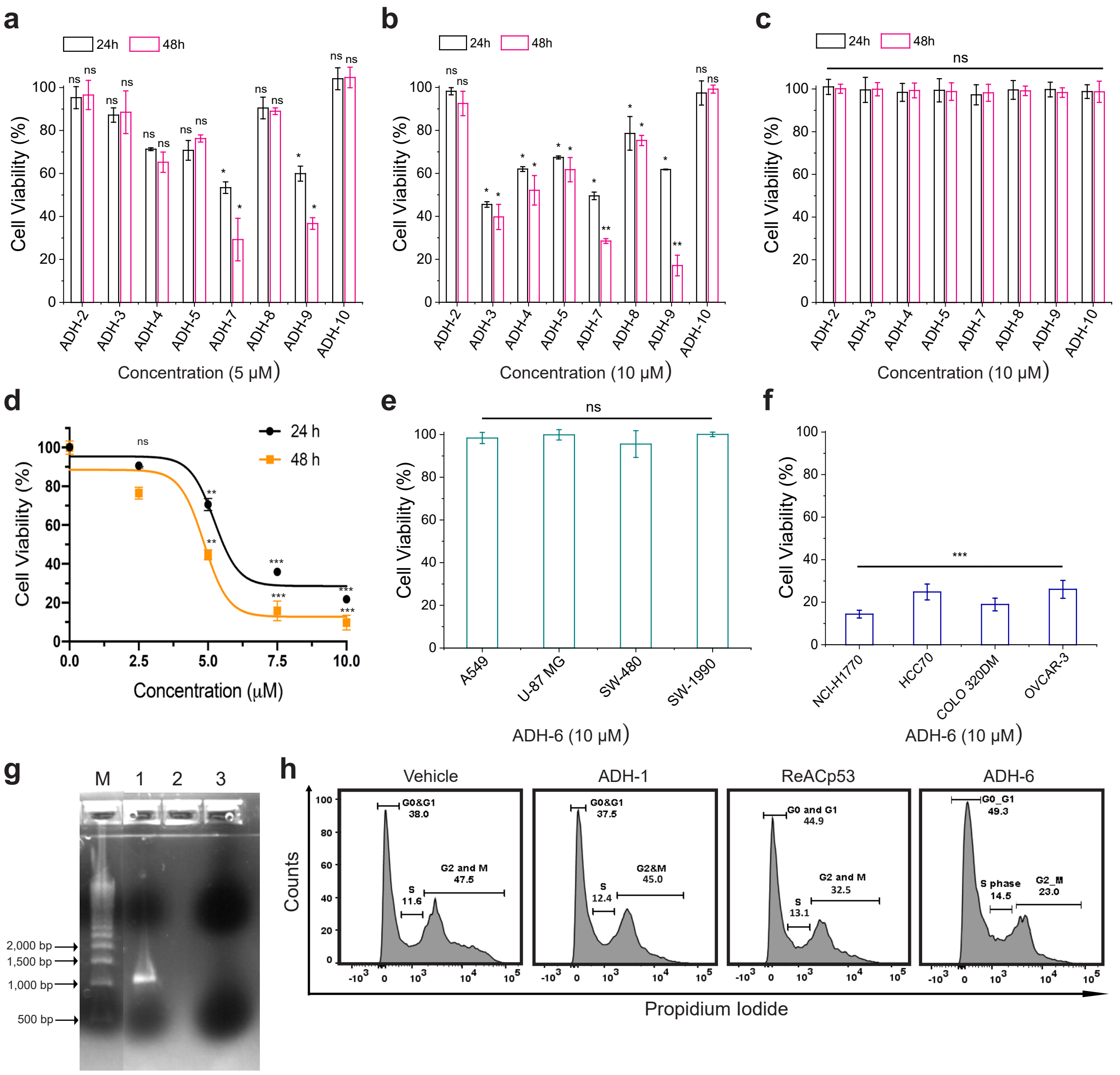


**Supplementary Figure 10. Effects of the oligopyridylamides on cancer cells harboring WT and mutant (R248W) p53.** (**a**–**c**) Screen to identify oligopyridylamides that are toxic to cancer cells bearing mutant, but not WT, p53. MIA PaCa-2 (mutant R248W p53) (**a**,**b**) and MCF-7 (WT p53) (**c**) cells were treated with the indicated concentrations of ADH compounds for 24 or 48 h. (**d**) Effects of control peptide ReACp53 on cancer cells bearing mutant p53. MIA PaCa-2 cells were treated with the indicated concentrations of ReAcp53 for 24 or 48 h. (**e**,**f**) Probing the effects of ADH-6 on viability of different cancer cells bearing WT (**e**) or mutant (**f**) p53. Lung (A549), brain (U-87 MG), colon (SW-480) and pancreatic (SW-1990) cancer cells bearing WT p53, or aggregation-prone mutant p53-bearing lung (NCI-H1770; mutant R248W p53), breast (HCC70; R248Q), colon (COLO 320DM; R248W) and ovarian (OVCAR-3; R248Q) cancer cells, were treated with 10 μM ADH-6 for 48 h. Cell viability (**a**–**f**) was assessed using the MTS assay, with the % viability determined form the ratio of the absorbance of the treated cells to the control cells (treated with vehicle alone). (**g**) Verification of successful transfection of SKOV-3 cells with mutant R248W p53. The vector was purified from transfected SKOV-3 cells, and the amplified *amp*$^r$ PCR product was electrophoresed on a 1.2% E-gel containing a SYBR Safe stain. Lane M: 500 bp Bio-Rad ladder; lane 1: the *amp*$^r$ gene from transfected SKOV-3 cells; and lane 3: negative control. (**h**) Effects of ADH-6 on cell cycle distribution of mutant p53 bearing cancer cells. MIA PaCa-2 cells were treated with vehicle or 5 μM ADH-1, ReACp53 or ADH-6 for 6 h. Cell cycle distribution of the cells was then evaluated using a cell cycle assay kit (Abcam), with measurements done on a BD FACSAria III cell sorter ($n$ = 4). Shown are representative flow cytometry histograms for the different treatment groups. *$P$ < 0.05, **$P$ < 0.01, ***$P$ < 0.001 or non-significant (ns, $P$ > 0.05) compared with controls.

### SECTION 4. Supplementary transcriptome and proteome analysis

#### 4.1. Supplementary transcriptome analysis

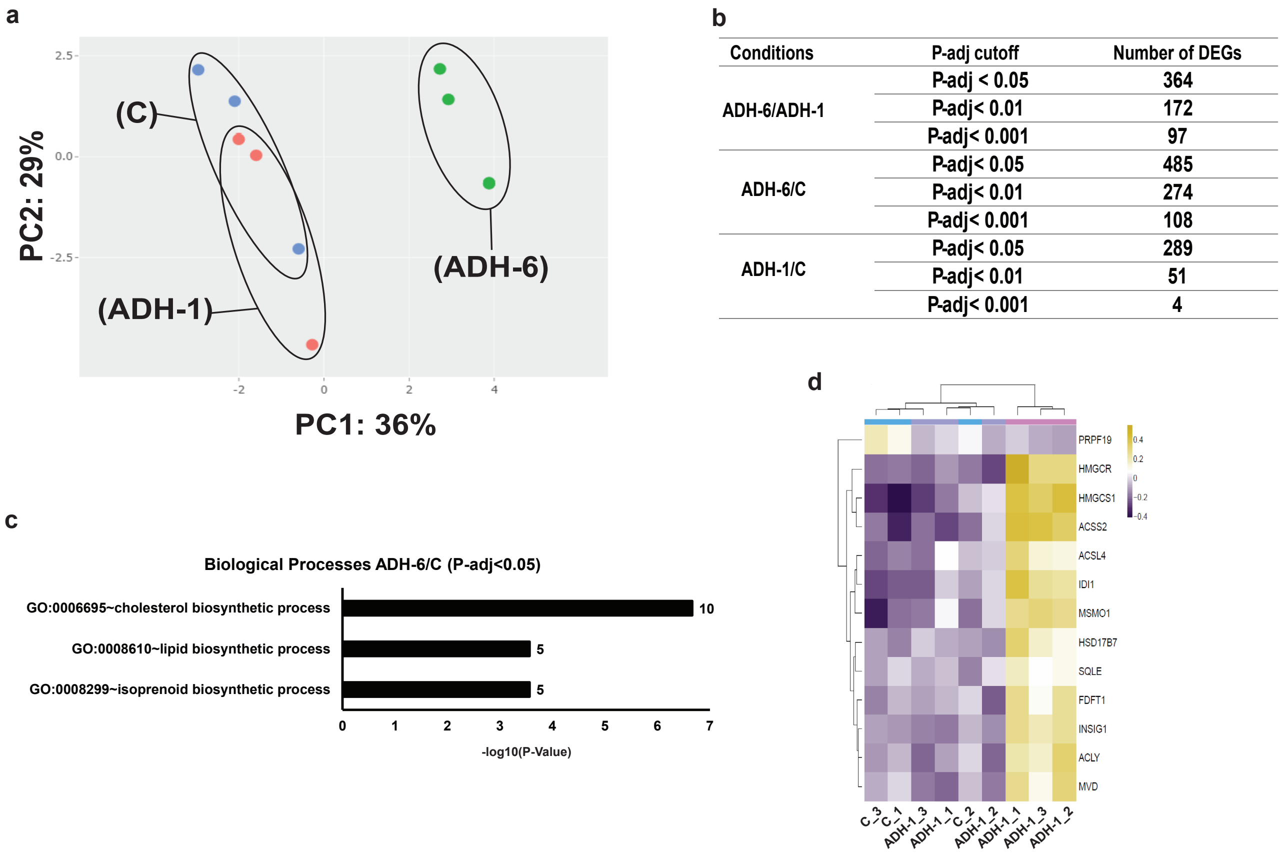


b

| Conditions | P-adj cutoff | Number of DEGs |
|---|---|---|
| ADH-6/ADH-1 | P-adj < 0.05 | 364 |
| | P-adj< 0.01 | 172 |
| | P-adj< 0.001 | 97 |
| ADH-6/C | P-adj< 0.05 | 485 |
| | P-adj< 0.01 | 274 |
| | P-adj< 0.001 | 108 |
| ADH-1/C | P-adj< 0.05 | 289 |
| | P-adj< 0.01 | 51 |
| | P-adj< 0.001 | 4 |

**Supplementary Figure 11. Determination of the best condition for differential gene expression analysis.** (**a**) Principal component analysis (PCA) illustrating total transformed variances including PC1 (36%) and PC2 (29%) for the vehicle-treated controls (C) and ADH-1 and ADH-6 treated cells. (**b**) The number of differentially expressed genes (DEGs) for comparative pairwise analysis for ADH-6/ADH-1, ADH-6/C and ADH-1/C based on statistical significance cut-offs of *P*-values less than 0.05, 0.01 and 0.001. (**c**) Gene ontology (GO) term analysis (biological processes) based on ADH-6/C (*P*-value < 0.05) displaying enrichments involved in cholesterol, lipid and isoprenoid biosynthetic processes. (**d**) A metric heatmap (scaled to log2cpm_voom) displaying expression patterns of all 15 DEGs from the previous GO term analysis.

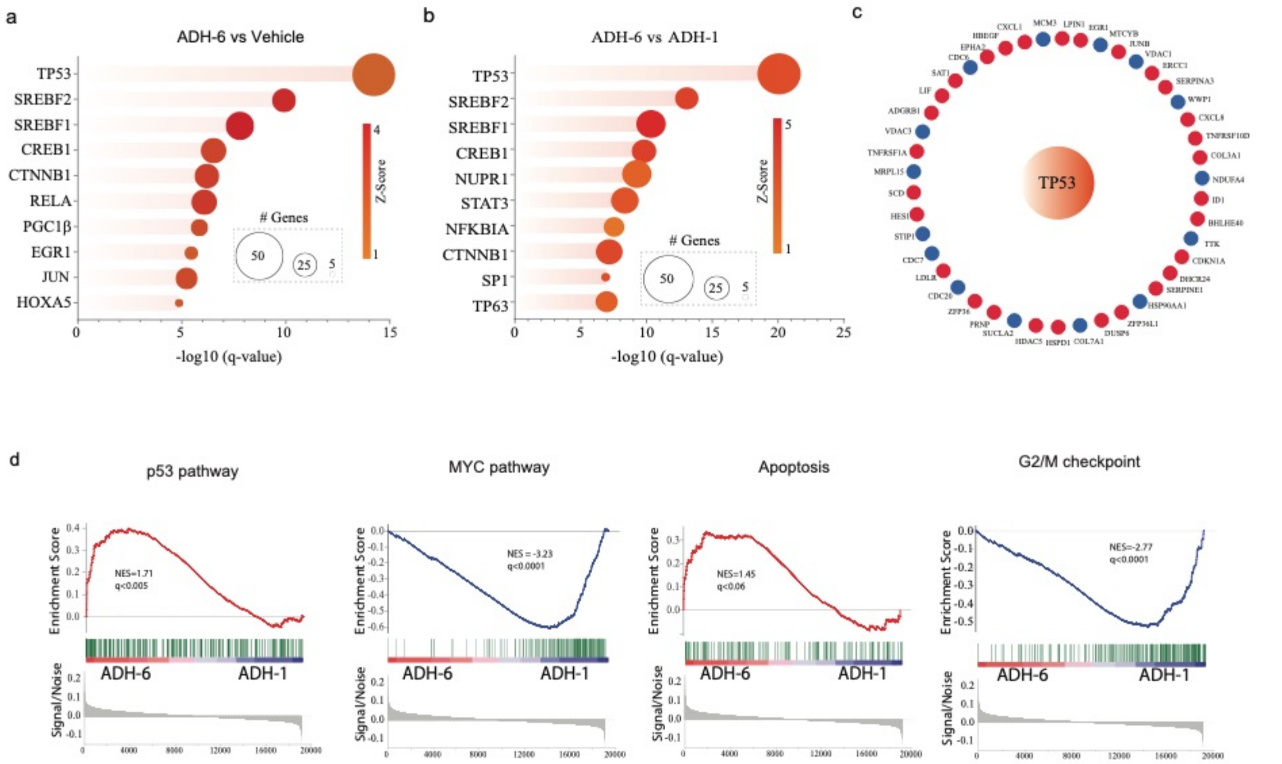


**Supplementary Figure 12. Identification of transcriptional regulators of dysregulated genes in oligopyridylamide-treated MIA PaCa-2 cells.** (**a**,**b**) Ingenuity pathway analysis (IPA) was performed on differentially expressed genes (DEGs) to identify transcriptional regulators (TRs) responsible for gene dysregulation in MIA PaCa-2 cells in ADH-6 vs control (**a**) and ADH-6 vs ADH-1 (**b**) samples. The top 10 TRs in each comparison (ranked based on q-values with a z-score cutoff of 1.4) are shown on the y-axis. The q-value (x-axis) for each TR represents the number of DEGs in the known pathway that overlapped with genes from our dataset. Circles sizes represent the subset of the DEGs with an expression pattern consistent with pathway activation. (**c**) Network plot for the genes that predict activation of the *TP53* pathway in the ADH-6 vs ADH-1 comparison. Red and blue nodes represent upregulated and downregulated genes, respectively. (**d**–**f**) Gene set enrichment analysis (GSEA) using gene expression data from ADH-6 and ADH-1 treated MIA PaCa-2 cells. The enrichment plots for *TP53* pathway, MYC pathway, apoptosis, and G2/M checkpoint are shown along with their normalized enrichment scores (NES) and q-values. Genes in the enrichment plots are marked with green vertical bars, and the gene enrichment scores are shown on the y-axis and ordered according to their signal/noise ratio. Upregulated pathways in the ADH-6 treatment group are represented by red lines (p53 and apoptosis) while downregulated pathways are represented by blue lines (MYC and G2/M).

#### 4.2. Supplementary proteome analysis

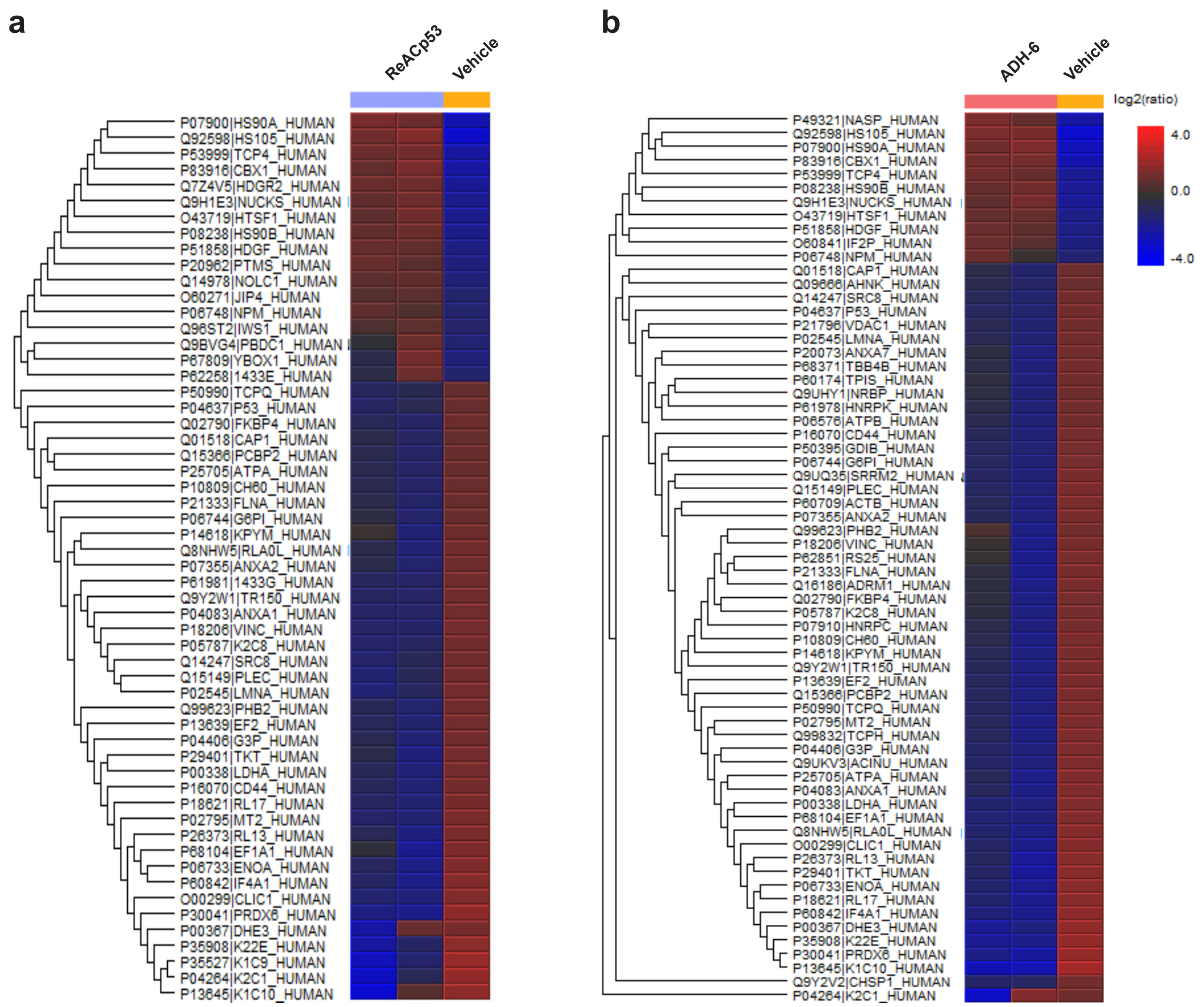


**Supplementary Figure 13. Phosphoproteome analysis.** Heat map representation of the identified proteins in the vehicle (control), ADH-1, ADH-6 and ReACp53 treated groups and their corresponding protein abundances. The digests were phosphopeptide enriched using titanium dioxide ($TiO_2$), tandem mass tag (TMT) labeled, combined in equal amounts (1:1), and analysed by liquid chromatography tandem mass spectrometry (LC-MS/MS). Reporter ion intensities in the $MS^2$ spectra for the TMT labeled phosphopeptides were used to quantify protein abundance. As shown in the color scale bar, red indicates proteins that are upregulated, while dark blue colour signifies proteins that are downregulated.

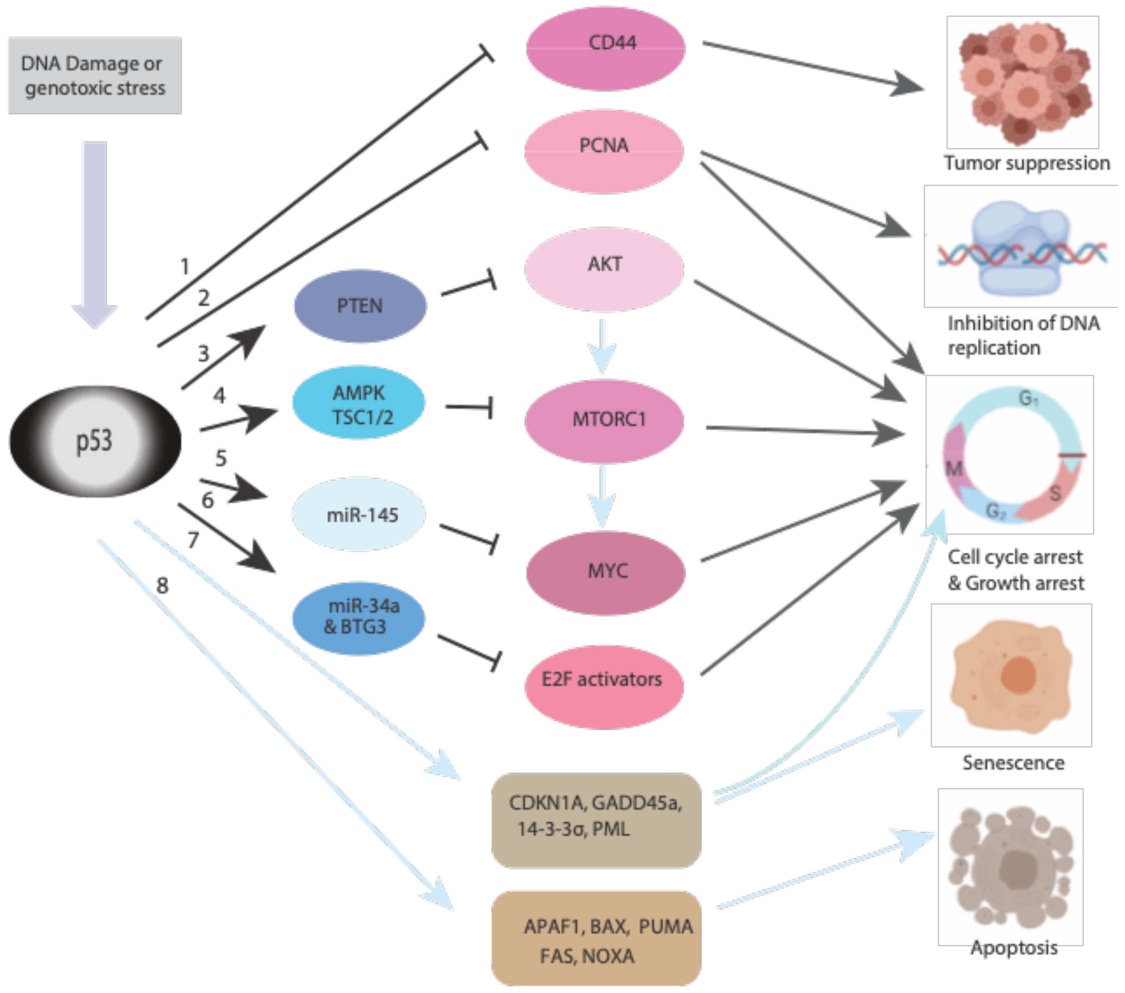


**Supplementary Figure 14. A simplified model of p53 mediated regulation of DNA replication/ repair and cell cycle progression/proliferation**. (**1**) p53 is reported to suppress the expression of CD44 (cluster of differentiation 44), a pro-survival protein, by binding to its promoter[4]. (**2**) Elevated expression of p53 downregulates PCNA (proliferating cell nuclear antigen) leading to the inhibition of DNA replication and cell cycle progression[5,6]. (**3**,**4**) DNA damage induced p53 expression is reported to suppress mTORC1 (mechanistic target of rapamycin complex 1) through AMPK (5′ adenosine monophosphate-activated kinase) and REDD1 (regulated in development and DNA damage 1) responses via the TSC1/2 (tuberous sclerosis proteins 1 and 2) complex and through the transactivation of PTEN (phosphatase and tensin homolog)[7–9]. (**5**) p53 induces the expression of the tumor suppressor microRNA, miR-145, which represses expression of c-Myc (avian myelocytomatosis virus oncogene cellular homolog) post-transcriptionally[10,11]. (**6**) Stress induced p53 represses the E2F (E2 promoter binding factor, an activator) pathway by inducing miR-34 and BTG3 (B-cell translocation gene 3)[12–14]. The downregulation of proliferative pathways (**2**–**6**) results in cell cycle arrest (black arrows). (**7**,**8**) Under stress, p53 also induces the transactivation of genes that induce senescence and cause cell death[15].

#### Supplementary Table 1. Biological roles of downregulated/upregulated phosphoproteins in DNA repair/replication and cell cycle progression/proliferation

| Supplementary Table 1a. Downregulated phosphoproteins | | | | | | | |
|---|---|---|---|---|---|---|---|
| **Protein** | **Access. Nº** | **Phosphory-lation site(s)** | **Implicated pathway(s)** | **Phosphorylating enzymes** | **Cellular function(s)** | **Refs** | **Cell line(s) used for studies** |
| Heterogeneous nuclear ribonucleo-protein D (HNRNPD) | Q14103 | Ser83 | MYC, E2F & G2M checkpoint | ? | Phosphorylated HNRNPD interacts and co-localizes in the cytoplasm with 14-3-3ζ. Cytoplasmic AUDF1 decreases the stability of cyclin-dependent kinase inhibitors which promotes cell proliferation. (**Cell proliferation**) | 16,17 | Oral squamous cell carcinoma (OSCC) cells & human thyroid carcinoma cells |
| Minichromosome maintenance complex component 2 (MCM2) | P49736 | Ser27, Ser41 & Ser139 | MYC, E2F, G2M checkpoint & MTORC1 signaling | Cdc7/Dbf4 | During cell cycle, phosphorylated MCM2 is proposed to be important for regulation of ATPase activity of MCM2 and initiation of DNA replication. (**DNA repair/replication**) | 18,19 | HeLa cells |
| Lactate dehydrogenase A (LDH-A) | P00338 | Y10 & Y83 | MYC, oxidative phosphoryl ation, glycolysis & MTORC1 | Receptor tyrosine kinase FGFR1 | Phosphorylation at Y10 regulates the activity of LDH-A by increasing the formation of active, tetrameric LDH-A, while phosphorylation at Y83 mediates its binding to its substrate to promote tumor growth. (**Cell proliferation**) | 20 | Various human cancer cell lines, including H1299 |
| Proliferating cell nuclear antigen (PCNA) | P12004 | Y211 | MYC, E2F & P53 | EGF receptor | The chromatin-bound form of PCNA is phosphorylated at Y211 during the S phase of cell cycle to promote DNA replication. (**DNA repair/replication**) | 21,22 | Breast cancer MDA-MB-231cells & epidermoid carcinoma A431 cells |
| Voltage dependent anion channel 1 (VDAC1) | P21796 | S193 | MYC & oxidative phosphory-lation | Nek1/PKC | Phosphorylation of VDAC1 is reported to prevent mitochondria-mediated apoptosis. (**Cell proliferation**) | 23,24 | Human kidney 2 (HK2) cells |
| Prohibitin 2 (PHB2) | Q99623 | Ser91 & Ser176 | MYC & oxidative phosphory-lation | Akt | Phosphorylated PHB2 in leukemia cells is proposed to regulate coordinated nuclear and mitochondrial responses, which increase cell survival. (**Cell proliferation**) | 25,26 | NB4 human leukemia cells |
| Poly(rC) binding protein 1 (PCBP1) | Q15365 | Ser43 | MYC | Akt2 | TGF-β mediates PCBP1 phosphorylation, which promotes epithelial to mesenchymal transition (EMT) in cells that, in turn, contributes to cancer progression. | 27–29 | Non-small cell lung cancer (NSCLC) A549 cells & gallbladder carcinoma GBC-SD cells |
| | | Thr60 & Thr127 | | p21-activated Kinase 1 (Pak1) | Phosphorylated PCBP1 causes the transactivation of eIF4E, leading to initiation of translation to promote cell growth. (**Cell proliferation**) | 30,31 | HeLa cells |
| Protein kinase, DNA-activated, catalytic subunit (PRKDC) | P78527 | S2056, S2609, T2647 & T3950 | E2F | Autophosphory-lation | During mitosis phosphorylated PRKDC is required for chromosome segregation and cell cycle progression. (**Cell proliferation**) | 32,33 | HeLa & HCT116 cells |
| | | S3205 | | Polo-like kinase 1 (PLK1) | Phosphorylation of PRKDC during mitosis is required for proper cytokinesis. (**Cell proliferation**) | 33,34 | HeLa cells |

| | | | | | | | |
|---|---|---|---|---|---|---|---|
| | | Thr-2609, Ser2612, Thr2638 & Thr2647 | | Autophosphory-lation | Phosphorylation of PRKDC leads to conformational changes within the protein that results in efficient DNA end processing and double strand break repair. (**DNA repair/replication**) | 35,36 | Human lymphoblastoid cells |
| Tripartite motif containing 28 (TRIM28) | Q13263 | Ser824 | MYC | ATM & DNA-PK | During DNA damage, phosphorylated TRIM28 results in de-repression of p21 and Gadd45, which causes cell cycle arrest to allow DNA repair. (**DNA repair/replication**) In cancer cells, phosphorylated TRIM28 promotes tumor growth by increasing DNA damage repair. (**Cell proliferation**) | 37–40 | HEK293 cells & A375 melanoma cells |
| | | Ser473 | | PKCδ | During S phase, phosphorylation of TRIM28 removes its repression on Cyclin A2, which leads to cell cycle progression. (**Cell proliferation**) | 41 | HEK293 cells |
| | | Tyr449, Tyr458 & Tyr517 | | Src family of kinases | Phosphorylation inhibits TRIM28's interaction with HP1 and heterochromatin, resulting in a loss of their transcriptional repression activity. It is speculated that TRIM28 enhances oncogenic transformation of cells. (**Cell proliferation**) | 42,43 | COS-1 & HeLa S3 cells |
| Heterogeneous nuclear ribonucleo-proteins C (HNRNPC) | P07910 | Ser240, Ser225 & Ser228 | MYC | CK1α | At physiological levels of $H_2O_2$, phosphorylated HNRNPC is predicted to mediated cell proliferation and survival. (**Cell proliferation**) | 44 | Human umbilical vein endothelial cells (HUVECs) |
| RACK1 | P63244 | Tyr 228 &/or Tyr246 | MYC & p53 | Src | SRC mediated phosphorylation and interaction with RACK1 is predicted to enhance cell survival, proliferation and migration. (**Cell proliferation**) | 45,46 | NIH3T3 cells |
| Fas cell surface death receptor (FAS) | P49327 | Y232 & Y291 | Apoptosis, p53, IL6/JAK/ STAT3 & EMT | Src & Yes | Phosphorylation of the Fas death domain results in suppression of apoptosis. (**Cell proliferation**) | 47,48 | Breast, ovarian & colorectal cancer cell lines |
| Lamin A/C (LMNA) | P02545 | Thr19, Ser22 & Ser392 | Apoptosis | Cdk group | Phosphorylation of Lamin A/C during late mitosis results in its depolymerization of nuclear lamina, which promotes cell cycle progression. (**Cell proliferation**) | 49,50 | Hela Cells |
| ATP citrate lyase (ACLY) | p53396 | Ser454 | MTORC1 | Akt/SREBP | Phosphorylation of ACLY is predicted to regulate cell growth and differentiation by mediating de novo lipogenesis (DNL). (**Cell proliferation**) | 51,52 | Adipocytes & myrAkt-ER cells |
| CD44 | p16070 | Ser325 | Apoptosis, IL6/JAK/ STAT3 & EMT | $Ca^{2+}$/calmodulin-dependent protein kinase II (CaMKII) | Phosphorylated CD44 recruits ezrin protein, which links it to the actin cytoskeleton and is predicted to promote cell proliferation and invasiveness | 53–55 | Flow2000 fibroblasts |

| | | | | | | | |
|---|---|---|---|---|---|---|---|
| | | | | | (**Cell proliferation**) | | |
| Annexin A1 (ANAX1) | p04083 | Ser5 | Apoptosis | TRPM7/Chak1 | Phosphorylated Annexin is predicted to form the complex TRPM7/Annexin A1/$Mg^{2+}$, which is proposed to enhance cell viability and cell cycle progression. (**Cell proliferation**) | 56,57 | Human vascular & smooth muscle cells |
| RAB1A, member RAS oncogene family (RAB1A) | p62820 | Ser194 | MTORC1 | Cdk1 | During mitosis, to ensure equal distribution of organelles between daughter cells, phosphorylation of RAB1A inhibits intracellular transport. (**Cell proliferation**) | 58,59 | HeLa cells |

**Supplementary Table 1b. Upregulated phosphoproteins**

| Protein | Access. Nº | Phosphory-lation site(s) | Implicated pathway(s) | Phosphorylating enzymes | Cellular function(s) | Ref | Cell line(s) used for studies |
|---|---|---|---|---|---|---|---|
| Splicing factor 3b subunit 2 (SF3B2) | Q13435 | Ser289 | DNA damage repair | ATM | During cell cycle, phosphorylated SF3B2 recruits CtIP at double stranded DNA breaks. CtIP also promotes p21 and Gadd45 causing cell cycle arrest. (**DNA repair/replication**) | 60–62 | U2OS cells |
| Vacuolar protein sorting 4 homolog B (VPS4B) | O75351 | Ser102 & Ser108 | Multi-vesicular body (MVB) sorting | CK2α pathway | Phosphorylated VPS4B, along with ESCRTIII subunits, regulates epidermal growth factor degradation through MVB sorting pathway. This is predicted to inhibit cell proliferation. (**Cell proliferation**) | 63,64 | HEK 293T & HeLa cells |

### SECTION 5. Supplementary *in vivo* tumor reduction data

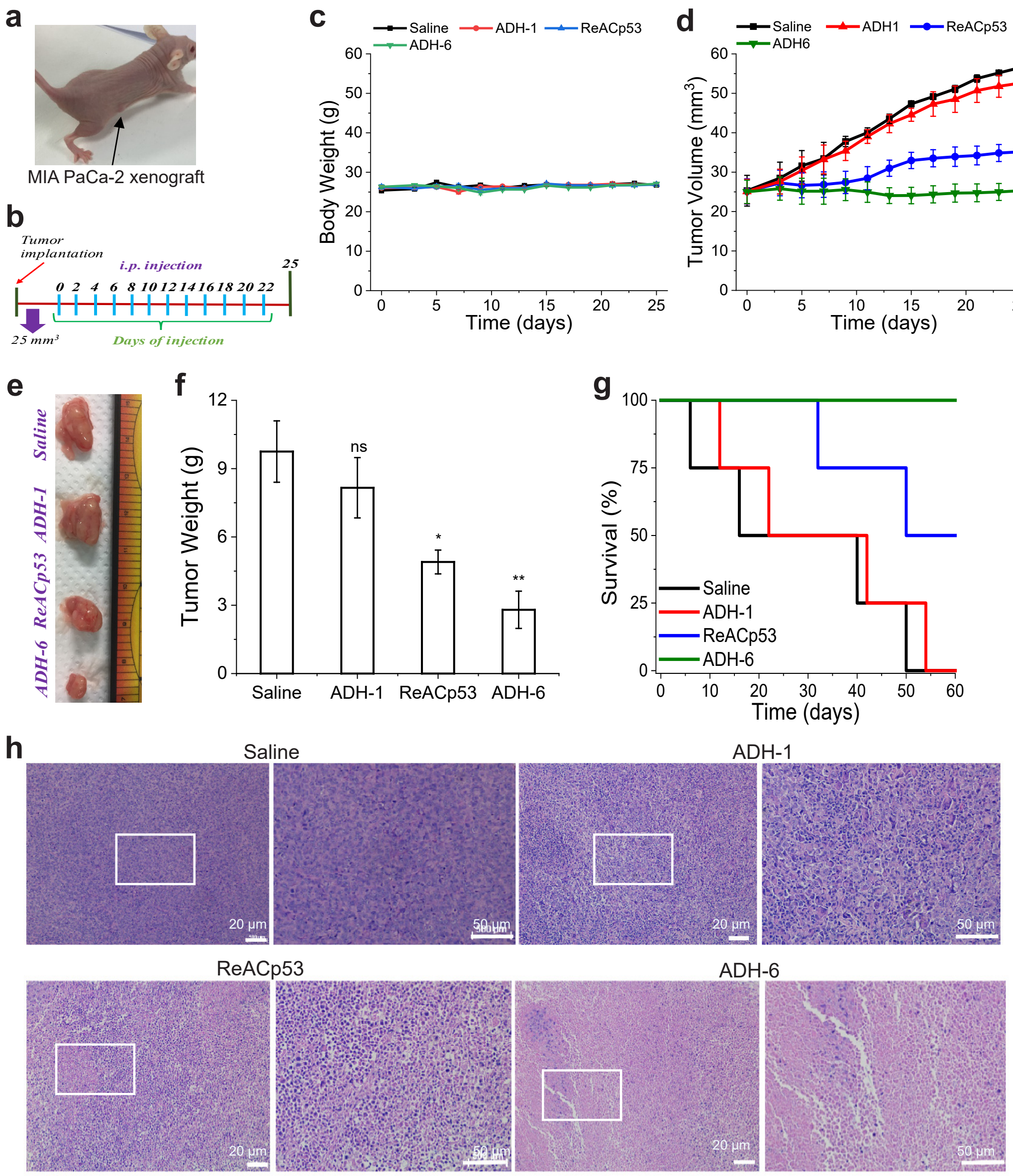


**Supplementary Figure 15. Effect of ADH-6 on xenografts bearing aggregation-prone mutant p53 *in vivo*.** (**a,b**) Design of the tumor reduction studies. A representative mouse bearing both MIA PaCa-2 (mutant R248W p53) xenografts (**a**) and the treatment schedule (**b**). Once the tumor volume reached ~25 mm$^3$, the mice were randomized into the different treatment groups ($n = 8$ per group), which were injected intraperitoneally with saline or 156.4 µM ADH-1, ReACp53 or ADH-6. Injections were done every 2 days for a total of 12 doses, with the first day of treatment defined as day 0. (**c**) Body weight changes of the tumor-bearing mice in the different treatment groups monitored for the duration of the experiment. (**d**) Tumor volume growth curves for the MIA PaCa-2 xenografts in the different treatment groups over 25 days of treatment ($n$ = 8 per group). (**e,f**) Tumor mass analysis for the different treatment groups. After 25 days of treatment, 4 mice per treatment group were sacrificed and the tumor tissues were isolated and imaged (**e**) and subsequently weighed to determine the tumor mass (**f**). (**g**) Survival curves for the saline, ADH-1, ReACp53 and ADH-6 treatment groups over 60 days ($n$ = 4 per group). (**h**) Hematoxylin and eosin (H&E)-stained images of xenograft sections from the different treatment groups following 25 days of treatment. Images on the right are magnified views of the boxed regions in the images on the left. Scale bar = 20 µm (50 µm for the magnified views). *$P < 0.05$, **$P < 0.01$, ***$P < 0.001$ or non-significant (ns, $P > 0.05$) for comparison with controls and amongst the different treatment groups.

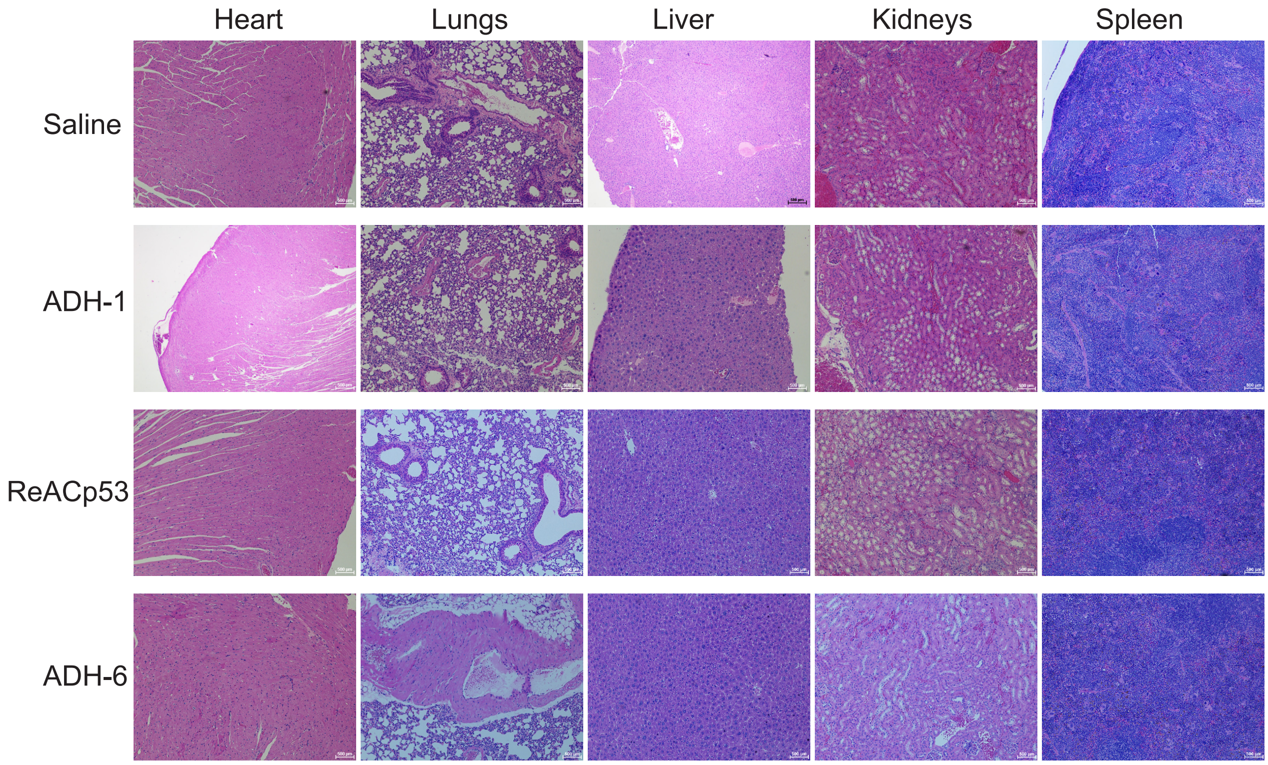


**Supplementary Figure 16. Histological analysis of vital organs following treatment with lower doses of ADH-6.** Hematoxylin and eosin (H&E) staining of heart, lung, liver, kidney and spleen sections from MIA PaCa-2 tumor-bearing mice after 25 days of treatment with saline, ADH-1, ReACp53 or ADH-6 (dosage of 155.6 µM in saline, administered every 2 days, for a total of 12 doses). Scale bar = 50 µm.

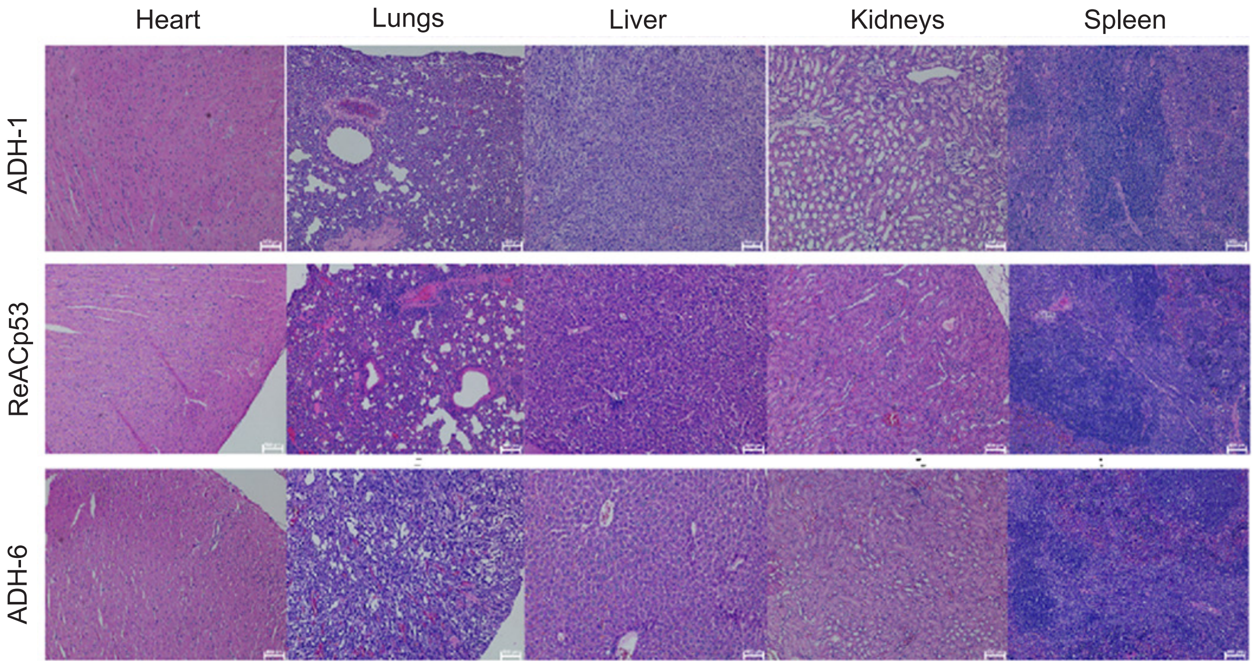


**Supplementary Figure 17. Histological analysis of vital organs following treatment with higher doses of ADH-6.** Hematoxylin and eosin (H&E) staining of heart, lung, liver, kidney and spleen sections from MIA PaCa-2 tumor-bearing mice after 25 days of treatment with ADH-1, ReACp53 or ADH-6 (dosage of 716.4 μM in 0.02% DMSO, administered every 2 days, for a total of 12 doses). Scale bar = 50 μm.

### SECTION 6. Synthesis and characterization of ADH-6

The synthesis and characterization of the monomer pyridyls have been reported previously[65].

#### General method for the reduction of arylamides

To a solution of nitro arylamide (0.1 mmol) in EtOAc (10 mL), Pd/C (10% by wt) was added and the reaction started with constant stirring in an $H_2$ (g) atmosphere at room temperature. The progress of the reaction was monitored using thin layer chromatography (TLC). The disappearance of the starting material confirmed the completion of the reaction. The reaction mixture was filtered and the filtrate was dried on a rotovap to afford the desired product as a yellow solid, which was used in the next step without further characterization.

#### General method for the deprotection of oligopyridylamides

To a solution of the oligopyridylamide (50 µmol) in dichloromethane (5 mL), triethylsilane (250 µL) was added, followed by addition of trifluoroacetic acid (TFA, 500 µL), and the reaction solution was stirred constantly for 4 h. The reaction solution was dried on a rotovap and washed with cold diethyl ether (3 × 5 mL) which resulted in a yellow powder.

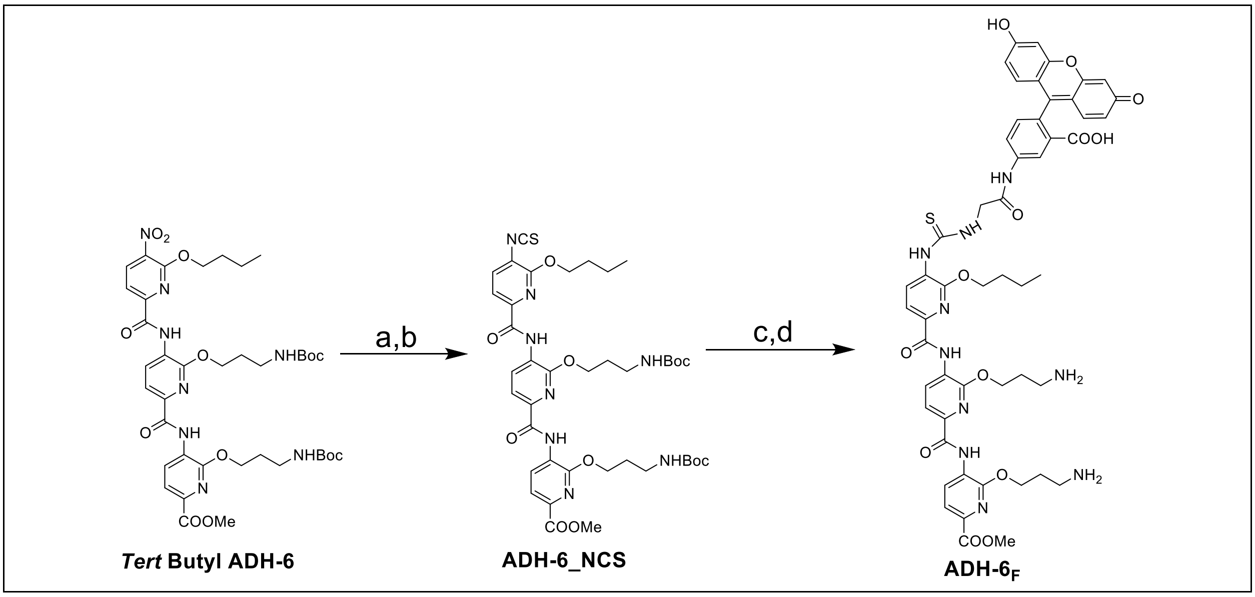


**Scheme S1. Synthetic route for the synthesis of ADH-6$_F$ (ADH-6$_{FITC}$).**

#### ADH-6

The synthesis and characterization of ADH-6 (previously identified as ADH-40) was reported elsewhere[66].

**ADH-6-NH2**

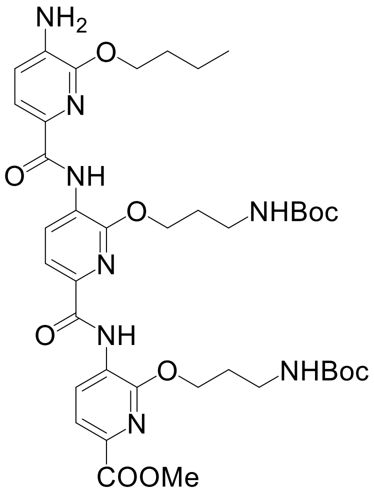


To a solution of ADH-6 (0.1 mmol) in EtOAc (10 mL), Pd/C (12% by wt) was added and the reaction started with constant stirring in an $H_2$ (g) atmosphere at room temperature. The progress of the reaction was monitored using TLC. The disappearance of the starting material confirmed the completion of the reaction (~4 h). The reaction mixture was filtered, and the filtrate was dried on a rotovap to afford the desired product as a yellow solid (yield = 86%), which was used in the next step without further characterization.

**ADH-6-NCS**

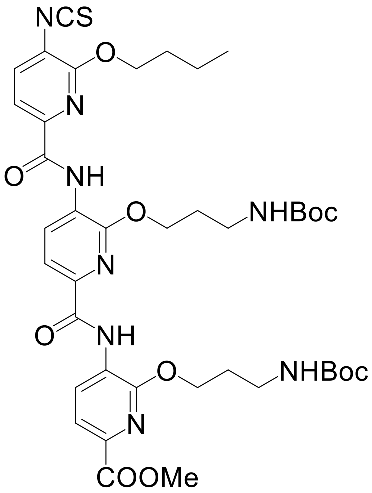


To a solution of **ADH-6-NH2** (40 mg, 0.049 mmol) was dissolved in dicholoromethane (anhydrous, 10 mL), followed by the addition of 1,1'-Thiocarbonyldi-2(1H)-pyridone (22.7 mg, 0.098 mmol, 2 eq.) and the reaction solution was stirred for 6 h at room temperature under inert atmosphere. The progress of the reaction was monitored by TLC. Flash chromatography (0 to 60% Ethyl acetate in hexane) yielded the desired product as a yellow solid (36.6 mg, 87%). $^1$H NMR (600 MHz, Chloroform-*d*) δ $^1$H NMR (600 MHz, Chloroform-*d*) δ 10.32 – 10.28 (s, 1H), 10.25 – 10.22 (s, 1H), 9.04 – 9.00 (d, $J$ = 8.0 Hz, 1H), 8.94 – 8.90 (d, $J$ = 8.0 Hz, 1H), 8.47 – 8.42 (d, $J$ = 7.8 Hz, 1H), 8.04 – 7.99 (d, $J$ = 8.0 Hz, 2H), 7.88 – 7.83 (d, $J$ = 8.0 Hz, 1H), 4.72 – 4.67 (t, $J$ = 6.3 Hz, 2H), 4.66 – 4.62 (t, $J$ = 6.2 Hz, 4H), 4.13 – 3.85 (s, 3H), 3.44 – 3.38 (q, $J$ = 6.2 Hz, 2H), 3.32 – 3.28 (m, 2H), 2.20 – 2.12 (p, $J$ = 6.6 Hz, 2H), 2.12 – 2.03 (h, $J$ = 5.2, 3.8 Hz, 2H), 1.97 – 1.90 (m, 2H), 1.71 – 1.59 (m, 5H), 1.49 – 1.46 (s, 9H), 1.41 – 1.37 (s, 9H). MS-ESI (*m/z*): calculated for $C_{40}H_{52}N_8O_{11}S$ (M): 852.9610, found 852.9689.

***Tert* Butyl ADH-6$_F$**

To a solution of **ADH-6-NCS** (25 mg, 0.029 mmol) in pyridine (5 ml, anhydrous), N, N-diisopropylethylamine (0.005 ml, 0.05 mmol) was added and the solution was stirred for 10 min. To this solution, 5-(aminoacetamido) fluorescein (23.9 mg, 0.056 mmol) was added and the reaction was started in dark with continuous stirring under an inert atmosphere. The reaction solution was stirred overnight in the dark. The product was purified using column chromatography (0–20% methanol in dichloromethane with 1% triethylamine, v/v) as an orange solid (20 mg, 54%). The compound (*tert*-butyl ADH-6$_F$) was characterized via MALDI-TOF and used in the next step without any further characterization. The $^1$H NMR peaks were very broad potentially because of the stacking of the molecule. We used the compound in the next step without further characterization.

#### ADH-6$_F$ (ADH-6$_{FITC}$)

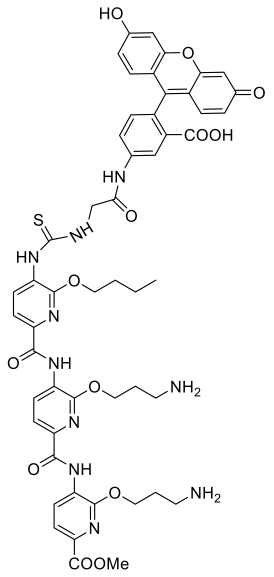


To a solution of **tert-butyl ADH-6$_F$** (17 mg, 0.014 mmol) in dichloromethane (4 mL), triethylsilane (0.1 mL, excess) was added, followed by the addition of TFA (0.4 mL, excess) and the reaction solution was stirred in the dark at room temperature for 4 h. The solution was then dried, and the orange solid was washed with cold diethyl ether (3 × 5 mL), which afforded the desired product (ADH-6$_F$) as an orange solid. The product was further reconstituted in DMSO/water (75:25, v/v) and purified on a semi-preparative reverse-phase HPLC. Solvents A (95% water, 5% ACN and 0.1% TFA) and B (95% ACN, 5% water and 0.1% TFA) were used for the purification. The desired product (ADH-6$_F$) was obtained as an orange solid (11.8 mg, 80%). $^1$H NMR (600 MHz, DMSO-$d_6$) δ 10.78 – 10.70 (s, 1H), 10.29 – 10.27 (s, 1H), 10.26 – 10.23 (m, 1H), 10.17 – 10.11 (s, 2H), 9.71 – 9.60 (s, 1H), 9.03 – 8.99 (d, *J* = 8.0 Hz, 1H), 8.95 – 8.89 (t, *J* = 6.6 Hz, 1H), 8.86 – 8.78 (d, *J* = 7.9 Hz, 1H), 8.41 – 8.33 (s, 1H), 8.00 – 7.95 (dd, *J* = 8.1, 2.9 Hz, 1H), 7.30 – 7.22 (d, *J* = 8.2 Hz, 1H), 6.70 – 6.67 (d, *J* = 2.2 Hz, 2H), 6.63 – 6.58 (d, *J* = 8.5 Hz, 2H), 6.57 – 6.52 (d, *J* = 7.8 Hz, 4H), 4.70 – 4.64 (t, *J* = 5.8 Hz, 2H), 4.63 – 4.58 (t, *J* = 6.3 Hz, 2H), 4.58 – 4.53 (q, *J* = 7.3, 6.8 Hz, 2H), 4.52 – 4.48 (d, *J* = 4.9 Hz, 2H), 3.98 – 3.78 (s, 3H), 3.08 – 3.01 (m, 6H), 2.24 – 2.18 (p, *J* = 7.5, 7.0 Hz, 2H), 2.18 – 2.11 (p, *J* = 6.9 Hz, 2H), 1.97 – 1.85 (h, *J* = 6.9 Hz, 2H), 1.64 – 1.51 (dh, *J* = 14.4, 7.4 Hz, 3H). MS-ESI (*m/z*): calculated for $C_{55}H_{50}N_{10}O_{13}S$ (M+H): 1058.1130, found 1058.1148.

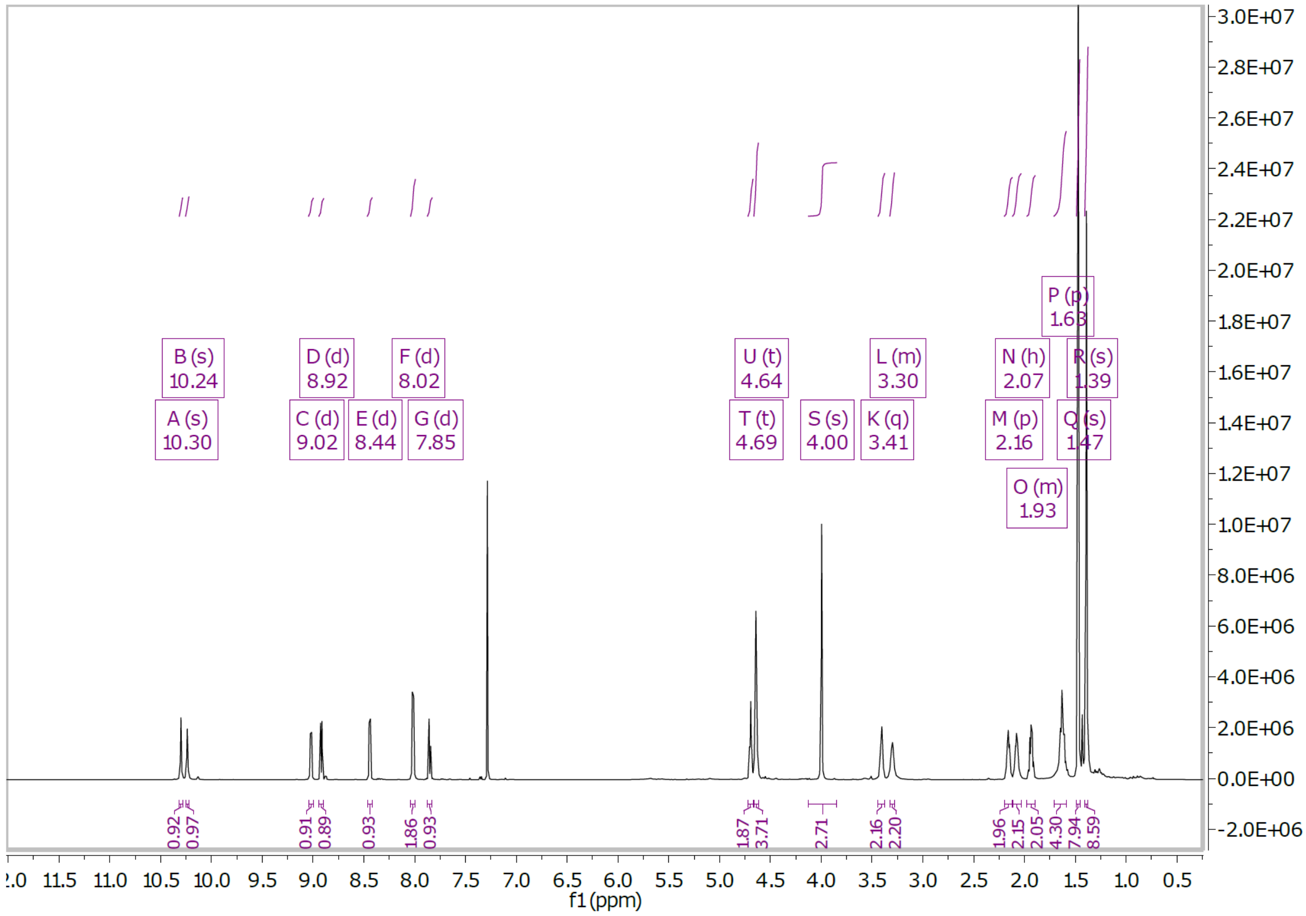


**Supplementary Figure 18.** $^{1}$H NMR of ADH-6-NCS.

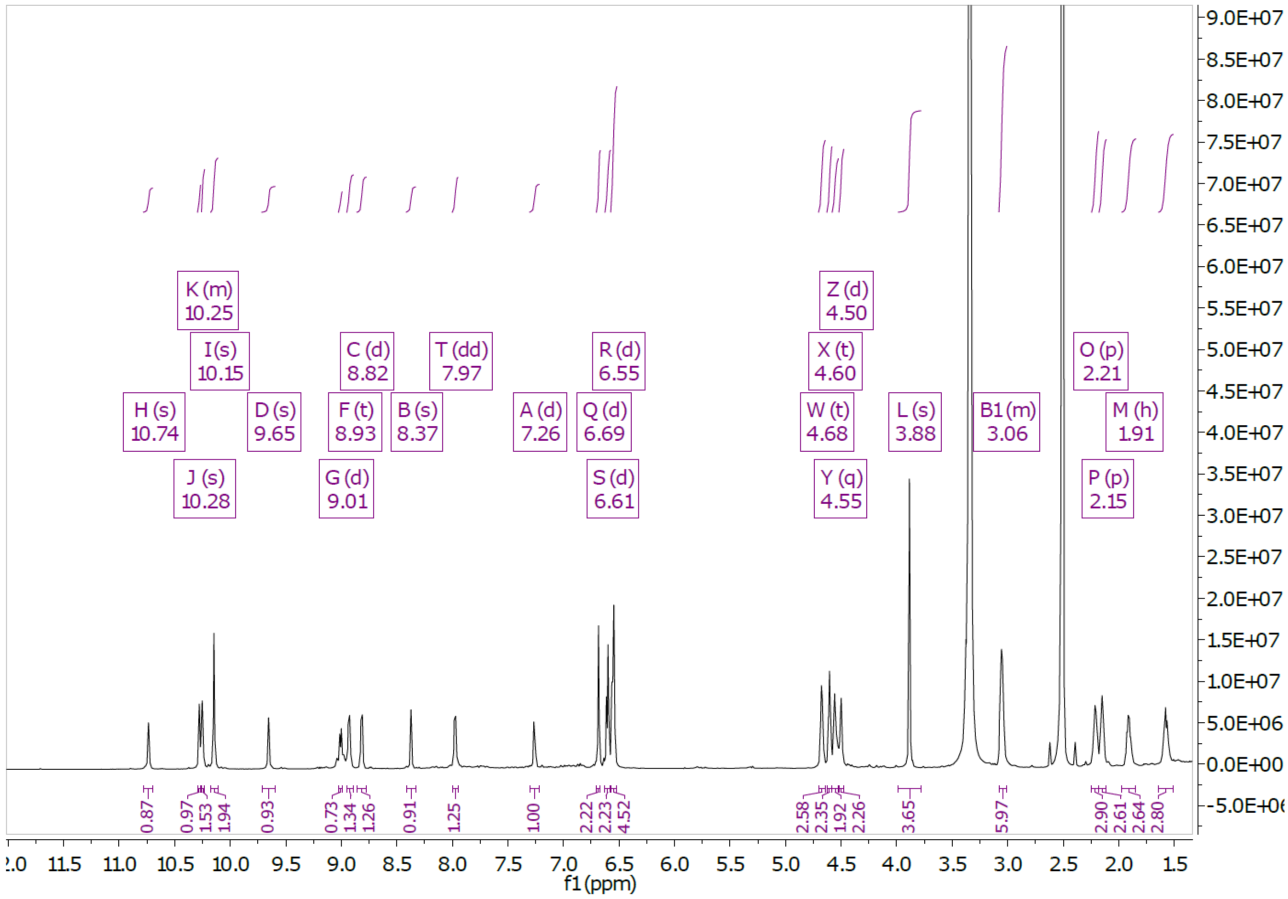


**Supplementary Figure 19.** $^{1}$H NMR of ADH-6$_{F}$ (ADH-6$_{FITC}$).
